# Amoebozoan testate amoebae illuminate the diversity of heterotrophs and the complexity of ecosystems throughout geological time

**DOI:** 10.1101/2023.11.08.566222

**Authors:** Alfredo L. Porfirio-Sousa, Alexander K. Tice, Luana Morais, Giulia M. Ribeiro, Quentin Blandenier, Kenneth Dumack, Yana Eglit, Nicholas W. Fry, Maria Beatriz Gomes E Souza, Tristan C. Henderson, Felicity Kleitz-Singleton, David Singer, Matthew W. Brown, Daniel J. G. Lahr

## Abstract

Heterotrophic protists are vital in Earth’s ecosystems, influencing carbon and nutrient cycles and occupying key positions in food webs as microbial predators. Fossils and molecular data suggest the emergence of predatory microeukaryotes and the transition to a eukaryote-rich marine environment by 800 million years ago (Ma). Neoproterozoic vase-shaped microfossils (VSMs) linked to Arcellinida testate amoebae represent the oldest evidence of heterotrophic microeukaryotes. This study explores the phylogenetic relationship and divergence times of modern Arcellinida and related taxa using a relaxed molecular clock approach. We estimate the origin of nodes leading to extant members of the Arcellinida Order to have happened during the latest Mesoproterozoic and Neoproterozoic (1054 - 661 Ma), while the divergence of extant infraorders postdates the Silurian. Our results demonstrate that at least one major heterotrophic eukaryote lineage originated during the Neoproterozoic. A putative radiation of eukaryotic groups (e.g. Arcellinida) during the early-Neoproterozoic sustained by favorable ecological and environmental conditions may have contributed to eukaryotic life endurance during the Cryogenian severe ice ages. Moreover, we infer that Arcellinida most likely already inhabited terrestrial habitats during the Neoproterozoic, coexisting with terrestrial Fungi and green algae, before land plant radiation. The most recent extant Arcellinida groups diverged during the Silurian Period, alongside other taxa within Fungi and flowering plants. These findings shed light on heterotrophic microeukaryotes’ evolutionary history and ecological significance in Earth’s ecosystems, using testate amoebae as a proxy.

**Significance Statement:** Arcellinida shelled amoebae are heterotrophic microbial eukaryotes with an extensive Neoproterozoic fossil record represented by the vase-shaped microfossils (VSMs), a diverse group that is abundant and widespread in late Tonian rocks (VSMs). Here we combined phylogenomic sampling and the fossil record to generate time-calibrated trees. Our results illuminate key events in the history of life, including: i) the Tonian origin of extant microbial eukaryote lineages; ii) a speculative proposed radiation of eukaryotes before the Cryogenian, “Tonian revolution”; iii) the establishment of complex terrestrial habitats before the Cryogenian; iv) a post-Silurian divergence of modern Arcellinida sub-clades in terrestrial (including freshwater) habitats. Our results provide valuable insights into the evolution of life throughout geological time and are congruent with recent discoveries regarding the early diversification of eukaryotes, including the Precambrian history of eukaryotic protosteroids.

## Introduction

Heterotrophic microbial eukaryotes play a crucial ecosystem role by contributing to the carbon and nutrient cycles (1, 2). These organisms, capable of phagocytosis, act as predators on bacterial and eukaryotic communities, playing a significant role in complex food webs supported by primary producers (1). Additionally, predation is an evolutionary innovation that likely contributed to the diversification of eukaryotes (3). The Last Eukaryotic Common Ancestor (LECA) was heterotrophic and capable of phagocytosis. However, the timing and specific conditions under which diverse lineages of heterotrophic microeukaryotes have proliferated in Earth’s ecosystems remain unclear (4–6).

Evidence from fossils, biomarkers, geochemical proxies, genomic data, and molecular clocks indicate that eukaryotes first originated during the Stenian (1200-1000 Ma) and Tonian periods (1000-720 Ma) (7–11). This led to a transition from a prokaryotic- to a eukaryotic-rich marine environment (6, 12–14), by 800 Ma, likely triggered by increased phosphorus, nitrate, and silica availability (14–17). From around this time, Neoproterozoic vase-shaped microfossils (VSMs) represent the remnants of an early eukaryotic divergence event. Organisms represented by VSMs are generally thought to have lived in marine environments, although a terrestrial habitat for these organisms is also plausible (18–20). The well-preserved nature of VSMs has allowed for detailed investigation and comparisons of their morphology to modern eukaryotic groups. These investigations support the current interpretation of a large fraction of VSMs being members of the stem or crown groups of testate amoeba order Arcellinida, due to both morphological affinities and congruence with molecular phylogenetic reconstructions (19, 21, 22). Other VSMs, such as *Melicerion poikilon*, had suggested affinities to Euglyphida, a distantly-related, convergent rhizarian clade of testate amoebae (18, 19). However, morphological evidence, in this case, is tentative, and the suggestion for a Euglyphida affinity is currently incongruent with molecular phylogenetic reconstructions, as Euglyphida is part of a clade of cercozoan filose amoebae, which appears to be much younger, around 292 Ma (23). Arcellinida is a diverse lineage of extant heterotrophic microeukaryotes within the Amoebozoa, found in terrestrial and freshwater environments (19–22). Since many VSMs have been recognized as Arcellinida, they are accepted to represent the oldest and most diverse fossil evidence of heterotrophic microeukaryotes (19–20, 24,25). Elucidating the origin and evolutionary history of Arcellinida (and derived lineages) in light of their microfossil record is pivotal to illuminate the early evolution and possible radiation of heterotrophic microbial eukaryotes, and serve as a proxy to infer the complexity of Earth’s ecosystems over geological time (19–20, 25, 26).

Recent efforts of sampling diverse amoebozoan testate amoebae in a phylogenomic framework have resolved their deep phylogenetic relationships (21, 27). Amoebozoa is home to at least two testate amoebae groups: Arcellinida and Corycidia. Arcellinida is a diverse order represented by lineages that build hard extracellular shells, with the potential to generate exceptionally preserved fossilized remains (19, 21). Corycidia is a recently established subclade of Amoebozoa represented by the lineages of testate amoebae that produce flexible shells and do not branch within Arcellinida (27). Despite recent advances, many lineages still remain unsampled (21).

In addition to expanding the diversity of sampled Arcellinida at the genome level, the potential of a highly resolved phylogeny to provide insight into timing the Arcellinida origin and divergence of sub-clades has not been explored (22, 25). VSMs can be interpreted either as stem or crown Arcellinida. In either case, these fossils can be used to calibrate a phylogenetic tree and estimate the divergence time of lineages both within Arcellinida, as well as closely related amoebozoans. These diverse VSMs found in sedimentary deposits around the world have been continuously investigated, and their stratigraphy refined over time, enabling us to constrain these fossils’ ages (19–20, 28–36). In this context, combining the VSMs and phylogenetic tree calibration opens up avenues to time the evolution of Arcellinida and closely related groups.

Here, we investigate the origin and divergence times of Arcellinida and closely related amoebozoan taxa using phylogenomics and a relaxed molecular clock approach. We expanded the taxonomic sampling for amoebozoan testate amoebae, including 14 taxa that lacked precise placement, to produce a new phylogenomic dataset (utilizing 226 genes). We considered the diverse record of VSMs and Metazoa fossils to time calibrate this well-resolved deep phylogenomic tree. Different calibration strategies and molecular clock models support the divergence of extant Arcellinida lineages during the latest Mesoproterozoic and early to mid-Neoproterozoic, between 1060 and 661 Ma. We thus corroborate the origin of a major eukaryotic group by the Neoproterozoic, including a recorded establishment of heterotrophy predating the Cryogenian Period. Overall, using amoebozoan testate amoebae as a proxy, we provide insights into the evolution of microbial eukaryotes and Earth’s early ecosystems.

## Results

### A resolved tree of amoebozoan testate amoebae

We constructed a concatenated supermatrix using 57 taxa and 226 genes (70,428 amino acid sites) using the PhyloFisher v. 1.2.11 package (37, 38). We include novel data for 14 testate amoebae taxa, obtained through single-cell or whole-culture transcriptomics (**Fig. 1; Dataset S01, Tables S1 - S4**). The remaining testate amoebae and sister-group taxa were sampled based on previously available genomes and transcriptomes (**Dataset S01, Table S1**). The resulting phylogenomic tree recovers a monophyletic Arcellinida, with three well-defined suborders (Phryganellina, Organoconcha, and Glutinoconcha) and five infraorders within Glutinoconcha (**Fig. 2; Appendix S01, Figs. S1 and S2**). The Corycidia clade is also recovered with full support, with two families, Trichosidae and Amphizonelliidae fam. nov. (**Fig. 2**). Nearly all nodes of the tree are fully (=100%) or highly (>92%) supported by Maximum Likelihood nonparametric Real Bootstraps (MLRB), except for a single node within Sphaerothecina clade that has lower support (MLRB = 76%; **Fig. 2**). We have also produced single-gene reconstructions using SSU rDNA and Cytochrome Oxidase subunit I (COI) **(Dataset S01, Tables S5-S6),** which are the genes traditionally used to reconstruct relationships in Arcellinida. These analyses present broader taxon sampling but failed to recover most of the deeper relationships in Arcellinida **(Appendix S01, Figs. S3-S4**)

**Figure 1.**
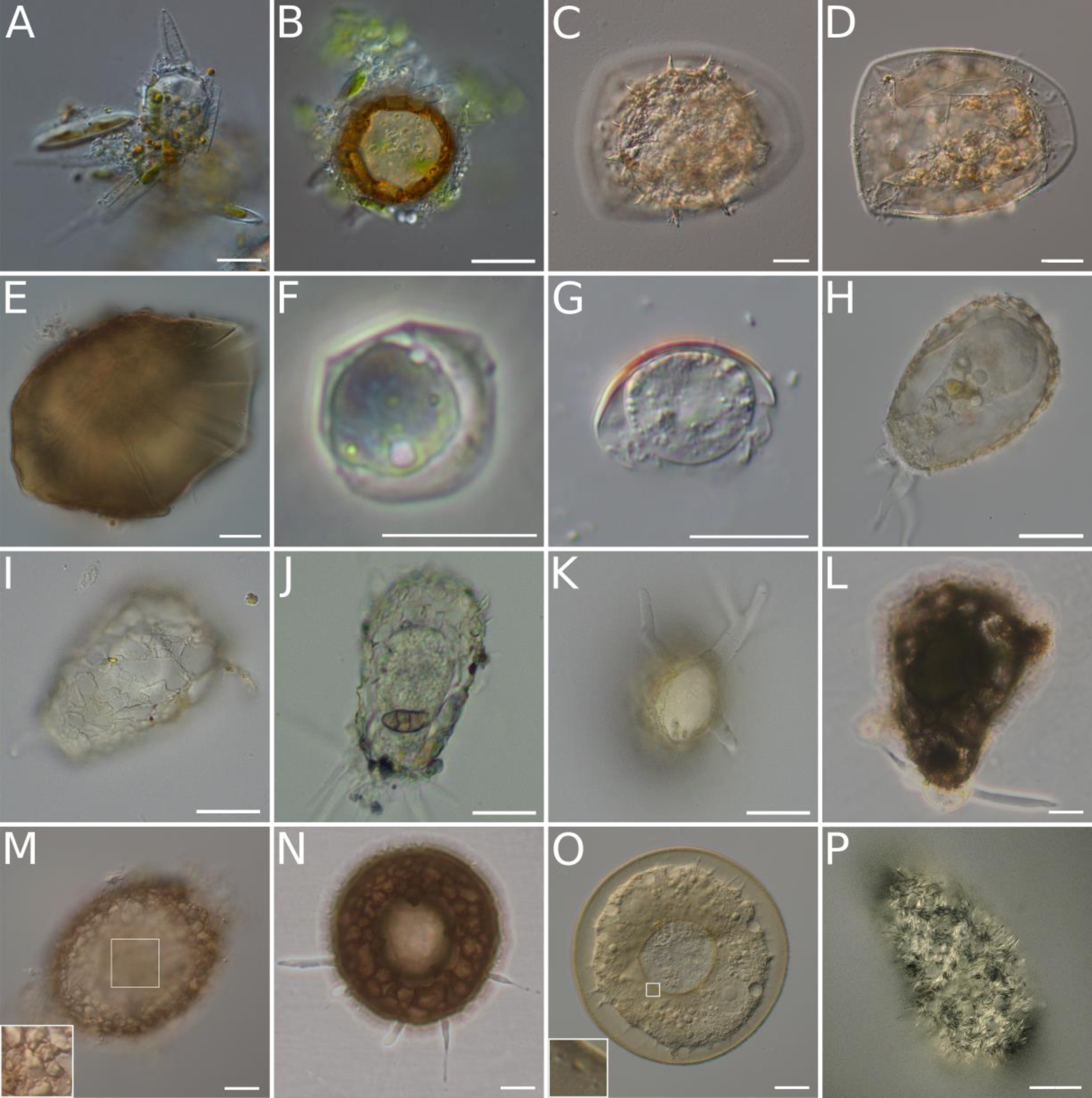
Sampled testate amoebae — Arcellinida and Corycidia. The pictured organisms were photodocumented prior to molecular processing and represent individuals or cultures from which the transcriptomic data was obtained. The scale bars represent 20 µm, except when specified. **A.** *Phryganella paradoxa* T, lateral view. **B.** *Phryganella acropodia* A, apertural view. **C - D.** *Microcorycia aculeata*, dorsal view (C) and apertural view (D); **E.** *Microcorycia flava*, dorsal view focusing on the flexible part of the shell; **F.** *Spumochlamys* sp., dorsal view; **G.** *Spumochlamys bryora*, lateral view; **H - K.** *Heleopera lucida* comb. nov. (previously *Difflugia lucida*), lateral view focusing on the cell within the shell (H), lateral view focusing on the shell (I - J), and apertural view focusing on the compressed aspect of the shell (K); **L.** *Difflugia* cf. *capreolata*, lateral view, scale bar 40 µm; **M.** *Netzelia lobostoma*, lateral view, white square focusing on details of the shell; **N.** *Cyclopyxis* sp., apertural view, scale bar 40 µm; **O.** *Galeripora* sp., apertural view, white square focusing on the pores which surround the shell aperture; **P.** *Trichosphaerium sp.* KSS. Measured morphometric characteristics of the newly sequenced testate amoebae taxa are present on **Dataset S01, Table S2**.

**Figure 2.**
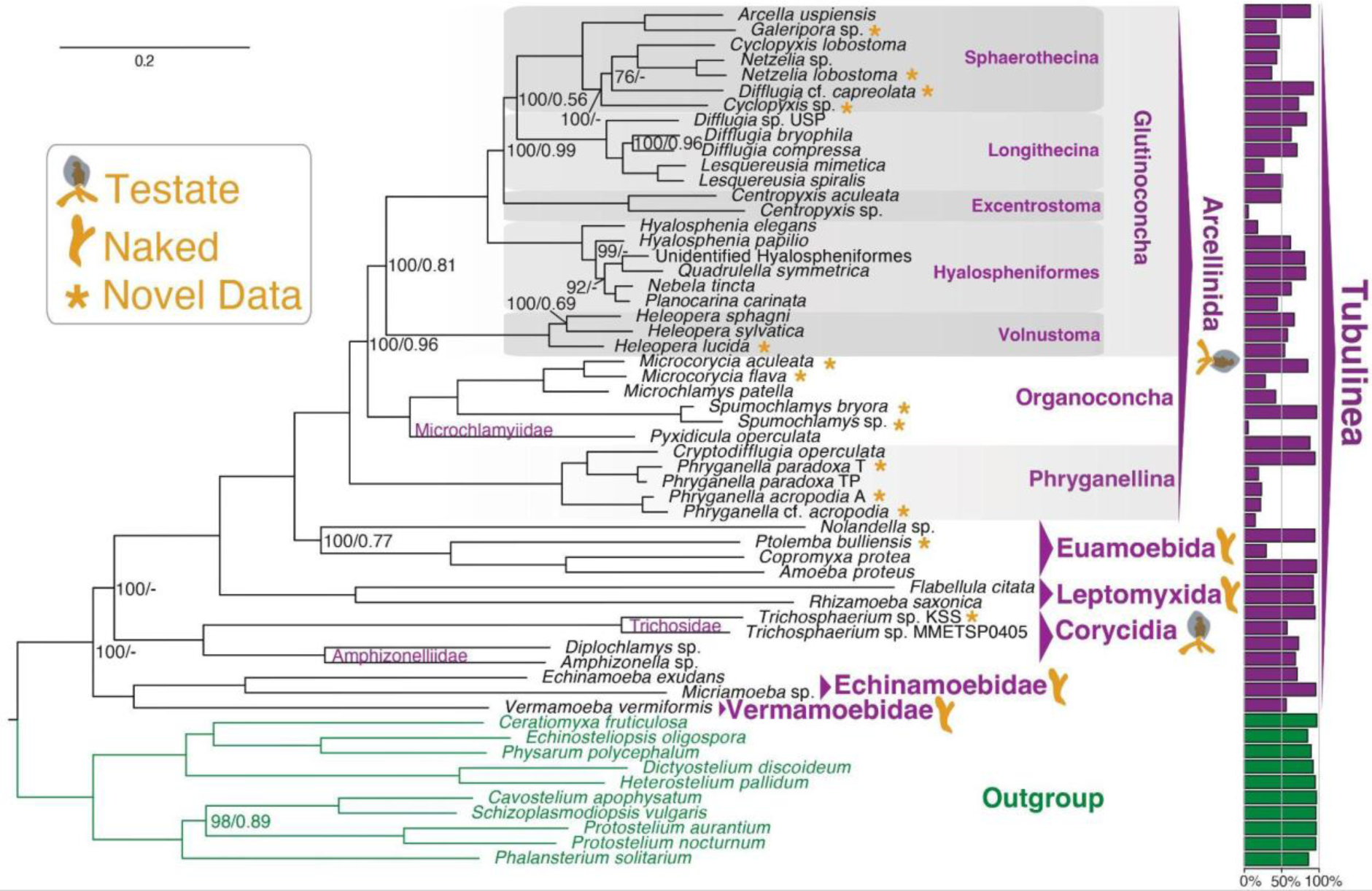
The tree of amoebozoan testate amoebae. 226 genes (70,428 amino acid sites) phylogeny of amoebozoan testate amoebae rooted with Evosea (Amoebozoa). The tree was initially built using IQ-TREE2 v. 2.0-rc1 under the LG+C20+G4 model of protein evolution and further used to infer a Posterior Means Site Frequency model using the ML model LG+C60+G4+PMSF. Topological support was assessed by 100 Maximum Likelihood Real Bootstrap Replicates (MLRB) and local posterior probability values (LPP) calculated using ASTRAL-III v. 5.7.3, and are shown in the format (MLRB/LPP).

### Time-calibrated tree of amoebozoan testate amoebae

For estimating divergence times in testate amoebae evolution and closely related taxa, we expanded our phylogenomic supermatrix to consider a representative sampling of Amorphea, including Amoebozoa, Fungi, Metazoa, and their protistan relatives **(Dataset S01, Table S7**). For a comprehensive approach, taking into account the alternative interpretations of the VSM record, we implemented four different calibration strategies: i. calibration of nodes within Metazoa, excluding the VSM record to calibrate amoebozoan nodes; ii. calibration of nodes within Metazoa and calibration of Glutinoconcha+Organoconcha and Glutinoconcha nodes, considering VSMs as derived crown Arcellinida; iii. calibration of nodes within Metazoa and calibration of the Arcellinida node, considering VSMs as basal crown Arcellinida; iv. calibration of nodes within Metazoa and calibration of Arcellinida+Euamoebida node, considering VSMs as stem Arcellinida (**Dataset S01, Table S8 - S10**). To implement these four fossil calibration strategies, we performed a total of 36 experiments considering three different distributions (i.e., Uniform, Skew-Normal, or Truncated-Cauchy short-tail) under an uncorrelated or autocorrelated relaxed clock model with either a drift parameter of α = 2 and β = 2 or α = 1 and β = 10. We ran each experiment in two independent MCMC chains to check for convergence, which was achieved for all analyses (**Appendix S01, Fig. S5**).

Comparing all the calibration strategies and experiments, we observed overall similar inferred times with the uncorrelated clock model (median = 591 - 531 Ma and mean = 627 - 549 Ma; **Dataset S01, Table S11 and Appendix S01, Figs. S6 - S43**) and the autocorrelated clock model (median = 592 - 531 Ma and mean = 671 - 557 Ma; **Dataset S01, Table S11**). Regarding implemented distributions, overall the uniform distribution inferred the youngest ages, while Truncated-Cauchy inferred slightly older ages (**Dataset S01, Table S11**). The drift parameter (i.e., *α* = 2 and β = 2 vs. α = 1 and β = 10) had virtually no impact independent of the clock model, distribution, and calibration strategy (**Dataset S01, Table S11**). The estimated times for the Arcellinida node by excluding VSMs from the calibration (mean = 911 - 734 Ma; 95% highest probability density confidence interval (HPD CI) = 1060 - 605) and by including VSMs in the calibration (mean = 930 - 746 Ma; 95% HPD CI = 1054 - 661 Ma) are highly congruent. This agrees with the current interpretation that VSMs represent fossil remains of Arcellinida, supporting their use to calibrate amoebozoan nodes. Aiming for a comprehensive approach, the time estimation results shown and discussed hereafter focus on the full range of times estimated based on the calibration strategies that included the VSMs, the three distributions, and the drift parameter of α = 2 and β = 2 under the uncorrelated or autocorrelated relaxed clock model.

The molecular clock analyses inferred a mean time for the root of Amorphea to be between 1640 and 1393 Ma (95% HPD CI = 1843 - 1088 Ma; **Fig. 3, Dataset S01, Table S11, and Appendix S01, Fig. S7 - S43**). For Metazoa, the mean time estimated ranged between 835 and 734 Ma (95% HPD CI = 872 - 673 Ma; **Fig. 3, Dataset S01, Table S11, and Appendix S01, Fig. S7 - S43**). The mean estimated for the origin of Amoebozoa ranged from 1607 - 1298 Ma (95% HPD CI = 1795 - 1045 Ma; **Fig. 3, Dataset S01, Table S11, and Appendix S01, Fig. S7 - S43**). The Arcellinida node is constrained within the mean 930 and 746 Ma (95% HPD CI = 1054 - 661 Ma; **Fig. 3, Dataset S01, Table S11, and Appendix S01, Fig. S7 - S43**), estimating an origin for Arcellinida during the latest Mesoproterozoic and Neoproterozoic (**Fig. 4A**). For the early divergence time of Arcellinida subclades, the estimated times suggest that the split between the Organoconcha and Glutinoconcha branches occurred during the Neoproterozoic (mean = 855 - 679 Ma; 95% HPD CI = 969 - 600 Ma; **Figs. 3-4B**). Regarding the suborders of Arcellinida, we inferred a mean origin for Phryganellida between 534 - 265 Ma (95% HPD CI = 661 - 175 Ma; **Figs. 3-4C**), for Organoconcha 735 - 550 Ma (95% HPD CI = 839 - 463 Ma; **Figs. 3-4D**), and for Glutinoconcha between 790 - 621 Ma (95% HPD CI = 897 - 539 Ma; **Figs. 3-4E**). For the deeper nodes of Glutinoconcha, the most sampled suborder of Arcellinida, the estimated divergence times ranged between Cryogenian and Carboniferous (mean = 643 - 393 Ma; 95% HPD CI = 705 - 335; **Figs. 3-4F-H**). The inferred ages for Glutinoconcha infraorders are relatively more widespread depending on the calibration strategy when compared to other nodes. Treating VSMs as derived crown taxa constrains the infraorders origin between Ediacaran and early Cretaceous (mean = 422 - 221 Ma; 95% HPD CI = 575 - 123 Ma; **Figs. 3-4I-M**) while considering VSMs as basal crown or stem Arcellinida estimate their origin mostly between Silurian and early Cretaceous (mean = 339 - 172 Ma; 95% HPD CI = 444 - 122 Ma; **Figs. 3-4I-M**). It is worth noting that only the calibration using a uniform autocorrelated clock model and VSMs as derived crown arcellinids inferred ages as old as the Ediacaran for the Glutinoconcha infraorders. All other distribution-clock models consistently led to ages constrained within the Paleozoic. Most inferred ages for the nodes representing the origin of the modern extant genera and species of Arcellinida postdate the Silurian (mean = 416 - 125 Ma; **Dataset S01, Table S11**). Besides the testate amoebae, the inferred mean times for the other orders and major groups of Amoebozoa we sampled ranged from 585 Ma to 1288 Ma, placing their origin mostly during the Neoproterozoic (**Dataset S01, Table S11**). The results and time-calibrated trees for all experiments are present in the supplemental material (**Dataset S01, Table S11 and Appendix S01, Fig. S6 - S43**).

**Figure 3.**
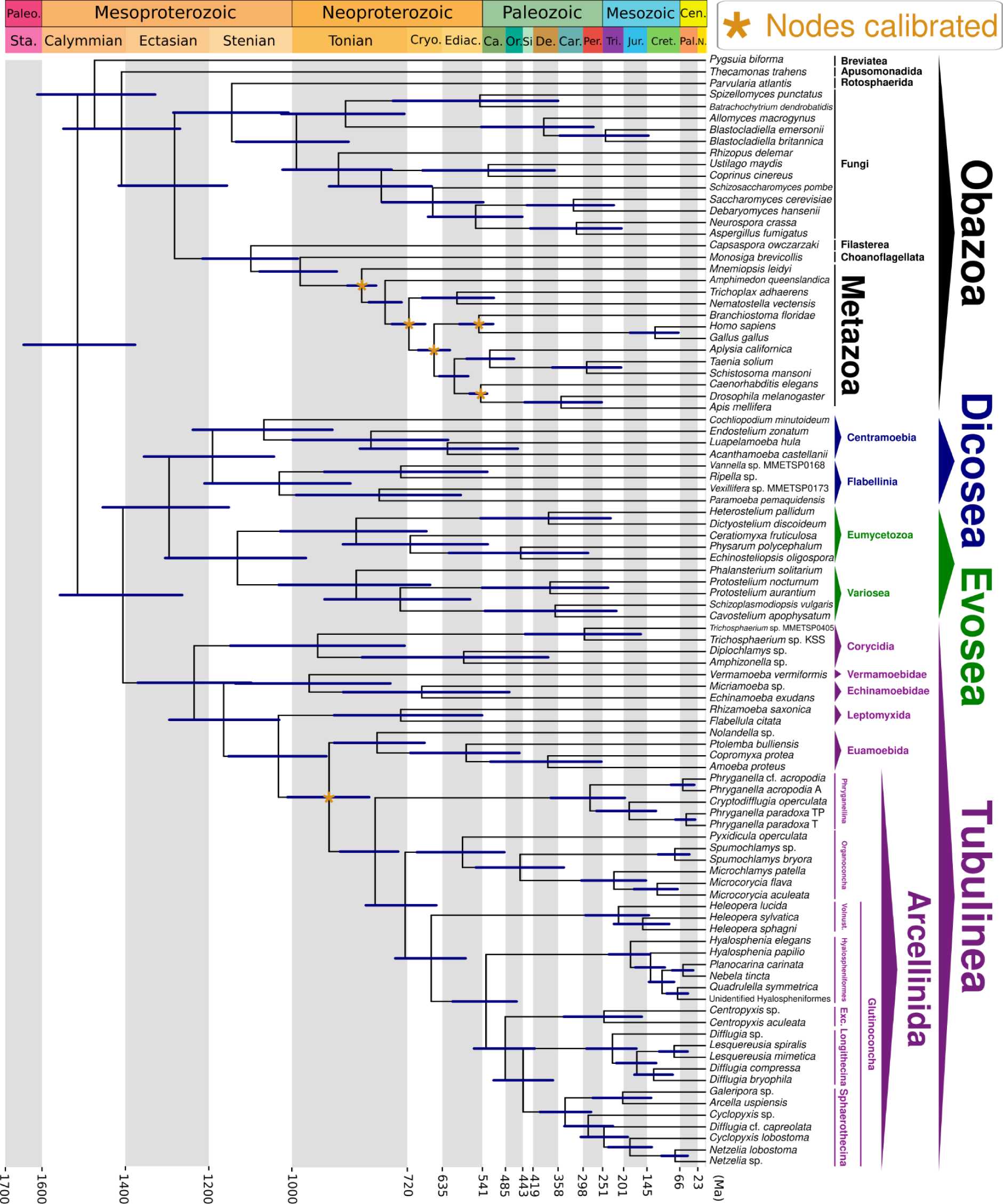
Amorphea time-calibrated tree inferred under autocorrelated relaxed clock model, applying a truncated-Cauchy distribution for node calibration and drift parameter of α = 2 and β = 2, considering VSMs as stem Arcellinida to calibrate Arcellinida+Euamoebida node. Bars at nodes are 95% highest probability density confidence intervals (HPD CI). Asterisks indicate the nodes calibrated based on external fossil information. Out of the 36 time-calibrated trees generated, we display here the one representing the analysis that estimated the youngest minimum 95% HPD CI for the two earliest nodes of Arcellinida, thus showing the youngest age estimated for the origin of nodes leading to extant members of the Arcellinida Order. The results and time-calibrated trees for all experiments are present in the supplemental material (**Dataset S01, Table S11 and Appendix S01, Fig. S6 - S43**). Abbreviations: Exc.- Excentrotoma; Vonust. - Volnustoma; Paleo. - Paleoproterozoic; Sta. - Statherian; Cryo. - Cryogenian; Ediac. - Ediacaran; Ca. - Cambrian; Or. Ordovician; Si - Silurian; De. - Devonian; Car. Carboniferous; Per. - Permian; Tri. - Triassic; Jur. - Jurassic; Cret. - Cretaceous; Cen. - Cenozoic; Pal. - Paleogene; N. - Neogene; My - Million Years.

**Figure 4.**
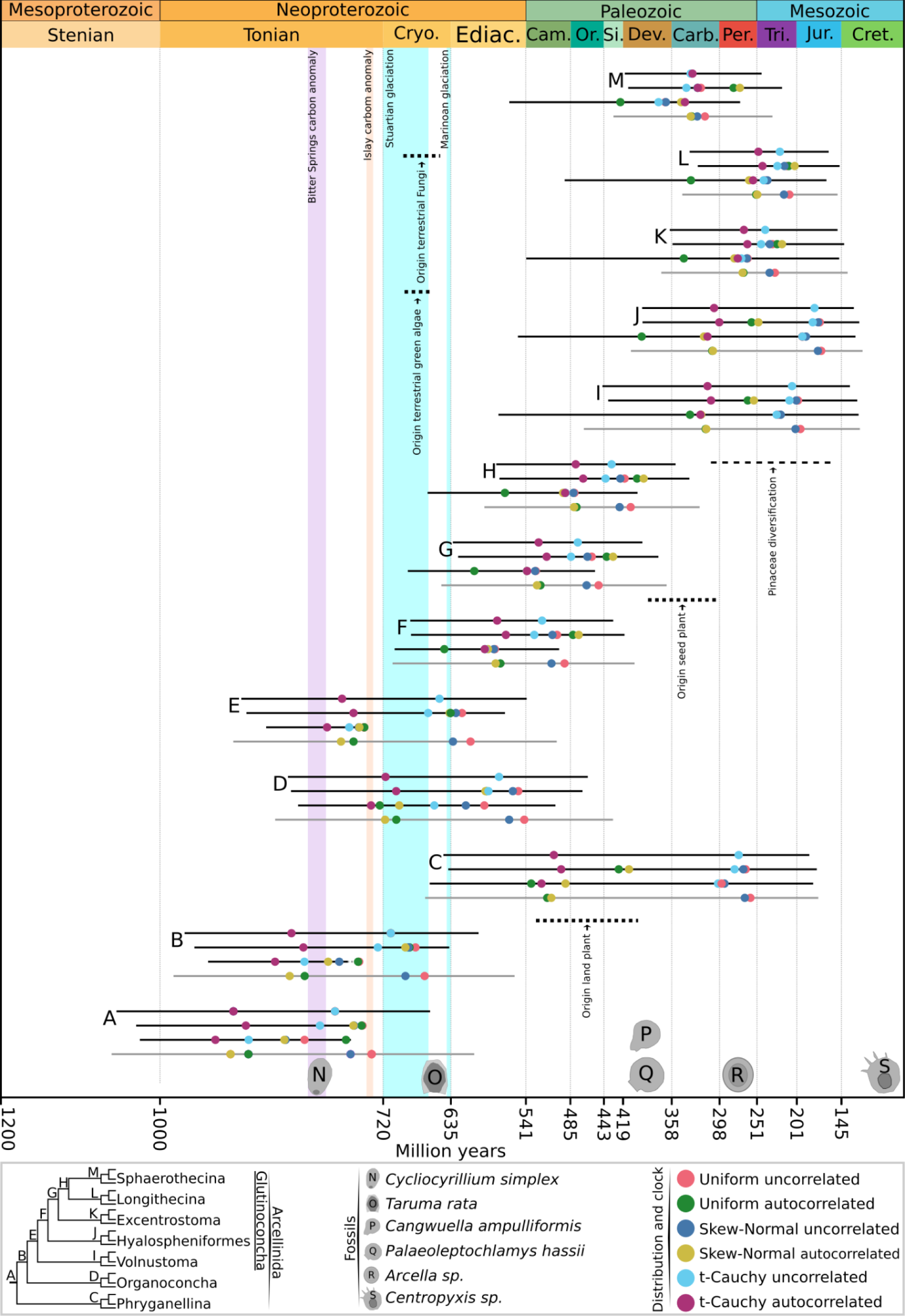
Estimated times for Arcellinida nodes throughout geological time. The figure highlights key carbon anomalies, ice age events, the Arcellinida fossil record (19, 26, 36, 44, 72), and the divergence time of other groups of organisms suggested by previous studies (horizontal dotted lines; 75). Displayed for each node are four bars representing, from bottom to top, the calibration strategy not considering VSM record to calibrate amoebozoan nodes (gray bar), the calibration strategy considering VSMs as derived crown Arcellinida, calibration strategy considering VSMs as basal crown Arcellinida, and calibration strategy considering VSMs as stem Arcellinida. The bars represent the combination of all 95% highest probability density confidence intervals estimated by each distribution-clock model considered and the colored dots represent the mean estimated time by each distribution-clock model. The results and time-calibrated trees for all experiments are present in the supplemental material (**Dataset S01, Table S11 and Appendix S01, Fig. S6 - S43**).

### Ancestral habitat reconstruction of Arcellinida

We performed statistical analyses on the ancestral habitat of key hypothetical ancestors within Arcellinida considering alternative scenarios of a terrestrial or marine origin for the crown group (**Dataset S01, Table S12 and Appendix S01, Fig. S44**). The unrestrained reconstruction inferred a terrestrial habitat (100% probability) for all nodes within Arcellinida. The ancestral reconstruction that sets the fixed value of a marine state on the last common ancestor of modern Arcellinida inferred a high probability of terrestrial habitat for all nodes within Arcellinida (>93%), implying at least two independent transition events (2TE) from marine to terrestrial habitats (**Fig. 5**). The ancestral reconstruction that sets the fixed value of a marine state on the last common ancestor of both the Arcellinida and the Organoconcha+Glutinoconcha clades inferred with high probability (>88%) a terrestrial habitat for the hypothetical ancestors of Phryganellina, Organoconcha, and Glutinoconcha, implying at least three independent transition events (3TE) from marine to terrestrial habitats (**Fig. 5**).

**Figure 5.**
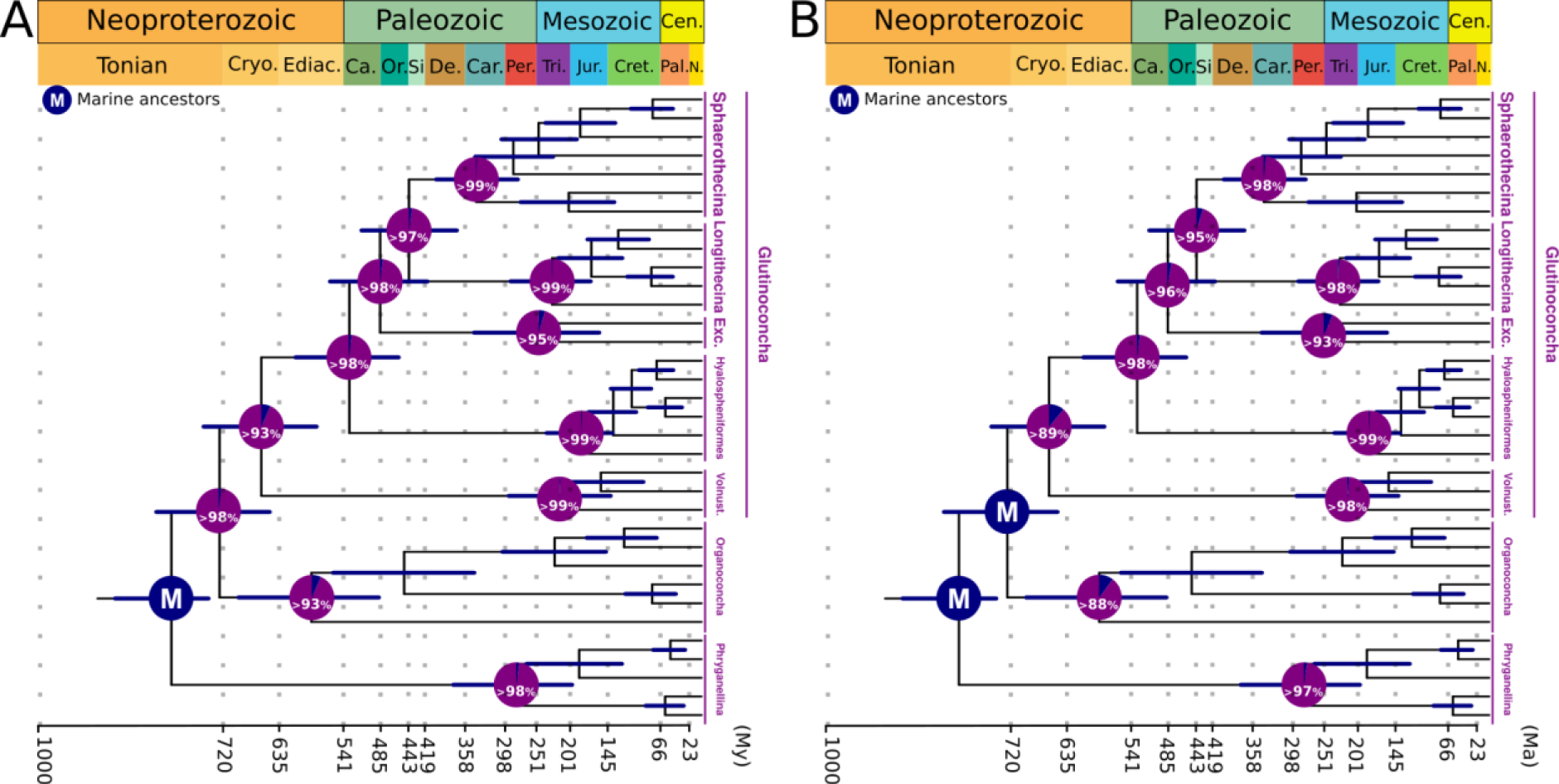
Ancestral reconstructions of Arcellinida habitats using BayesTraits. Pie charts at each node indicate the mean probabilities of a hypothetical marine ancestor (blue) or a hypothetical terrestrial ancestor (purple). **A.** Reconstruction considering Arcellinida ancestor as marine, implying at least two independent transition events (2TE). **B.** Reconstruction considering Arcellinida and Organoconcha+Glutinoconcha ancestors as marine, implying at least three independent transition events (3TE). Bars at nodes are 95% highest probability density confidence intervals estimated by the calibration strategy using an autocorrelated relaxed clock model, applying a truncated-Cauchy distribution for node calibration and drift parameter of α = 2 and β = 2, considering VSMs as stem Arcellinida to calibrate Arcellinida+Euamoebida node. The complete results are shown on **Dataset S01, Table S12 and Appendix S01, Fig. S44**.

## Discussion

### A resolved tree of amoebozoan testate amoebae

The phylogenomic dataset constructed in this study improves several aspects of the previously available amoebozoan testate amoebae dataset (21, 27). Through the PhyloFisher pipeline, we were able to construct a curated phylogenomic matrix that is free of paralogs and contamination (**Appendix S01, SI1**). Moreover, the matrix constructed is an accessible and easy-to-update dataset, since newly sequenced transcriptomes can be easily added through PhyloFisher to further expand the taxonomic sampling of amoebozoan testate amoebae in a phylogenomic approach. In general terms, the phylogenomic tree obtained here is consistent with the previously published phylogenomic tree for amoebozoan testate amoebae (**Fig. 2**; 21, 27). By recovering all major groups of the Arcellinida and Corycidia clades with full support, in accordance with previous reconstructions, we corroborate the robustness of the backbone of the amoebozoan testate amoebae tree. The phylogenomic analyses enable us to place previously unsampled taxa within Arcellinida suborders, supporting the taxonomic actions regarding *Heleopera lucida* comb. nov. and the genus *Microcorycia* (**Appendix S01, SI2 - SI3**). Moreover, the phylogenomic tree identifies taxa that will need future taxonomic revision based on further morphological and molecular studies, those being *Difflugia* cf. *capreolata* and the genera *Cyclopyxis* and *Phryganella*. Detailed discussion on the placement of newly sequenced amoebozoan testate amoebae is presented in **Appendix S01, SI2 - SI3**.

### Robust Amorphea time-calibrated trees using VSMs

Our time-calibrated trees are congruent with several molecular clocks that have sampled the diversity of eukaryotes (**Fig. 3, Dataset S01, Table S11, and Appendix S01, Fig. S6 - S43**). The estimated ages for the root of our tree fall within the range inferred by a recent molecular timescale for eukaryotes (95% confidence interval = 2177 - 1356 Ma for Amorphea; 39). Similarly, the estimated ages for the origin of other major clades align with previous studies, including Obazoa (95% HPD CI = 2305 - 1526 Ma), opisthokonts (95% HPD CI = 2019 - 1051 Ma), and animals (95% HPD CI = 833 - 680 Ma) (39–42). The inferred times place the origin of Arcellinida during the latest Mesoproterozoic and early to mid-Neoproterozoic, most likely during the Tonian Period and no later than the Cryogenian (**Fig. 4, Dataset S01, Table S11, and Appendix S01, Fig. S6 - S43**). It is worth noting that by considering different calibration strategies to account for alternative plausible interpretations of VSMs and the unavoidable fossil record uncertainties, the times we estimated for each node have wide ranges. However, independently of the calibration strategy, distribution, and clock model, the Arcellinida origin is mostly placed within the Neoproterozoic, and all estimated mean times suggest a Tonian origin. Notably, the time-calibrated trees we generated without using the VSMs as calibration data inferred times congruent with our analyses considering the Tonian VSMs. This demonstrates that inferred ages are not a fossil calibration bias and serves as further corroboration for the interpretation of VSMs as fossil members of, at the very least, the stem Arcellinida group (**Fig. 4, Dataset S01, Table S11**). Also, the ages we inferred for the origin of modern groups (i.e., genera) are highly congruent with the recent Arcellinida fossil record which postdates the Carboniferous Period and preserves fossils assigned to genera like *Arcella*, *Difflugia*, and *Centropyxis* (**Dataset S01, Table S11;** 43-45). Collectively, the consistency of the estimated times, and their congruence with previous molecular clocks and with the fossil record, support that including VSMs to calibrate a comprehensive phylogenomic sampling of Amorphea leads to robust results.

### The Neoproterozoic diversification of heterotrophic microbial eukaryotes

Our time-calibrated trees reveal that Arcellinida originated most likely during the Neoproterozoic, with most inferred times and all of the means falling within the Tonian Period (**Fig. 4A - B**). The mean estimated times for other major groups of Amoebozoa also indicate a Tonian origin. Similarly, previous molecular clock analyses have indicated the divergence of multiple heterotrophic eukaryotes during this period (40, 41). However, their diversity has been challenging to examine due to the lack of a fossil record for these organisms. Nevertheless, the estimated Neoproterozoic origin for the Arcellinida crown group and the diversity of Tonian VSMs, currently represented by 14 morphologically diverse genera (19–20, 25, 31), suggest that by the Tonian Period, Earth’s ecosystems had witnessed the origin of modern heterotrophic microbial eukaryotes.

An inferred Tonian origin for diverse heterotrophic microbial eukaryotes is congruent with proposals of ecosystem establishment during the Neoproterozoic, stemming from various disciplines. Previous studies speculate the existence of a “Tonian revolution”, based on evidence from biomarkers, fossil records, and molecular data that infer a marked transition from prokaryotic- to eukaryotic-rich ecosystems by 800 Ma (6, 11–14, 46). This transition was likely facilitated by factors such as increased availability of nitrate, phosphorus, silica, and reduced toxicity, which provided a favorable condition linked to the documented diversification of eukaryotic phototrophs at that time (14–17). In turn, the establishment of a community of phototrophs may have served as a favorable condition for the diversification of heterotrophs. Fossil and geochemical evidence suggest that photosynthetic biological mats contributed to the establishment of Oxygen Oases during the Tonian, likely triggered by a higher capacity of oxygen productivity of eukaryotic phototrophs (7, 10, 47, 48). These oases probably represented an increase in aerobic conditions, and food availability, that were permissive to the survival and proliferation of heterotrophs like Arcellinida. Consequently, a combined interpretation of the geochemical and fossil records indicates that complex ecosystems were established by the Neoproterozoic.

VSMs serve as a unique testimony of the putative “Tonian revolution” on eukaryotic diversification and ecosystems established during the Neoproterozoic. The VSMs have been found in rocks characterized by organic-rich sediments corroborating the close association of heterotrophic eukaryotes with microbial mats (19). Also, the organisms represented by these VSMs likely preyed on both the bacterial and eukaryotic communities, similar to extant Arcellinida (19, 49, 50). Culture observations have demonstrated diverse strategies of extant Arcellinida to prey on various organisms, including diatoms, fungi, and nematodes (51–54). Moreover, several VSMs exhibit holes on their shells interpreted as predation marks, suggesting they also served as prey to other heterotrophs (3, 55). Predation has been interpreted as one of the triggers for eukaryotic diversification, including for animals, and VSMs are among the oldest records of this evolutionary innovation (3, 56–58). Thus, while most of the microbial eukaryotes did not leave fossils, VSMs stand out as a robust fossil record highlighting the rise of predation and increase of food web complexity on Earth’s ecosystems no later than the Tonian Period.

Our time-calibrated trees suggest that the divergence of some modern Arcellinida lineages happened during the Neoproterozoic. The inferred times for the split and origin of the Organoconcha and Glutinoconcha suborders mostly fall within the Neoproterozoic between the Tonian and Ediacaran Periods, with only some estimated times suggesting an early-Paleozoic origin for Organoconcha. Specifically, the ages predicted for the Organoconcha and Glutinoconcha split either predate or overlap with the Cryogenian Period and its glaciations. It is worth noting that Phryganellina is currently the least genomically sampled Arcellinida group, represented only by two genera, thus it is difficult to assess when this suborder originated. Altogether, the origin and early divergence of the Arcellinida crown group, and the diverse VSMs record, imply the capability of Neoproterozoic ecosystems not only to sustain heterotrophic eukaryote groups but also to allow their diversification.

### A possible Pre-Cryogenian Eukaryotic Diversification and Survival during Earth’s most severe Ice Ages

The Sturtian and Marinoan glaciation periods witnessed extensive ice coverage across the planet, with glaciers extending into tropical regions (59–61). The survival of life during Cryogenian glaciations can be explained by the presence of refugia and the adoption of dormancy strategies (62–69). The presence of cyst-like structures identified inside VSMs serves as direct evidence that these organisms were already capable of entering dormancy stages, similar to extant Arcellinida (70). Additionally, recent discoveries have revealed that Tonian VSM taxa persisted into the Cryogenian Period, providing fossil records that showcase the diversity of heterotrophic eukaryotes during glaciation periods (69). The inferred times for the split and origin of Glutinoconcha and Organoconcha suggest they possibly originated during Cryogenian, indicating not only survival but also divergence of novel modern eukaryotic taxa during Cryogenian glaciations (**Fig. 4D-E**). Consequently, the VSM record and the timing of early Arcellinida evolution enable speculation about a possible radiation of heterotrophic eukaryotic life shortly before and during the Cryogenian Period. This, coupled with a capacity for dormancy and the exploitation of habitat refugia, may explain the endurance of life during Earth’s most severe ice age.

### Timing of terrestrial conquest by Arcellinida

Currently, while it is largely suggested that the organisms represented by Tonian VSMs lived in shallow marine environments, a terrestrial habitat cannot be ruled out. To date, VSMs have been reported from Tonian sedimentary deposits described as fully or partially marine (18–19, 25). However, although scarcely discussed in the literature, it is plausible to hypothesize that the organisms represented by the VSMs may have lived in terrestrial environments and were deposited in marine sediments through a number of possible mechanisms, including: surface runoff, river discharge, wind blowing, or supratidal spillover. In any case, these organisms ultimately fossilized in a marine setting (71). Consequently, considering the alternative interpretations of VSMs’ natural habitat and their affinity to Arcellinida (i.e., stem or crown Arcellinida), different scenarios can be explored regarding Arcellinida’s conquest of terrestrial habitat.

Our ancestral habitat reconstructions indicate three alternative scenarios, a terrestrial origin for Arcellinida, a marine origin with two independent transition events (2TE) from marine to terrestrial habitats, and a marine origin with three independent transition events (3TE) (**Fig. 5 and Appendix S01, Fig. S44**). Although the reconstruction of a terrestrial origin is statistically superior to the other scenarios (Likelihood Ratio Test - LRT), this was already expected since all extant Arcellinida are terrestrial/freshwater inhabiting, leading to the reconstruction of a terrestrial ancestor (i.e. a possible systemic bias). However, interpreting VSMs as stem or crown Arcellinida and enforcing a marine origin for the Arcellinida stem and early crown groups, 2TE or 3TE are plausible and statistically equivalent based on LRT, in accordance with previous hypotheses (18–21, 25, 72). Multiple transition events are biologically plausible: Arcellinida-related amoebozoan lineages are often represented by both marine and terrestrial species, even within the same genera (e.g., *Vannella*, *Mayorella*, and *Trichamoeba*).

Coupling the reconstructed scenarios with the timing of Arcellinida origin and early divergence of its sub-clades, we infer that many Arcellinida probably inhabited terrestrial environments already in the Neoproterozoic, no later than the Ediacaran Period. Even if we consider the latest transition scenario reconstructed (3TE) the inferred times place the terrestrialization event of Glutinoconcha and Organoconcha most likely between the Tonian and Ediacaran periods (**Fig. 5**). The time of diversification of life in terrestrial habitats has been traditionally discussed based on the time of divergence of land plants (embryophytes), which is constrained within a Paleozoic diversification (73, 74). However, recent inferences based on phylogenomic reconstructions and molecular clocks have suggested that modern eukaryotic lineages, like Fungi and green algae, diverged on land no later than Cryogenian (73, 74). Congruently, our estimated times for Arcellinida terrestrialization are constrained within the Neoproterozoic. These findings suggest a Neoproterozoic establishment of relatively complex terrestrial ecosystems inhabited by diverse organisms, including phototrophic (green algae), absorptive heterotrophic (Fungi), and phagotrophic heterotrophic protists.

The inferred divergence of Arcellinida sub-clades in terrestrial habitats, well represented by Glutinoconcha (currently the best-sampled suborder), is also congruent to the diversification timing estimated for other eukaryotic groups. The Glutinoconcha infraorders’ split is mostly constrained between the late-Neoproterozoic and mid-Paleozoic (Devonian Period). The radiation of Fungi and the diversification of extant land plants are estimated to the same window (ca. 480 Ma) (75). Subsequently, the origin of extant Arcellinida groups, represented by the origin of all Glutinoconcha infraorders, is mostly constrained between the Silurian and Cretaceous. This is contemporaneous with the documented Late Paleozoic diversification of seed plants and saprotrophic mushrooms (75). Similarly, the estimated time for the divergence of Arcellinida genera, mostly post-dating early Mesozoic, is congruent to the radiation of diverse groups of Fungi and land plants, including pine trees (Pinaceae) and flowering plants (Angiosperm) (75). Altogether, congruences between the timing of the origin of various eukaryotic groups suggest an integrated and synchronous diversification of life in terrestrial habitats, enabling speculation about a possible radiation of Arcellinida in this time period. However, this claim requires explicit testing and corroboration via well-sampled studies of background diversification rates using fossils.

## Conclusions

Timing the origin of modern Arcellinida testate amoebae and the divergence times of their subclades illuminate the evolution of heterotrophic microbial eukaryotes in geological time. To estimate this timing, we expanded the phylogenomic sampling of amoebozoan testate amoebae and generated robust time-calibrated Amorphea trees based on both Arcellinida and Metazoa fossil records. The estimated times for the origin of Arcellinida and other amoebozoans, mostly constrained within the Tonian Period, are congruent with the previously speculated “Tonian revolution” for a diversification of eukaryotes in this Period. This consistency suggests that Earth’s ecosystems had witnessed the divergence of both phototrophic and heterotrophic eukaryotic lineages, including Arcellinida, during the Neoproterozoic, no later than the Tonian Period. A putative radiation of eukaryotes before the Cryogenian Period, coupled with the exploitation of refugia habitats and dormancy strategy, may have contributed to their endurance during Earth’s most severe ice ages. Although the ancestral habitat of Arcellinida and the possibility of transition between environments (marine vs. terrestrial) remain contentious, considering the plausible alternative scenarios we infer that Arcellinida were most likely already inhabiting terrestrial habitats between the Tonian and Ediacaran Periods. Together with the previously suggested diversification of Fungi and green algae on land during the Cryogenian Period, the inferred time for terrestrial Arcellinida is congruent with a Neoproterozoic establishment of relatively complex ecosystems composed of phototrophic (green algae), absorptive heterotrophic (Fungi), and phagotrophic heterotrophic eukaryotes, preceding the diversification of land plants. Similarly, the estimated post-Silurian origin of modern Arcellinida (i.e., infraorders) suggests a contemporaneity to the diversification of other groups, including diverse Fungi and land plants. Ultimately, we suggest the Arcellinida testate amoebae are a key group to further explore the diversification of heterotrophic microbial eukaryotes and the establishment of ecosystems starting in the Neoproterozoic.

## Material and Methods

### Sampling, RNA extraction, and sequencing

We generated transcriptomes for 14 previously genomically unsampled amoebozoan testate amoeba species through mRNA extraction from either monoclonal cultures or single-cells isolated from natural samples (**Dataset S01, Table S1**). We synthesized cDNA from RNA extractions using an adaptation of the Smart-seq2 protocol (77). We prepared our cDNA libraries for sequencing on the Illumina platform using a NexteraXT DNA Library Preparation Kit (Illumina) following the manufacturer’s recommended protocol. Libraries were then pooled and sequenced (**Appendix S01, SI1; Dataset S01, Table S1**).

### Trimming, Transcriptome Assembly, and Quality Assessment

We trimmed primers, adaptors, and low-quality bases from raw Illumina reads using Trimmomatic v. 0.36 (77). We then assembled the surviving reads with Trinity v. 2.12.0 (78). We predicted amino acid sequences (proteomes) from the assembled transcriptomes using Transdecoder v. 5.5.0. Finally, we assessed the completeness of all newly sequenced transcriptomes using BUSCO v. 5.3.2 (79) (**Dataset S01, Table S1**). Further details on trimming, transcriptome assembly, and quality assessment are presented in **Appendix S01, SI1.**

### Phylogenomic dataset construction

We constructed our amoebozoan phylogenomic dataset using the database and tools available in PhyloFisher v. 1.2.11 (37) following the steps outlined at https://thebrownlab.github.io/phylofisher-pages/detailed-example-workflow and in Jones et al. (38). Our final concatenated matrix used in subsequent phylogenetic analyses consisted of 226 genes (70,428 amino acid sites) and 57 amoebozoan taxa (**Dataset S01, Table S4**). From each individual ortholog that was concatenated to create the aforementioned matrix, we constructed single ortholog trees to be used as input for coalescent-based phylogenomic analyses. Further details on our approach for phylogenomic dataset construction are presented in **Appendix S01, SI1.**

### Phylogenomic analyses

We performed maximum likelihood phylogenetic reconstruction using our final matrix with IQ-TREE2 v. 2.0-rc (80). We initially inferred a tree from our matrix under the LG+C20+G4 model. We used the resulting tree as a guide tree to infer a Posterior Means Site Frequency model (81) using the ML model LG+C60+G4+PMSF in IQ-TREE2. We assessed the topological support for the resulting tree by 100 Real nonparametric Bootstrap replicates in IQ-TREE (IQ-TREE v. 2.1.2 COVID-edition) using the PMSF model. The topological support values inferred from MLRB were mapped onto the ML tree using RAxML v. 8.2.12 (82) using the “-f b” option. We carried out coalescent-based phylogenomic analyses with ASTRAL-III v. 5.7.3 (83). Statistical support for our ASTRAL-III analysis was assessed via local posterior probability values.

### Bayesian molecular dating

#### Dataset and Topology

Utilizing previously identified orthologs already present in the publicly available PhyloFisher database, we expanded our amoebozoan phylogenomic dataset to include a representative sampling of Amorphea. Amorphea is the eukaryotic clade composed of Amoebozoa, Fungi, Animals, and some other unicellular lineages. These ortholog sequences were aligned, trimmed, and concatenated as described above (*Phylogenomic dataset construction section*). Our final expanded dataset used in the subsequent phylogenetic reconstruction and for the molecular dating analysis consisted of 230 genes (73,467 amino acid sites) and 96 taxa (**Dataset S01, Table S7**). Maximum likelihood phylogenetic analysis was performed as described above (*Phylogenomic analyses section*).

#### Fossil calibrations

As external calibration information, we considered fossils to calibrate five internal nodes of the Metazoa clade and explored three different strategies to calibrate amoebozoan nodes (**Dataset S01, Tables S8 - S10**). We strictly derived the fossil calibration for Metazoa from dos Reis et al. (42) and Benton et al., (84), which have carefully evaluated the diversity of fossils available and calibration strategies for the Metazoa lineage. We derived the fossil calibration for Arcellinida based on previous analyses and interpretations of the morphological relationship between VSMs and extant Arcellinida (19, 21, 22). Currently, VSM’s can be interpreted either as: i) the fossil record of stem Arcellinida; ii) basal crown Arcellinida closely related to Arcellinida common ancestor, or; iii) derived crown Arcellinida (19, 21, 22). Specifically, interpretations of Bayesian and maximum likelihood ancestral reconstructions of Arcellinida shell morphology suggest a morphological congruence between the VSM *Melanocyrillium* to the Glutinoconcha+Organoconcha hypothetical ancestor and between the VSM *Cycliocyrillum* to Glutinoconcha hypothetical ancestor, thus suggesting they may represent derived crown Arcellinida (21). Consequently, three different calibration strategies can be implemented from these alternative interpretations. VSM’s can be considered to calibrate: i) Glutinoconcha+Organoconcha and Glutinoconcha nodes; ii) calibrate the Arcellinida node, or; iii) calibrate the node shared between Arcellinida and its closest sister group, the amoebozoan order Euamoebida. Aiming for a comprehensive approach we considered these three strategies and generated comparable time tree estimations. To constrain the VSMs ages, we considered the literature which has described these microfossils and refined their stratigraphical distribution (**Dataset S01, Tables S9;** 36). Since molecular dating considers the fossil information as statistical distributions, and different distributions may impact the time estimation differently, we followed dos Reis et al. (42) strategy and used a total of three different distributions to represent the calibrations derived from the fossil record: i. uniform; ii. skew-normal; and iii. truncated-cauchy short-tail. Full details on our approach for fossil calibration are presented in **Appendix S01, SI1.**

#### Divergence time estimation

We performed Bayesian molecular dating with the MCMCTree program, implemented within the PAML package (**Dataset S01, Table S10**; 85). We estimated a mean substitution rate of 0.03135 replacement site^-1^10^-8^Myr^-1^ for our dataset with IQ-TREE2 v. 2.0.6 (80) phylogenetic dating under the LG+G model. Within the PAML package, we set the overall substitution rate (‘rgene_gamma = α, β’ parameter) as a gamma-Dirichlet prior following dos Reis et al. (42), with α = 2 and β = 63.78. This substitution rate was implemented for all dating experiments. For the rate drift parameter (‘sigma^2^_gamma = α, β’), we independently implemented two alternatives, α = 2 and β = 2 or α = 1 and β = 10. Similarly, we considered both uncorrelated and autocorrelated relaxed clock models. For all experiments, we analyzed the data under the LG+G model as a single partition, constrained the root age between 1.6 - 3.2 Ga, and considered a uniform birth-death tree prior and 100 million years as one time unit. We performed a total of 36 experiments, considering three distributions (i.e., uniform, skew-normal, and truncated-Cauchy short-tail), varying the rate drift parameter (i.e., α = 2 and β = 2 or α = 1 and β = 10) and clock model (i.e., uncorrelated and autocorrelated). For each experiment, we ran two independent MCMC chains to verify convergence, discarding the first 2,000 iterations as burn-in and considering the following 20 million generations. To check the influence of fossil calibrations using Neoproterozoic VSMs on the estimated dates, we performed experiments calibrating only the nodes within the Animal clade, applying Uniform and Skew-Normal calibration strategies, under an uncorrelated or autocorrelated relaxed clock model with a drift parameter of α = 2 and β = 2 or α = 1 and β = 10, following the same approach described above, as detailed in **Appendix S01, SI1.**

### Ancestral reconstruction of Arcellinida habitat

We applied a maximum likelihood (ML) method, implemented in BayesTraits v. 4.0.1 (86), to statistically reconstruct the ancestral habitat states of Arcellinida and compare evolutionary scenarios. Currently, the diverse Tonian VSM record has been documented from environments described as fully or partially marine, suggesting the organisms represented by these fossils inhabited marine environments (19, 25). However, we cannot rule out the possibility of a terrestrial habitat for these organisms, since their dead remains could have been transported from terrestrial to marine environments where they fossilized. To explore this issue, we combine the fossil evidence and the phylogenomic reconstruction with branch lengths to reconstruct the potential ancestral habitat states (i.e., marine habitat *vs.* terrestrial habitat) of key Arcellinida clades through the BayesTraits MultiState method (86). Specifically, we used the 100 topologies with branch lengths obtained for the phylogenomic Real Bootstrap topological support assessment (see *Phylogenomic analyses section*) and implemented four different ancestral reconstruction analyses: i. ancestral reconstruction without fossilizing (assigning a fixed ancestral state value) nodes; ii. ancestral reconstruction fossilizing Arcellinida node as terrestrial, which interprets the organisms represented by VSMs as terrestrial; iii. ancestral reconstruction fossilizing Arcellinida node as marine, which interprets the organisms represented by VSMs as marine; iv. ancestral reconstruction fossilizing Arcellinida and Organoconcha+Glutinoconcha nodes as marine, which interprets the organisms represented by VSMs as derived crown Arcellinida that lived in marine habitat (**DatasetS01, Table S12 and Appendix S01, Fig. S44**). To compare the reconstructed scenarios, we applied a likelihood ratio test (LRT), which we considered as significant a difference of LRT ≥ 2 (87).

## Data Availability

Raw sequencing files are deposited at the NCBI SRA repository under the Bioproject PRJNA1032600. Phylogenomic supermatrix, single gene marker datasets, and input information for the molecular clock and ancestral reconstruction analyses are presented in DatasetS01. All molecular data associated with this manuscript are available on FigShare (10.6084/m9.figshare.25749276). This includes transcriptome assemblies, predicted proteomes, alignments (trimmed and untrimmed), as well as phylogenetic trees.

## Acknowledgments

We are deeply indebted to Susannah Porter for providing thoughtful insights to an earlier version of this manuscript, as well as a second anonymous reviewer. We thank Enrique Lara for the discussions and critiques during the construction of this project. We thank Cori Tice for her help with R and advice on plot aesthetics. This work was supported in part by the Fundação de Amparo à Pesquisa do Estado de São Paulo (FAPESP) grants 2019/22692-8 and 2021/09529-0 awarded to A.L.P.S. and grant 2019/22815-2 awarded to D.J.G.L., and by the United States National Science Foundation (NSF) Division of Environmental Biology (DEB) grant 2100888 (http://www.nsf.gov) awarded to M.W.B. KD was funded by the German Research Foundation grant 399699069 - DU 1863/1, Y.E. was supported by NSERC grant 298366-2019 to Alastair Simpson (Dalhousie University).

