## Supplemental information (SI) for "Amoebozoan testate amoebae illuminate the diversity of heterotrophs and the complexity of ecosystems throughout geological time"

**This PDF file includes:**

SI1: Material and methods

SI2: *A resolved tree of amoebozoan testate amoebae*

SI3: Taxonomic Actions

Figures S1 to S44

Legends of supplemental Dataset S01, Tables S1 to S12

SI references

**SI1: Material and methods**

**Sampling, RNA extraction, and sequencing.** We newly generated transcriptomes for 14 previously genomically unsampled amoebozoan testate amoeba species from cultured or freshly isolated specimens. We established cultures of *Galeripora* sp. from single-cells isolated from a brackish inlet on Mississippi's Gulf Coast (Biloxi, Mississippi - USA; **Dataset S01, Table S1**) and maintained as monoclonal cultures at room temperature in a medium composed of cerophyl boiled in filtered water from the source of isolation. For *Galeripora* sp. we generated 3 single-cell transcriptomes. We acquired cultures of *Spumochlamys bryora* from Culture Collection of Algae and Protozoa (CCAP) and maintained as monoclonal cultures at room temperature in NCL - 0.025% (w/v) cerophyl boiled in PJ medium (1). For *Spumochlamys bryora* we generated a whole-culture transcriptome. We established cultures of *Trichosphaerium* sp. strain KSS from single-cells isolated from Kejimkujik National Park Seaside (Nova Scotia - Canada; **Dataset S01, Table S1**) and maintained as monoclonal cultures at room temperature containing *Synechocystis* (cyanobacteria) as food source. For *Trichosphaerium* sp. strain KSS we generated a single-cell transcriptome. To obtain transcriptomes of *Phryganella* we used previously established cultures of *Phryganella* cf. *acropodia, Phryganella acropodia* strain A, and *Phryganella paradoxa* strain T (**Dataset S01, Table S1**; 2-3). We generated two single-cell transcriptomes for *P.* cf. *acropodia* (31112 and 3116), one single-cell transcriptome for *P. acropodia* A, and one single-cell transcriptome for *P. paradoxa* T. For the remaining newly sequenced taxa we isolated single-cells directly from environmental samples, cleaned through sequential dilutions (6 times) in filter sterilized (0.02µm) Deer Park (https://www.deerparkwater.com/) spring water (DPSW), and left in petri dishes containing DPSW to starve overnight prior to single-cell RNA extraction to avoid eukaryotic food contamination. To generate the single-cell transcriptomes, we extracted RNA from single cells and synthesized double-stranded cDNA (dscDNA) using a modified version of Smart-seq2 protocol (4) that includes additional freeze-thaw steps to improve cell lysis as described in Onsbring et al. (5). To generate whole-culture transcriptomes, we performed whole-culture total RNA extraction by suspending and collecting a culture of the target testate amoebae in a sterile 15 ml conical tube. We centrifuged the culture at 4000 x g at 4 °C for 5 min to pellet the cells and discarded the supernatant. We extracted the total RNA from the pelleted amoeboid cells using TRIzol reagent (Sigma-Aldrich, St Louis MO) according to the manufacturer's protocol (TRI Reagent RNA isolation reagent). Messenger RNA was isolated using New England Biolab's NEBnext magnetic mRNA isolation kit following the manufacturer's instructions. 2.3 µl of the resulting mRNA was converted to dscDNA via Smart-seq2. For all synthesized dscDNA, we assessed the quantity and quality through Qubit and electrophoresis in 1.8x Tris-borate-EDTA (TBE) agarose gel prior to library preparation. Following the Smart-seq2 protocol, we prepared libraries to be sequenced on Illumina platform using a Nextera XT DNA Library Prep Kit (Illumina, CA). The **Dataset S01, Table S1** provides the details on sampling and transcriptome sequencing for each newly sequenced lineage.

***Trimming, Transcriptome Assembly, and Quality Assessment***. For each newly sequenced transcriptome and any publicly available testate amoeba data not generated in Lahr et al. (6), we trimmed of primers, adaptors, and low-quality bases from raw Illumina reads using the Trimmomatic v. 0.36 (7) using the parameters 2:30:10 SLIDINGWINDOW:4:5 LEADING:5 TRAILING:5 MINLEN:25. We then assembled the surviving reads with Trinity v. 2.12.0 (8). We predicted amino acid sequences (proteomes) from the assembled transcriptomes using Transdecoder.LongORrfs from the Transdecoder v. 5.5.0 package (https://github.com/TransDecoder/TransDecoder). Transcriptomes produced in Lahr et al. (2019) were previously assembled and their proteome predicted following the same strategy. We assessed the completeness of all newly sequenced transcriptomes produced in this study using BUSCO v. 5.3.2 (9) and its companion eukaryota_odbODB10 database (**Dataset S01, Table S1**).

***Phylogenomic dataset construction****.* We constructed the amoebozoan phylogenomic dataset using the database and tools found in PhyloFisher v. 1.2.11 (https://github.com/TheBrownLab/PhyloFisher) (10-11). Phylofisher database consists of 240 orthologous genes useful for inferring the deep (> 200 mya) phylogeny of eukaryotic taxa (10). We used the phylogenetically aware route of *fisher.py* to collect the putative homologs of these 240 genes from the newly predicted amoebozoan proteomes. Specifically, previously identified orthologs present in the PhyloFisher database from the amoebozoans *Arcella intermediata* (*uspiensis*), *Cryptodifflugia operculata*, and *Copromyxa protea* were used as queries in the HMMER and BLAST searches performed by *fisher.py* . For each of the 240 orthologs we queried for, we added the sequences retained by the BLAST search to their respective corresponding alignments of previously identified orthologs and paralogs from 304 eukaryotic taxa covering the known diversity of eukaryotes using *working_dataset_constructor.py*. From these alignments containing the 304 provided eukaryotic taxa and the newly added amoebozoan taxa, we constructed homolog trees using *sgt_constructor.py* and manually inspected each using ParaSorter to assure correct ortholog selection and removal of any sequences from contaminating eukaryotes if present. Final decisions regarding ortholog and paralog designations were applied to the existing database via *apply_to_db.py*. To mitigate the effect of missing data on our downstream phylogenomic analyses two measures were taken, proteomes of single-cells of the same taxon derived from cultures or natural isolation were collapsed to produce a single most-complete chimera for use in our final analyses (**Dataset S01, Table S1**) and only orthologs found in at least 33% of our final set of taxa were used in the final dataset. Our final dataset was constructed using *prep_final_dataset.py* and *matrix_constructor.py* with default settings. Our final concatenated matrix used in subsequent phylogenetic analyses consisted of 226 genes (70,428 amino acid sites) and 57 taxa. Our single ortholog trees used as input for coalescent-based phylogenomic analyses were constructed using *sgt_constructor.py*.

To construct our taxonomically expanded dataset of Amorphea, we added previously identified orthologs from diverse amorpheans to the orthologs selected for newly sequenced taxa identified above using *prep_final_dataset.py*. Our matrix was then created using *matrix_constructor.py* with default settings. Again, to mitigate the effect of missing data on our downstream phylogenomic analyses proteomes of single-cells of the same taxon derived from cultures or natural isolation were collapsed to produce a single most-complete chimera for use in our final analyses (**Dataset S01, Table S1**) and only orthologs found in at least 33% of our final set of taxa were used in the final dataset. Our final expanded dataset used in the subsequent phylogenetic reconstruction and for the molecular dating analysis consisted of 230 genes (73,467 amino acid sites) and 96 taxa.

***SSU and COI phylogenetic analysis.*** We retrieved Small Subunit ribosomal RNA (SSU) and Cytochrome C oxidase subunit I (COI) sequences from transcriptomes through similarity searches implemented in Blast+ v. 2.14.1 (12). To perform similarity searches we built SSU and COI datasets of Arcellinida based on sequences available on NCBI and used them as queries in the BLAST searches to retrieve SSU and COI sequences from the available transcriptomes. Specifically, from each transcriptome (nucleotides) we build a local BLAST database via command *makeblastdb -in Transcriptome_FASTA -dbtype nucl*. We performed independent blastn searches (12) in each transcriptome using the built SSU or COI datasets via the command *blastn -query query_dataset -db Transcriptome_BLAST_database*. The sequences retrieved by the similarity searches were retained for downstream analysis. The remaining amoebozoan SSU and COI sequences were obtained from NCBI considering a representative diversity of amoebozoan testate amoebae and other amoebozoans to set as an outgroup. Prior to the phylogenetic reconstructions, we performed SSU and COI alignments considering the sequences retrieved from the transcriptomes and from NCBI using MAFFT v. 7.453 (13) via the command *mafft input_dataset > output_aligned_dataset*, alignments were visualized in AliView and regions containing primers and low-quality sequences were manually removed if present. We performed automated alignment trimming using trimAl v. 1.2 (14) via the command *trimal -in aligned_dataset -out output_trimmed_aligned_dataset -gt 0.7*. We obtained trimmed SSU and COI alignments composed of 764 and 1358 nucleotides, respectively, that we used for the phylogenetic analyses. We inferred all the maximum-likelihood trees using ModelFinder (15) and obtained node supports with the ultrafast bootstrap (16), both implemented in the IQ-TREE v. 2.2.2.7 software (17) via the command *iqtree -s trimmed_aligned_dataset -m TEST -bb 1000*.

**Bayesian molecular dating**

**Calibration information**

**Fossil record:** Tonian vase-shaped microfossils, including *Melanocyrillium* and *Cycliocyrillum*.

***Locality and stratigraphy:*** Tonian Chuar Group, Kwagunt Formation, Grand Canyon, Arizona and Tonian Callison Lake Formation of Yukon, Canada.

**Phylogenetic and calibration justification:** The well-preserved nature of the vase-shaped microfossils has allowed morphological interpretations that suggest these fossils have a close affinity to the amoebozoan order Arcellinida (6, 18-20). Currently, VSM's can be interpreted as the fossil record of stem arcellinids, basal crown arcellinids closely related to Arcellinida common ancestor, or even derived crown Arcellinida, and there is no evidence to justify favoring one of these interpretations over the others (6, 18-19). Regarding the interpretation of VSMs as derived crown arcellinids, analyses of Bayesian and Maximum Likelihood ancestral reconstructions of arcellinids shell morphology suggest a morphological congruence between the VSM *Melanocyrillium* to the Glutinoconcha+Organoconcha hypothetical ancestor and between the VSM *Cycliocyrillum* to Glutinoconcha hypothetical ancestor, thus suggesting they may represent derived crown Arcellinida (6). Consequently, three different calibration strategies can be derived from these alternative interpretations. VSM's can be considered to calibrate Glutinoconcha+Organoconcha and Glutinococha nodes (VSMs as derived crown arcellinids), calibrate Arcellinida node (VSM as basal crown arcellinids), or calibrate the node shared between Arcellinida and its closest sister group, the amoebozoan order Euamoebida (VSM as stem arcellinids). Thus, aiming for a comprehensive approach we consider these three strategies to generate comparable time tree estimations.

**Soft Minimum age:** 743.4 Ma (751±7.6)
**Soft Maximum age:** 750.2 Ma (757±6.8)

***Age justification:*** Currently, Tonian Chuar Group, Kwagunt Formation, Grand Canyon, Arizona and Tonian Callison Lake Formation of Yukon, Canada represent some of the best dated fossiliferous sedimentary successions bearing VSMs, including specimens of *Melanocyrillium* and *Cycliocyrillum* (18, 21). The ages of these successions have been recently refined and carefully discussed based on precise rhenium-osmium (Re-Os) and uranium–lead (U-Pb) detrital zircon geochronology (211). Thus, we consider their age constraints to determine the minimum and maximum ages of the VSM record we used for our calibrations. Importantly, the ages of Tonian Chuar Group and Tonian Callison Lake Formation are consistent with the ages estimated for other formations around the globe where VSMs are also found, however, these other formations have been dated based on less precise methods (e.g., chemostratigraphy and biostratigraphy) thus their ages are not considered. From Kwagunt Formation the oldest VSM record is found within Awatubi Member which has been dated 751±7.6 Ma and from Callison Lake Formation dated 739.9±6.1 Ma (21-22). As the minimum age for the VSM record, we considered the oldest reported evidence for these microfossils. Thus we constrain the minimum age for the VSM record as 743.4 Ma, the lower end of the geochronological uncertainty assigned to Awatubi Member, which bears the oldest reported VSM record to date. The soft maximum constraint is based on the lower end of the geochronological uncertainty of the fossiliferous sedimentary successions that lie beneath the Awatubi Member, the Carbon Canyon (757±6.8 Ma), which have been investigated for microfossils and no VSM has been reported to date (21-22).

**Calibration implementation:** Since molecular dating considers the fossil information as statistical distributions, and different distributions may impact the time estimation differently, we followed dos Reis et al. (23) strategy and used a total of three different distributions to represent the calibrations derived from the fossil record (**Dataset S01, Table S10**): i. uniform, a relatively more conservative interpretation of the fossil's minimum and maximum ages and considers that all dates between this range have the same probability; ii. skew-normal, a more literal interpretation of the fossil's minimum and maximum ages, and considers that the minimum age and younger ages have a higher probability than older ages and the maximum age; and iii. truncated-Cauchy short-tail, a looser interpretation of the fossil record which requires only the minimum age to be implemented and, thus does not consider the soft maximum constraint, which is arguably a more arbitrary date since older fossils can always be found. Consequently, implementing and comparing these three distributions aim for a comprehensive approach to conducting time tree calibrations. We used the three distributions to independently implement the calibrations considering the VSMs as derived crown arcellinids or VSMs as basal crown arcellinids, while we used only truncated-Cauchy distribution to implement the calibrations consider the VSMs as stem arcellinids, since considering fossils representative of a stem group to calibrate phylogenetic trees requires distribution that takes into consideration only minimum age constraints, such as truncated-Cauchy, to avoid underestimation of the nodes age (24). To check the influence of fossil calibrations using Neoproterozoic VSMs on the estimated dates, we performed experiments calibrating only the nodes within the Animal clade, applying Uniform and Skew-Normal calibration strategies, under an uncorrelated or autocorrelated relaxed clock model with a drift parameter of $\alpha$ = 2 and β = 2 or $\alpha$ = 1 and β = 10, following the same approach described above.

**SI2:** ***A resolved tree of amoebozoan testate amoebae***

The new phylogeny represents a significant expansion of the amoebozoan testate amoebae sampled in a phylogenomic framework and enabled the precise placement of several taxa that have been tentatively classified based on morphological data (**Appendix S01, SI3**). *Phryganella acropodia* and *Phryganella paradoxa* expand the phylogenomic sampling for the Phryganellina suborder adding its type genus (*Phryganella*). To date, *Phryganella paradoxa* and *Phryganella acropodia* have been molecularly sampled and placed within Phryganellina based on SSU rDNA and COI, although their monophyletic relationship was not resolved arguably due to lack of a better taxonomic sampling or longer marker sequences (3, 25). Here we observed *P. acropodia*, the type species of Phryganellina genus, branching as a more basal group of Phryganellina while *Phryganella paradoxa* branches sister to *Cryptodifflugia operculata*. Taking into account their phylogenetic placement and morphology, here we suggest that the diversity and classification of the genera *Phryganella* and *Cryptodifflugia* require clarification from further sampling based on morphological and molecular studies. The oldest described genus of Arcellinida, *Arcella* Ehrenberg 1830, was recently split into two different genera, *Arcella* and *Galeripora*, based on COI marker and morphology (26). The phylogenomic placement of the newly sequenced *Galeripora* sp. and the relatively long branch length corroborate the differentiation of these two genera. The addition of *Netzelia lobostoma* and *Cyclopyxis* sp. expands the sampling of Sphaerothecina, within which *Cyclopyxis lobostoma* and *Cyclopyxis* sp. do not form a monophyletic group. Interestingly, the placement of *Difflugia* cf. *capreolata* within Sphaerothecina expands the phylogenetic diversity of the *Difflugia*-like taxa. Difflugidae is one of the most diverse Arcellinida families but has been consistently shown to be paraphyletic (27). It is worth noting that most Sphaerothecina taxa have not yet been sampled, which may explain the unresolved node within this infraorder. Given that the type species of *Cyclopyxis* (*Cyclopyxis arcelloides*) and *Difflugia* (*Difflugia proteiformis*) have not been molecularly sampled and phylogenetically placed, here we do not take a taxonomic action regarding *Difflugia* cf. *capreolata* and *Cyclopyxis* paraphyly. A further expansion of morphological and molecular investigation is welcomed to guide taxonomic actions that will lead to a stable and consistent clarification of Sphaerothecina diversity and classification.

Another enigmatic lineage we sampled is *Heleopera lucida* comb. nov. (previously *Difflugia lucida*), an Arcellinida originally classified as a member of Longithecina characterized by its rigid bilaterally symmetrical and laterally compressed shell, which is 50-70 µm long and 30-40 µm width (28), with terminal aperture. However, *H. lucida* comb. nov. also shares resemblances to other taxonomic entities, such as the characteristic laterally compressed shells covered with quartz particles found within *Heleopera* (e.g., *Heleopera rosea* and *Heleopera baetica*, Heleoperidae:Volnustoma infraorder). The similarity to *Heleopera* has been already noted in its original description (28), but *H. lucida* comb. nov. lacks a key character traditionally used for the identification of *Heleopera* species (but absent in the original description of the genus by Joseph Leidy; 29), the presence of a conspicuous organic lip around the terminal aperture (30). Here, our phylogenomic reconstruction shows the placement of *H. lucida* comb. nov. within Volnustoma infraorder, illuminating the relevance of their similarity with *Heleopera* previously noted (28). Another recently described Heleoperidae corroborates the evolutionary affinity of *H. lucida* comb. nov. morphotype to the *Heleopera* genus, *Heleopera baetica* has been investigated based on morphology and SSU and, similarly to *H. lucida* comb. nov., does not have the characteristic organic lip of the genus, expanding the morphological diversity of the group (31). Accordingly, here we transfer *H. lucida* comb. nov. as a species of the genus *Heleopera* (**Appendix S01, SI3**).

The precise phylogenomic placement of *Microcorycia* and *Spumochlamys* within Arcellinida modifies the classification of the amoebozoan testate amoebae with flexible shells (**Appendix S01, SI3**).

The current classification separates amoebozoan amoebae with a flexible shell into three different families. Within Arcellinida are the Microchlamiidae family, which includes *Microchlamys*, *Spumochlamys,* and *Pyxidicula* (6), while within Corycidia are the Microcoryciidae family, which includes *Microcorycia*, *Diplochlamys*, *Amphizonella*, *Penardochlamys*, *Zonomyxa,* and *Parmulina* (32-33)*,* finally, the Trichosidae family includes *Trichosphaerium* (33). While previous phylogenomic reconstructions support the precise position and classification of *Microchlamys, Diplochlamys, Amphizonella*, and *Trichosphaerium* (6, 33), the remaining flexible shell-bearing taxa, including the type genus *Microcorycia* (Microcoryciidae) as well as *Spumochlamys* (Microchlamiidae), have been tentatively classified based on morphology. *Spumochlamys* has also been investigated using SSU ribosomal RNA, however, its precise placement has been impaired by the lack of robustness of this single marker and a characteristic long branch for the taxon (34). Our phylogenomic reconstruction robustly places *Spumochlamys* within Organoconcha. Similarly, the fully supported placement of *Microcorycia aculeata* and *Microcorycia flava* within Organoconcha leads to transferring the *Microcorycia* genus from Corycidia to the order Arcellinida. Although initially classified within Corycidia, this phylogenomic placement is supported by morphology and raises a homologous trait of Organoconcha previously overlooked. *Microcorycia*'s hemispherical shell, with an aperture as wide as the widest breadth of the shell, is remarkably similar to *Microchlamys, Spumochlamys,* and *Pyxidicula*. Specifically, the rigid *Pyxidicula* shell has a thin flexible veil on the margin of the aperture, which may be homologous to the flexible lower region of *Microchlamys*, *Spumochlamys,* and *Microcorycida* shells. *Microcorycia* is the type genus for the family Microcoryciidae de Saedeleer 1934, as such, the name must be transferred with the taxon. In its new home, *Microcorycia* falls within the already established family Microchlamyiidae Ogden 1985, and as such must be treated as subjective synonyms in a family containing *Microcorycia*, *Microchlamys, Spumochlamys,* and *Pyxidicula*. Concomitantly, we raise Amphizonellidae fam. nov. (Corycidia) to accommodate the remaining *Amphizonella*, *Diplochlamys*, *Penardochlamys*, *Zonomyxa*, and *Parmulina* (**Appendix S01, SI3**).

**SI3: Taxonomic Actions**

Importantly, the placement of enigmatic lineages sampled in this study leads to a significant modification in the current classification of amoebozoan testate amoebae, since taxa have been moved from one monophyletic group (Corycidia) to the other (Arcellinida). The following classification complements the one in Lahr et al., (6), which is the most recent comprehensive classification of amoebozoan testate amoebae. As such, only the changes are recorded here in detail. We attempt to reconcile the prevailing

Phylocode usage in higher-level protistan classifications with the traditional classification of testate amoebae based on the International Code of Zoological Nomenclature (35). Our ultimate goal is to maintain the stability of names, while additionally generating a classification congruent with current phylogenetic data. We propose here to continue using the Phylocode (36) as the organizational backbone, because it is more easily applicable to higher ranks in protists (37). We adopt the system established by Adl and collaborators for indicating hierarchical levels in protists (38-40), where increasing numbers of bullet points indicate less inclusive and nested hierarchical levels. However, we also suggest how these hierarchical levels integrate with the Linnean ranks, which are a legacy of hundreds of years of classification. Therefore, the highest taxon treated here (Arcellinida), is indicated by "•••" and is at the Linnean rank of "Order", those indicated by "••••" are at the level of "Suborder", those indicated by "•••••" are at the level of "Infraorder".

**Amorphea** Adl et al. 2012

**Amoebozoa** Lühe 1913, *sensu* Cavalier-Smith 1998

• **Tubulinea** Smirnov et al. 2005

•• **Elardia** Kang et al. 2017
••• **Order Arcellinida** Kent 1880
•••• **Suborder Glutinoconcha** 6
••••• **Infraorder Volnustoma** 6
•••••• Family Heleoperidae Jung 1942
••••••• Genus *Heleopera* Leidy 1879
•••••••• *Heleopera lucida* Penard 1890 comb. nov.
Synonym *Difflugia lucida* Penard 1890

Observations: *Heleopera lucida* comb. nov. was originally described in the *Difflugia* genus (28). Here we demonstrate that the organism branches at a basal position to the *Heleopera* using a phylogenomic approach. Combined with morphological congruence with the original description for genus *Heleopera*, we suggest a new combination, transferring the taxon to the genus *Heleopera*. Noteworthy morphological congruences are a compressed shell, composed of agglutinated particles (in this case minera), and an elliptical to slit-like aperture. Note that while many recent authors include an organic lip in the aperture as a defining feature, this is NOT present in the original description of the genus.

•••• **Suborder Organoconcha** Lahr et al 2019

•••••• *Family:* Microchlamyiidae Ogden 1985

We now include the genus *Microcorycia* Cockerell 1911

••• **Order Corycidia**

•••••• *Family:* Amphizonelliidae fam. nov. Porfirio-Sousa, Tice, Brown, and Lahr

••••••• Genus *Amphizonella* Greeff 1866

••••••• Genus *Diplochlamys* Greeff 1888

••••••• Genus Zonomyxa Nüsslin 1882

••••••• Genus Parmulina Penard 1902

••••••• Genus Penardochlamys Deflandre 1953

**Supplementary figures**

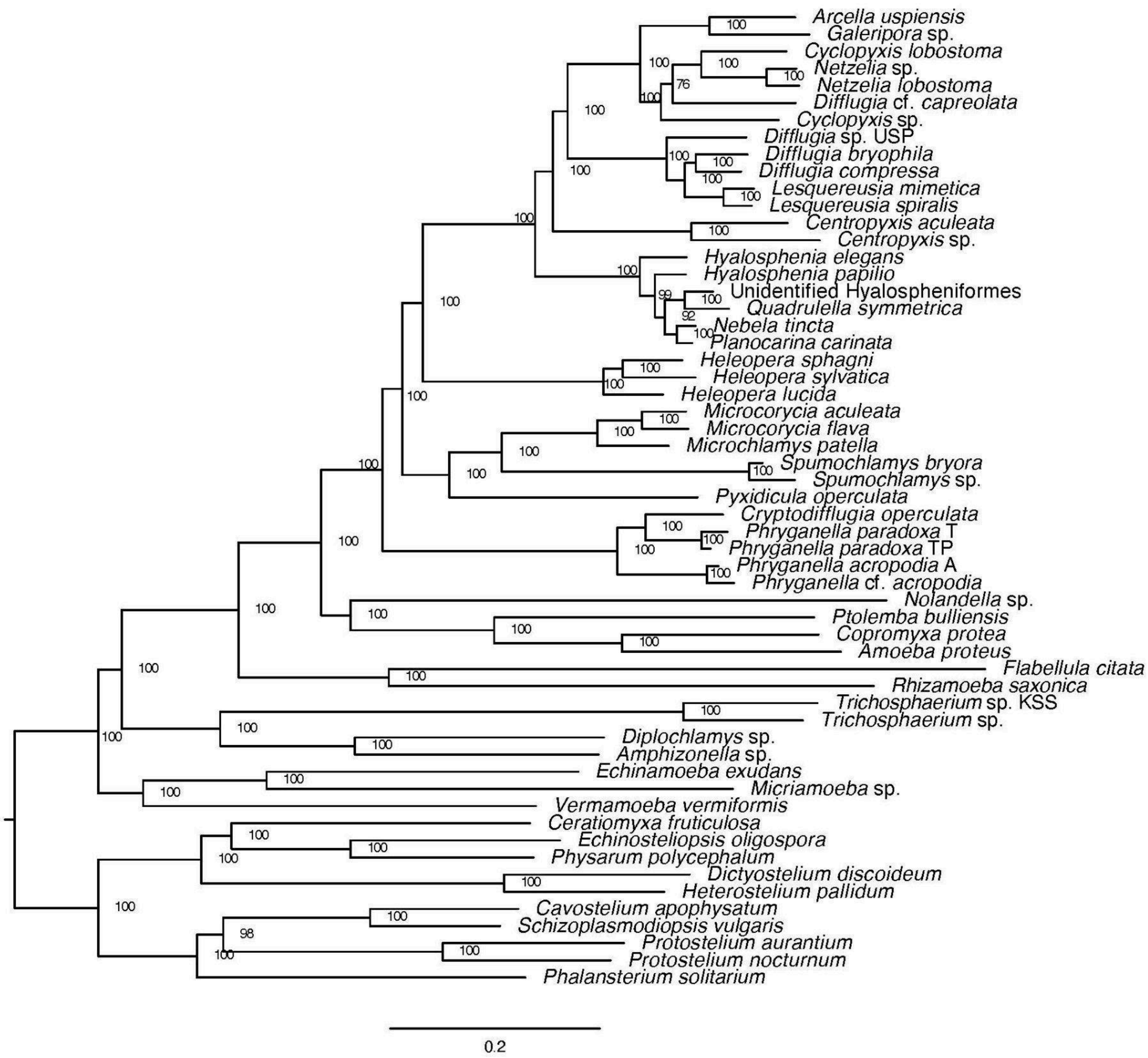


**Figure S1.** The phylogenetic tree of amoebozoan testate amoebae presented on Fig. 2. 226 gene (70,428 amino acid sites) phylogeny of amoebozoan testate amoebae rooted with Evosea (Amoebozoa). The tree was initially built using IQ-TREE2 v. 2.0-rc1 under the LG+C20+G4 model of protein evolution and further used to infer a Posterior Means Site Frequency model using the ML model LG+C60+G4+PMSF. Topological support shown on nodes was assessed by 100 Maximum Likelihood Real Bootstrap (MLRB).

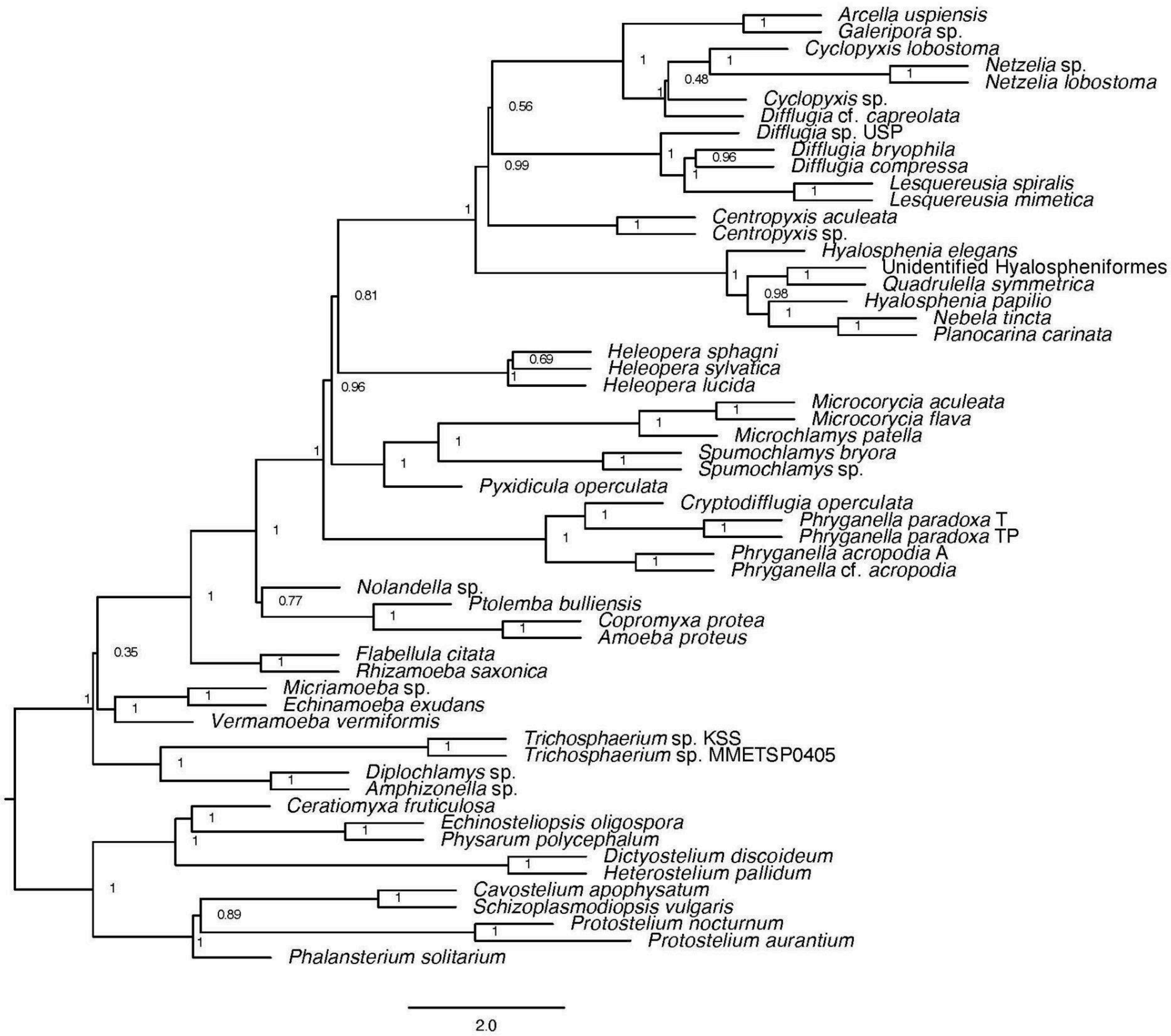


**Figure S2.** Coalescent-based phylogenomic (CP) reconstruction of amoebozoan testate amoebae using ASTRAL-III v. 5.7.3 based on 226 individual gene trees. Support values at nodes are local posterior probability values.

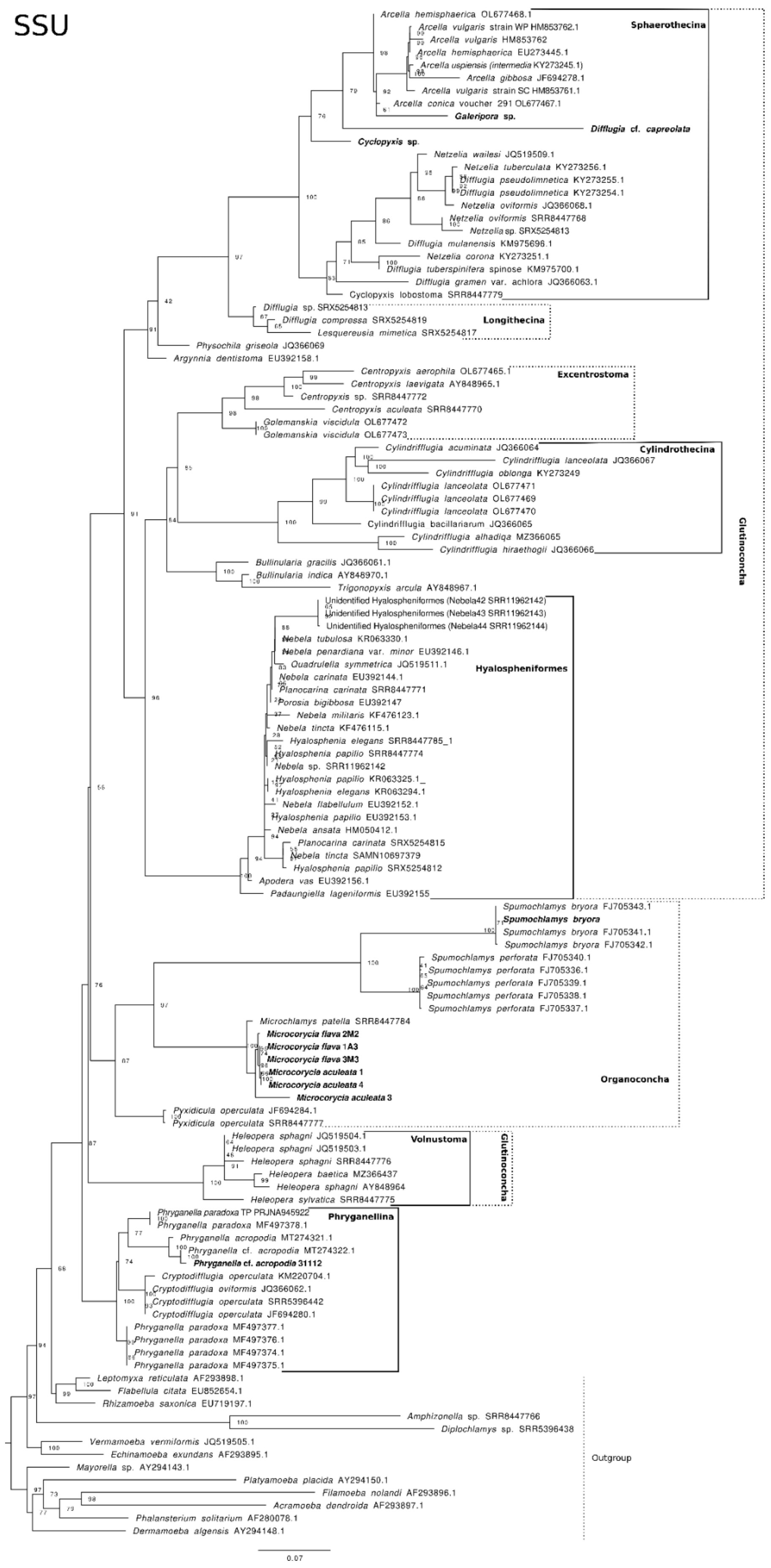
SSU
Arcella hemisphaerica OL677468.1
Sphaerothecina
Arcella vulgaris strain WP HM853762.1
Arcella vulgaris HM853762
Arcella hemisphaerica EU273445.1
Arcella uspiensis (intermedia KY273245.1)
Arcella gibbosa JF694278.1
Arcella vulgaris strain SC HM853761.1
Arcella conica voucher 291 OL677467.1
Galeripora sp.
Difflugia cf. capreolata
Cyclopyxis sp.
Netzelia wailesi JQ519509.1
Netzelia tuberculata KY273256.1
Difflugia pseudolimnetica KY273255.1
Difflugia pseudolimnetica KY273254.1
Netzelia oviformis JQ366068.1
Netzelia oviformis SRR8447768
Netzelia sp. SRX5254813
Difflugia mulanensis KM975696.1
Netzelia corona KY273251.1
Difflugia tuberspinifera spinose KM975700.1
Difflugia gramen var. achlora JQ366063.1
Cyclopyxis lobostoma SRR8447779
Difflugia sp. SRX5254813
Difflugia compressa SRX5254819
Longithecina
Lesquereusia mimetica SRX5254817
Physochila griseola JQ366069
Argynnia dentistoma EU392158.1
Centropyxis aerophila OL677465.1
Excentrostoma
Centropyxis laevigata AY848965.1
Centropyxis sp. SRR8447772
Centropyxis aculeata SRR8447770
Golemanskia viscidula OL677472
Golemanskia viscidula OL677473
Cylindrifflugia acuminata JQ366064
Cylindrothecina
Cylindrifflugia lanceolata JQ366067
Cylindrifflugia oblonga KY273249
Cylindrifflugia lanceolata OL677471
Cylindrifflugia lanceolata OL677469
Cylindrifflugia lanceolata OL677470
Cylindrifflugia bacillariarum JQ366065
Cylindrifflugia alhadiqa MZ366065
Cylindrifflugia hiraethogli JQ366066
Glutinoconcha
Bullinularia gracilis JQ366061.1
Bullinularia indica AY848970.1
Trigonopyxis arcula AY848967.1
Unidentified Hyalospheniformes (Nebela42 SRR11962142)
Unidentified Hyalospheniformes (Nebela43 SRR11962143)
Unidentified Hyalospheniformes (Nebela44 SRR11962144)
Nebela tubulosa KR063330.1
Nebela penardiana var. minor EU392146.1
Quadrulella symmetrica JQ519511.1
Hyalospheniformes
Nebela carinata EU392144.1
Planocarina carinata SRR8447771
Porosia bigibbosa EU392147
Nebela militaris KF476123.1
Nebela tincta KF476115.1
Hyalosphenia elegans SRR8447785_1
Hyalosphenia papilio SRR8447774
Nebela sp. SRR11962142
Hyalosphenia papilio KR063325.1
Hyalosphenia elegans KR063294.1
Nebela flabellulum EU392152.1
Hyalosphenia papilio EU392153.1
Nebela ansata HM050412.1
Planocarina carinata SRX5254815
Nebela tincta SAMN10697379
Hyalosphenia papilio SRX5254812
Apodera vas EU392156.1
Padaungiella lageniformis EU392155
Spumochlamys bryora FJ705343.1
Spumochlamys bryora
Spumochlamys bryora FJ705341.1
Spumochlamys bryora FJ705342.1
Spumochlamys perforata FJ705340.1
Spumochlamys perforata FJ705336.1
Spumochlamys perforata FJ705339.1
Spumochlamys perforata FJ705338.1
Spumochlamys perforata FJ705337.1
Microchlamys patella SRR8447784
Microcorycia flava 2M2
Microcorycia flava 1A3
Microcorycia flava 3M3
Microcorycia aculeata 1
Microcorycia aculeata 4
Microcorycia aculeata 3
Organoconcha
Pyxidicula operculata JF694284.1
Pyxidicula operculata SRR8447777
Heleopera sphagni JQ519504.1
Volnustoma
Glutinoconcha
Heleopera sphagni JQ519503.1
Heleopera sphagni SRR8447776
Heleopera baetica MZ366437
Heleopera sphagni AY848964
Heleopera sylvatica SRR8447775
Phryganella paradoxa TP PRJNA945922
Phryganellina
Phryganella paradoxa MF497378.1
Phryganella acropodia MT274321.1
Phryganella cf. acropodia MT274322.1
Phryganella cf. acropodia 31112
Cryptodifflugia operculata KM220704.1
Cryptodifflugia oviformis JQ366062.1
Cryptodifflugia operculata SRR5396442
Cryptodifflugia operculata JF694280.1
Phryganella paradoxa MF497377.1
Phryganella paradoxa MF497376.1
Phryganella paradoxa MF497374.1
Phryganella paradoxa MF497375.1
Leptomyxa reticulata AF293898.1
Flabellula citata EU852654.1
Rhizamoeba saxonica EU719197.1
Amphizonella sp. SRR8447786
Diplochlamys sp. SRR5396438
Vermamoeba vermiformis JQ519505.1
Echinamoeba exundans AF293895.1
Outgroup
Mayorella sp. AY294143.1
Platyamoeba placida AY294150.1
Filamoeba nolandi AF293896.1
Acramoeba dendroida AF293897.1
Phalansterium solitarium AF280078.1
Dermamoeba algensis AY294148.1
0.07

**Figure S3.** The maximum-likelihood Small Subunit ribosomal RNA (SSU) tree of amoebozoan testate amoebae. The tree was built using ModelFinder (15) and obtained node supports with the ultrafast bootstrap (16), both implemented in the IQ-TREE v. 2.2.2.7 software (17). Based on ultrafast bootstrap a clade is considered well supported if its support is >= 95% (16). Continuous lines delimit well supported clades and dashed lines delimit clades reconstructed with low ultrafast bootstrap (<95%) or paraphyletic. The taxa with newly sequenced transcriptomes from which we successfully retrieved SSU sequences are shown in bold.

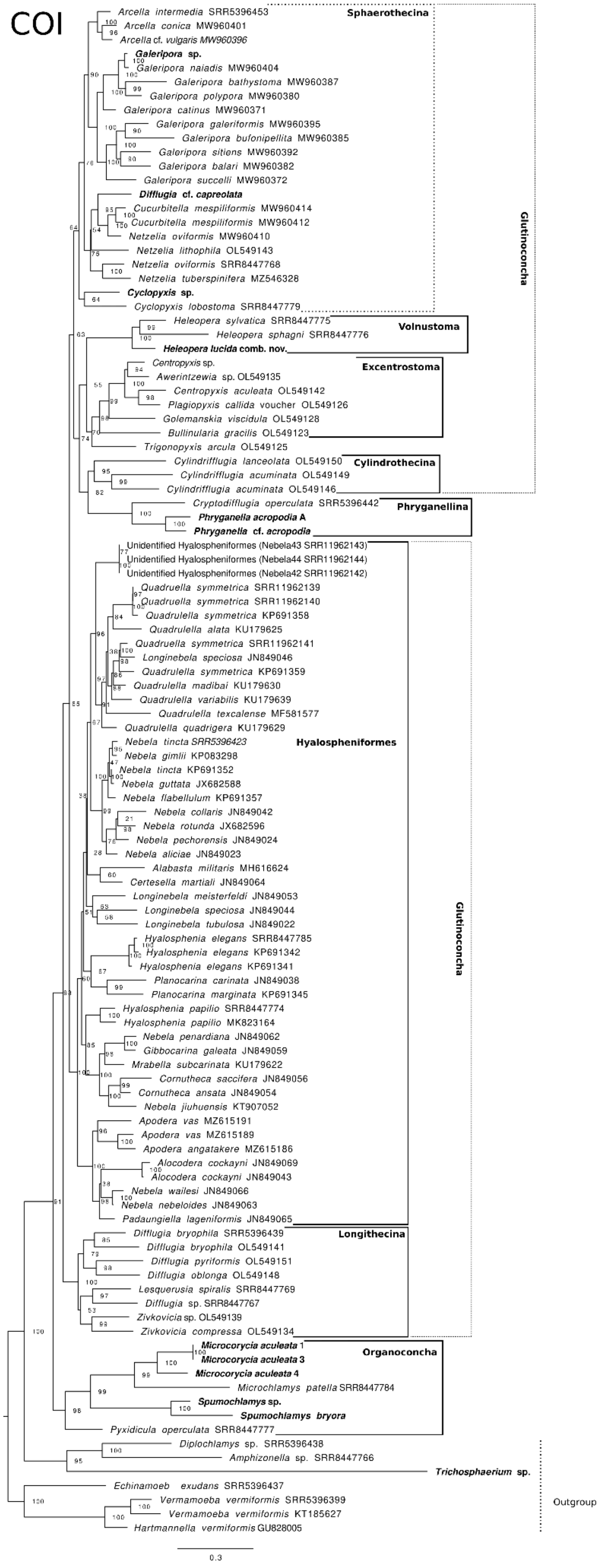
COI
Arcella intermedia SRR5396453
Arcella conica MW960401
Arcella cf. vulgaris MW960396
Galeripora sp.
Galeripora naiadis MW960404
Galeripora bathystoma MW960387
Galeripora polypora MW960380
Galeripora catinus MW960371
Galeripora galeriformis MW960395
Galeripora bufonipellita MW960385
Galeripora sitiens MW960392
Galeripora balari MW960382
Galeripora succelli MW960372
Difflugia cf. capreolata
Cucurbitella mespiliformis MW960414
Cucurbitella mespiliformis MW960412
Netzelia oviformis MW960410
Netzelia lithophila OL549143
Netzelia oviformis SRR8447768
Netzelia tuberspinifera MZ546328
Cyclopyxis sp.
Cyclopyxis lobostoma SRR8447779
Sphaerothecina
Heleopera sylvatica SRR8447775
Heleopera sphagni SRR8447776
Heleopera lucida comb. nov.
Volnustoma
Centropyxis sp.
Awerintzewia sp. OL549135
Centropyxis aculeata OL549142
Plagiopyxis callida voucher OL549126
Golemanskia viscidula OL549128
Bullinularia gracilis OL549123
Trigonopyxis arcula OL549125
Excentrostoma
Cylindrifflugia lanceolata OL549150
Cylindrifflugia acuminata OL549149
Cylindrifflugia acuminata OL549146
Cylindrothecina
Glutinoconcha
Cryptodifflugia operculata SRR5396442
Phryganella acropodia A
Phryganella cf. acropodia
Phryganellina
Unidentified Hyalospheniformes (Nebela43 SRR11962143)
Unidentified Hyalospheniformes (Nebela44 SRR11962144)
Unidentified Hyalospheniformes (Nebela42 SRR11962142)
Quadruella symmetrica SRR11962139
Quadruella symmetrica SRR11962140
Quadrulella symmetrica KP691358
Quadrulella alata KU179625
Quadruella symmetrica SRR11962141
Longinebela speciosa JN849046
Quadrulella symmetrica KP691359
Quadrulella madibai KU179630
Quadrulella variabilis KU179639
Quadrulella texcalense MF581577
Quadrulella quadrigera KU179629
Nebela tincta SRR5396423
Nebela gimlii KP083298
Nebela tincta KP691352
Nebela guttata JX682588
Nebela flabellulum KP691357
Nebela collaris JN849042
Nebela rotunda JX682596
Nebela pechorensis JN849024
Nebela aliciae JN849023
Alabasta militaris MH616624
Certesella martiali JN849064
Longinebela meisterfeldi JN849053
Longinebela speciosa JN849044
Longinebela tubulosa JN849022
Hyalosphenia elegans SRR8447785
Hyalosphenia elegans KP691342
Hyalosphenia elegans KP691341
Planocarina carinata JN849038
Planocarina marginata KP691345
Hyalosphenia papilio SRR8447774
Hyalosphenia papilio MK823164
Nebela penardiana JN849062
Gibbocarina galeata JN849059
Mrabella subcarinata KU179622
Cornutheca saccifera JN849056
Cornutheca ansata JN849054
Nebela jiuhuensis KT907052
Apodera vas MZ615191
Apodera vas MZ615189
Apodera angatakere MZ615186
Alocodera cockayni JN849069
Alocodera cockayni JN849043
Nebela wailesi JN849066
Nebela nebeloides JN849063
Padaungiella lageniformis JN849065
Hyalospheniformes
Difflugia bryophila SRR5396439
Difflugia bryophila OL549141
Difflugia pyriformis OL549151
Difflugia oblonga OL549148
Lesquerusia spiralis SRR8447769
Difflugia sp. SRR8447767
Zivkovicia sp. OL549139
Zivkovicia compressa OL549134
Longithecina
Glutinoconcha
Microcorycia aculeata 1
Microcorycia aculeata 3
Microcorycia aculeata 4
Microchlamys patella SRR8447784
Spumochlamys sp.
Spumochlamys bryora
Pyxidicula operculata SRR8447777
Organoconcha
Diplochlamys sp. SRR5396438
Amphizonella sp. SRR8447766
Trichosphaerium sp.
Echinamoeb exudans SRR5396437
Vermamoeba vermiformis SRR5396399
Vermamoeba vermiformis KT185627
Hartmannella vermiformis GU828005
Outgroup
0.3

**Figure S4.** The maximum-likelihood Cytochrome C oxidase subunit I (COI) tree of amoebozoan testate amoebae. The tree was built using ModelFinder (15) and obtained node supports with the ultrafast bootstrap (16), both implemented in the IQ-TREE v. 2.2.2.7 software (17). Based on ultrafast bootstrap a clade is considered well supported if its support is >= 95% (16). Continuous lines delimit well-supported clades and dashed lines delimit clades reconstructed with low ultrafast bootstrap (<95%) or paraphyletic. The taxa with newly sequenced transcriptomes from which we successfully retrieved COI sequences are shown in bold.

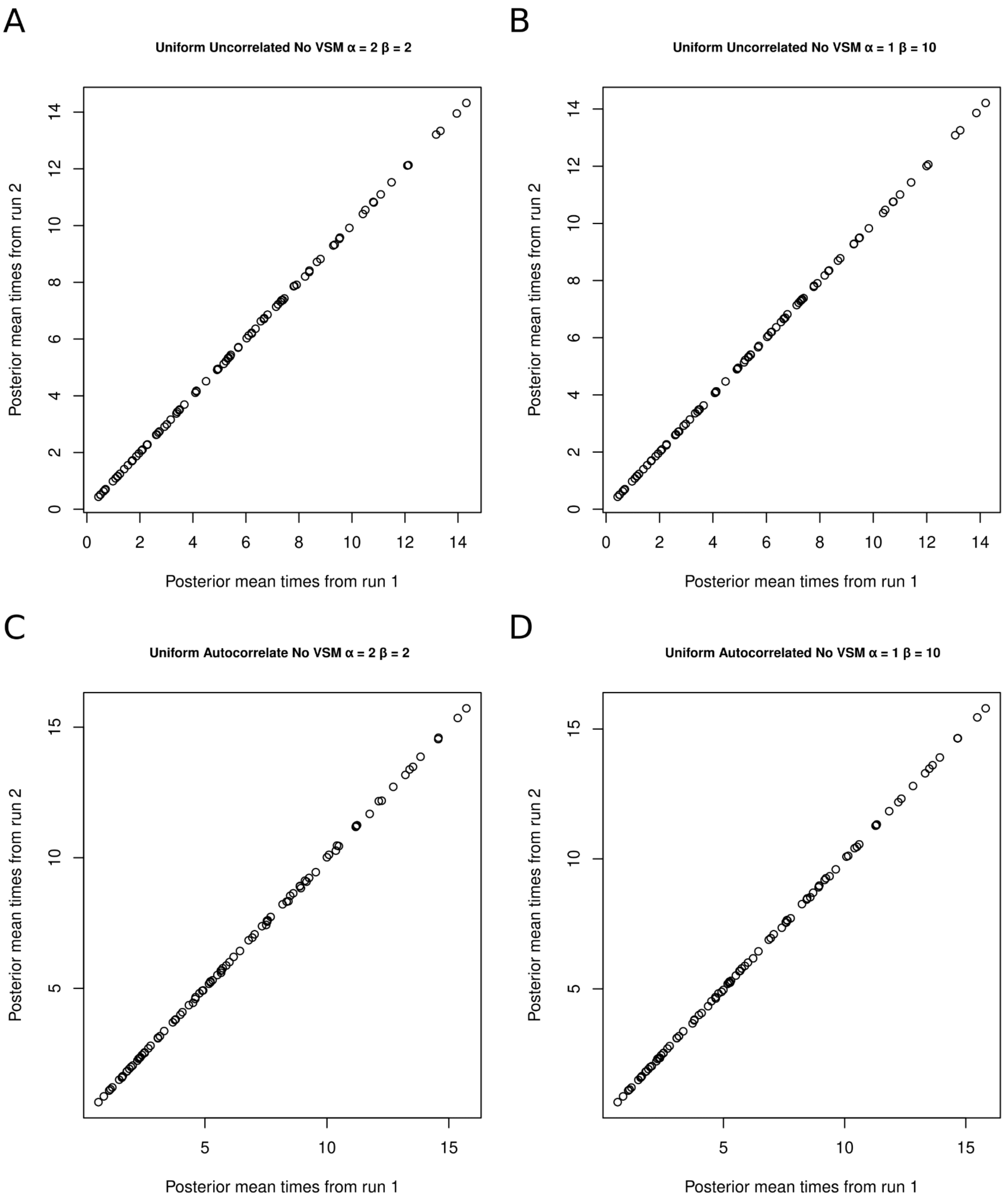


**Figure S5. Plot of mean estimated time by chain (run) 1 and chain (run) 2 of each experiment to check for convergence. The mean times represent million years, considering 100 million years as one time unit.**

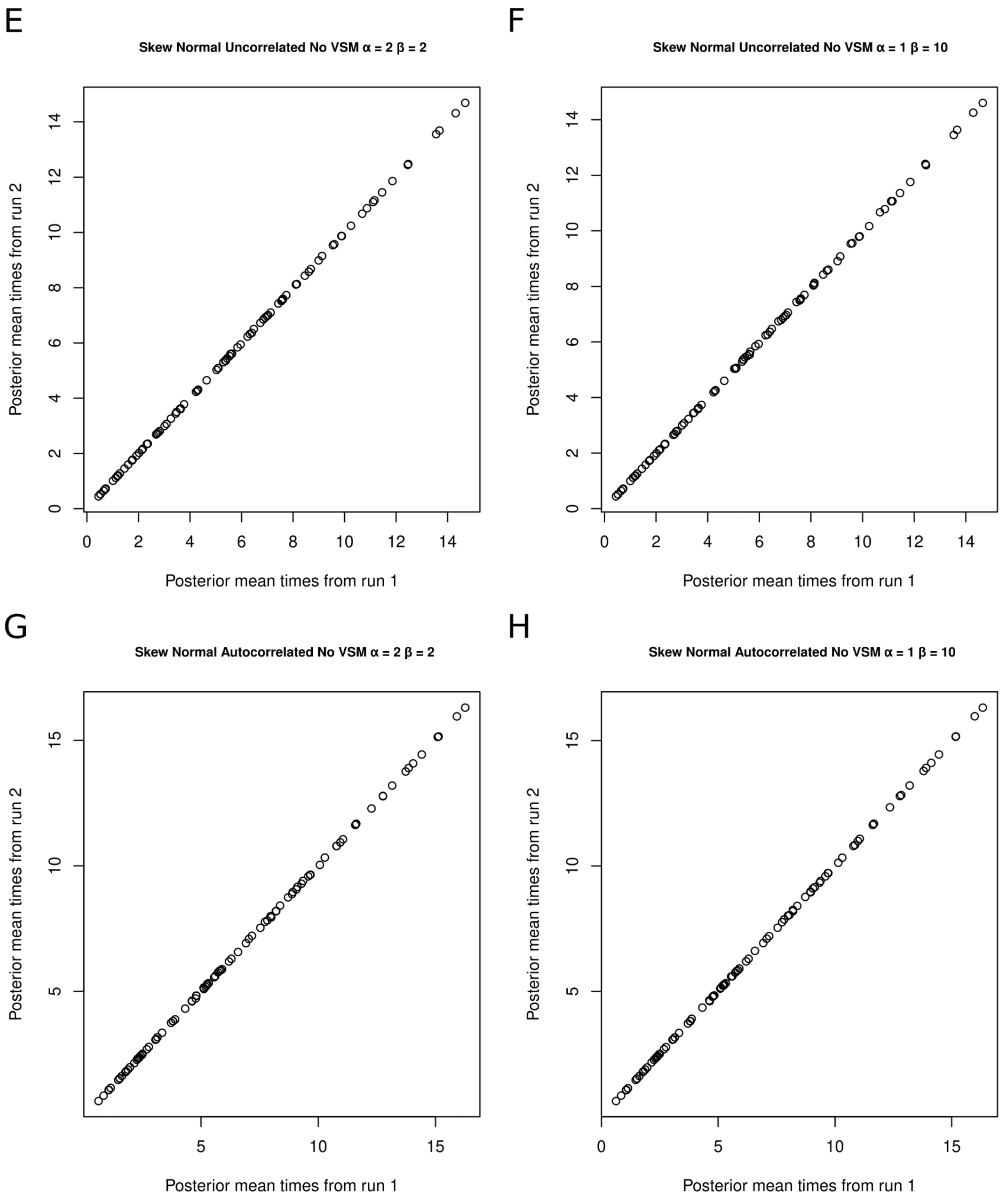


**Figure S5 (continued)**

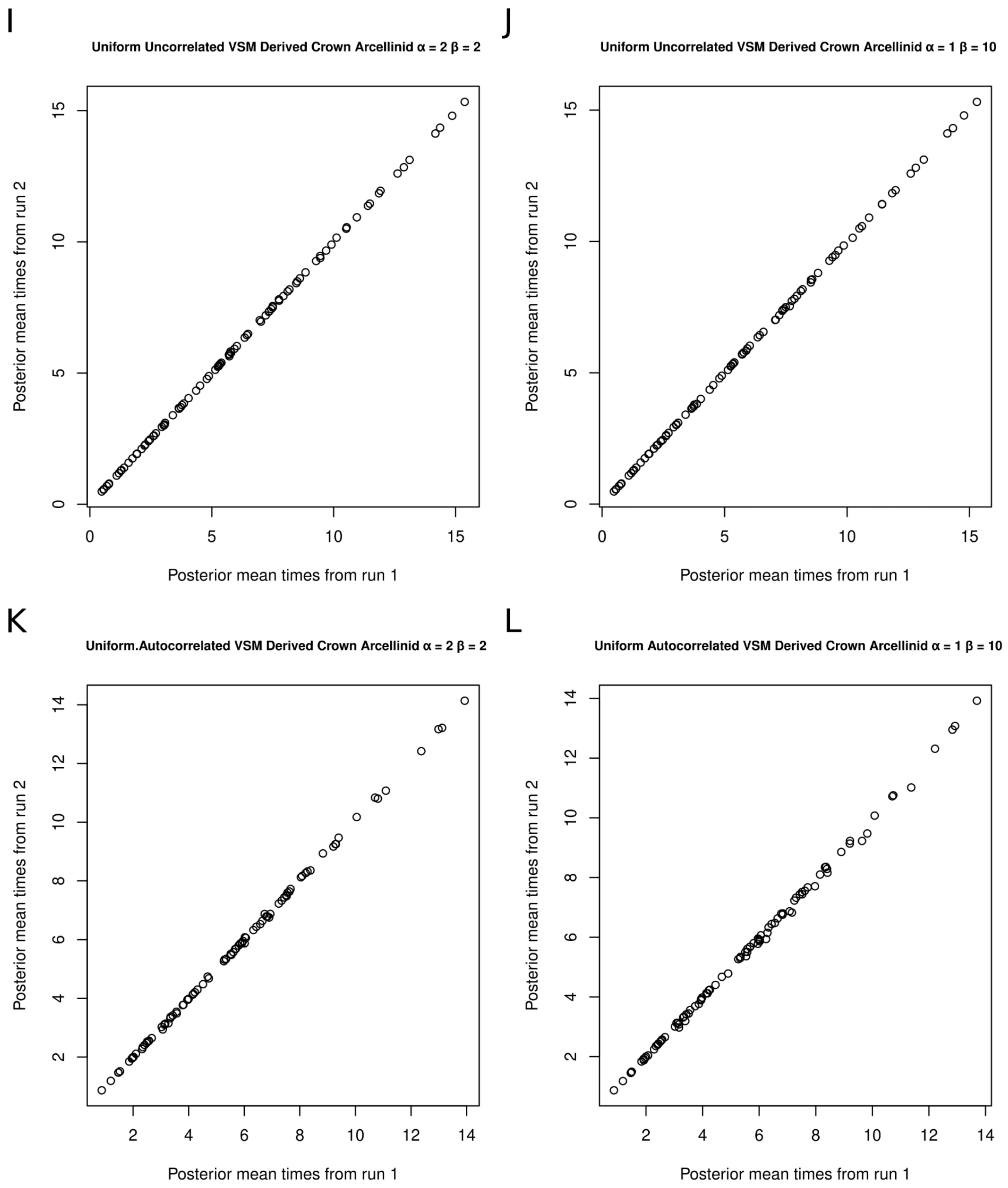


**Figure S5 (continued)**

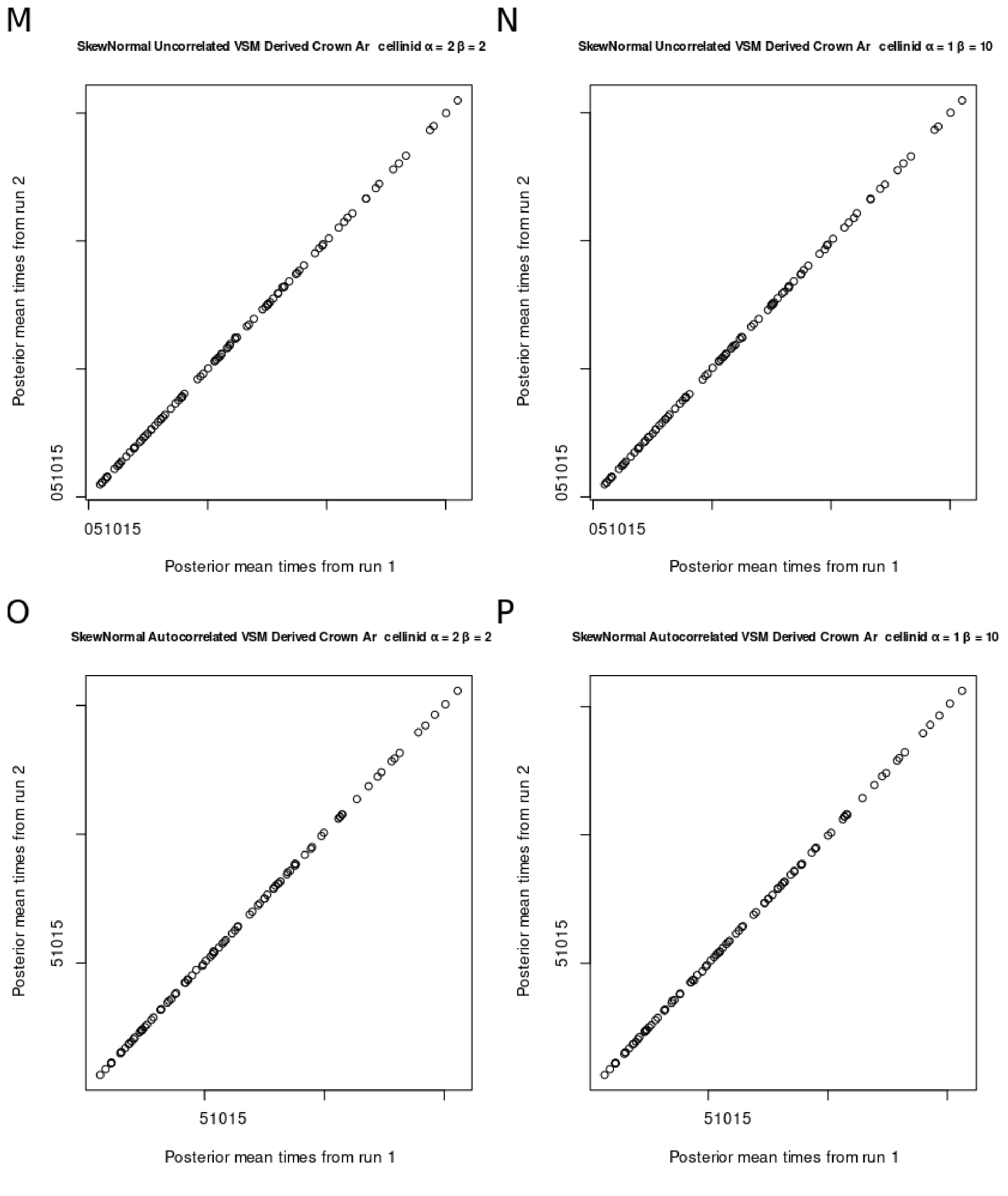


**Figure S5 (continued)**

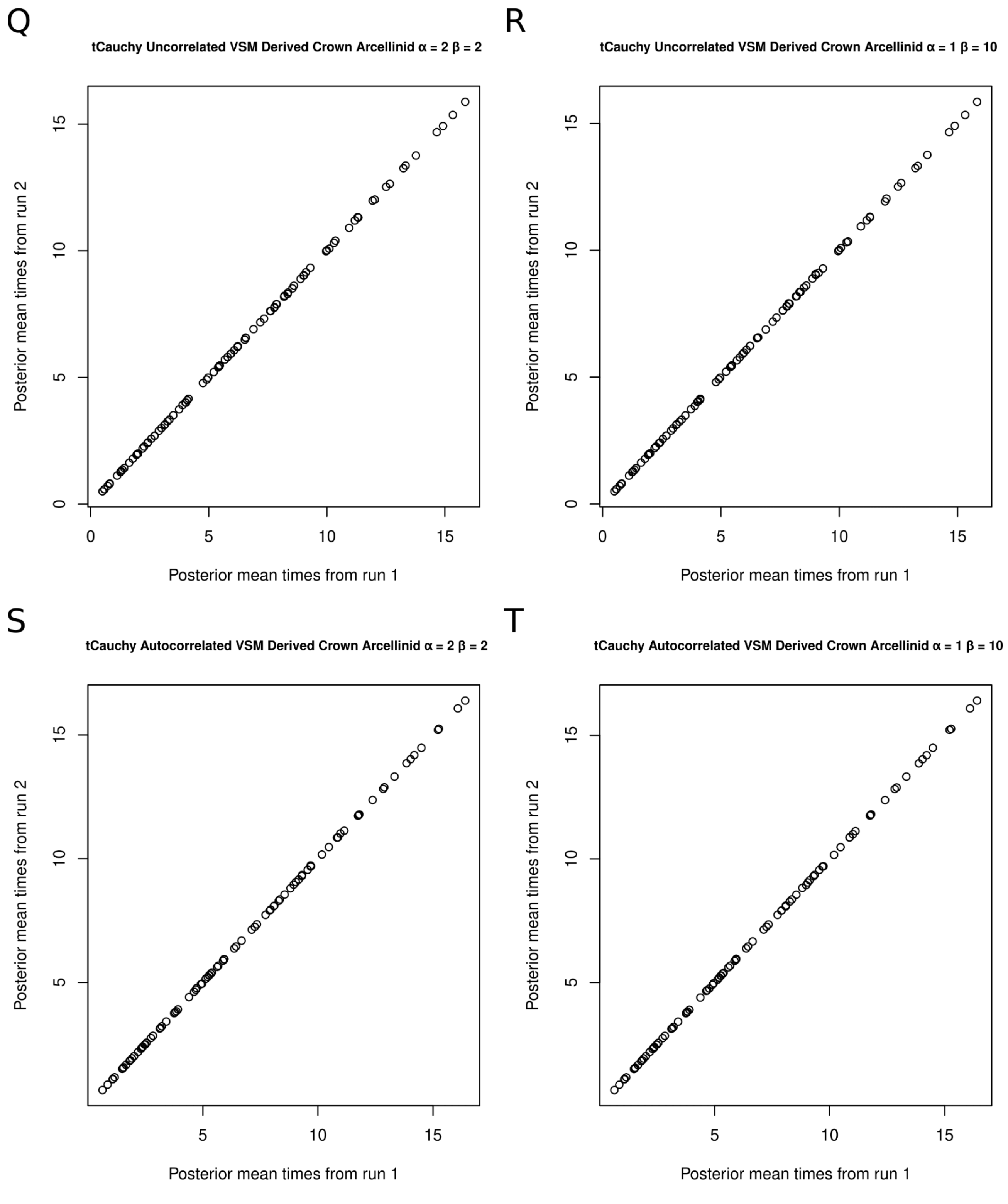


**Figure S5 (continued)**

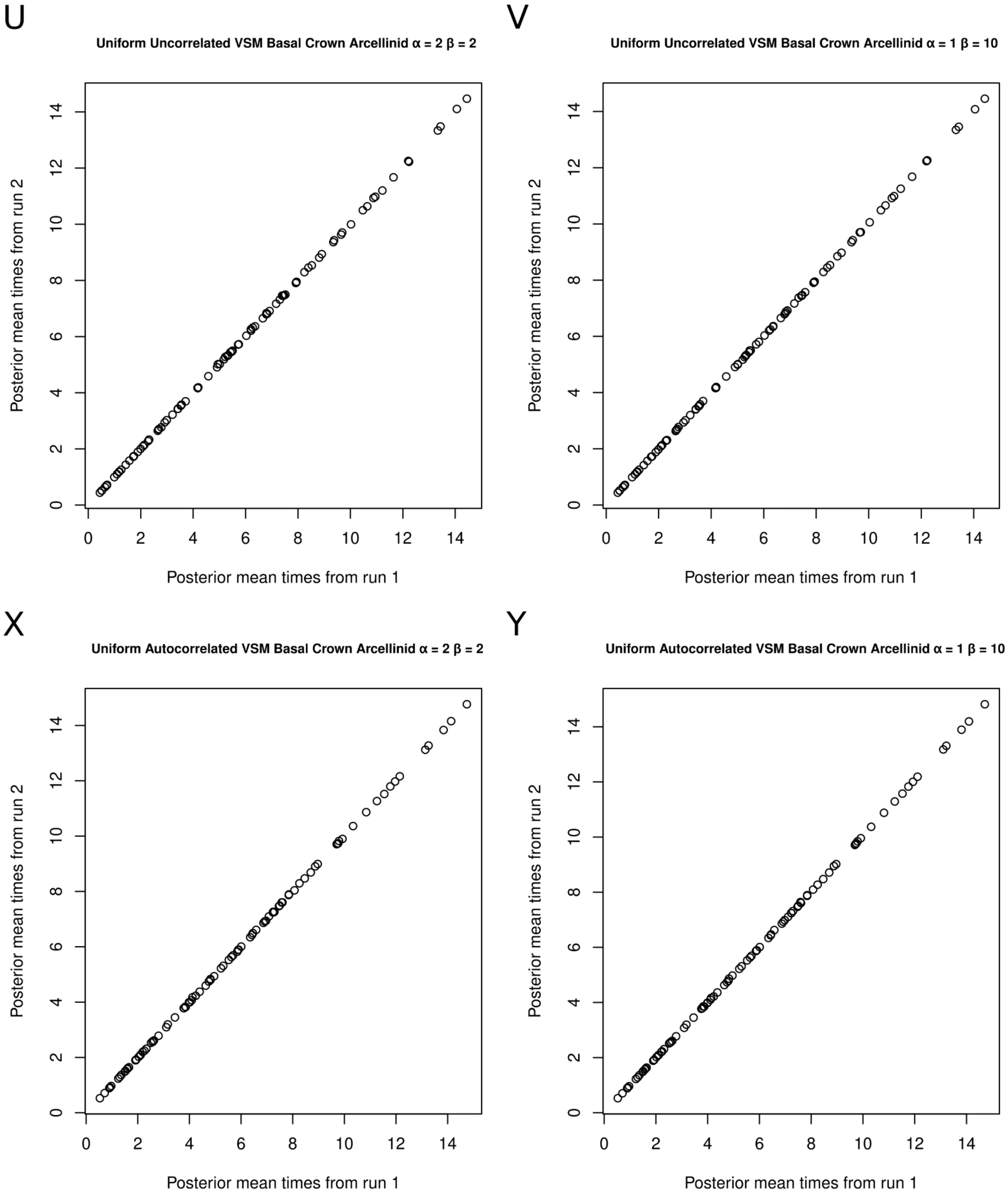


**Figure S5 (continued)**

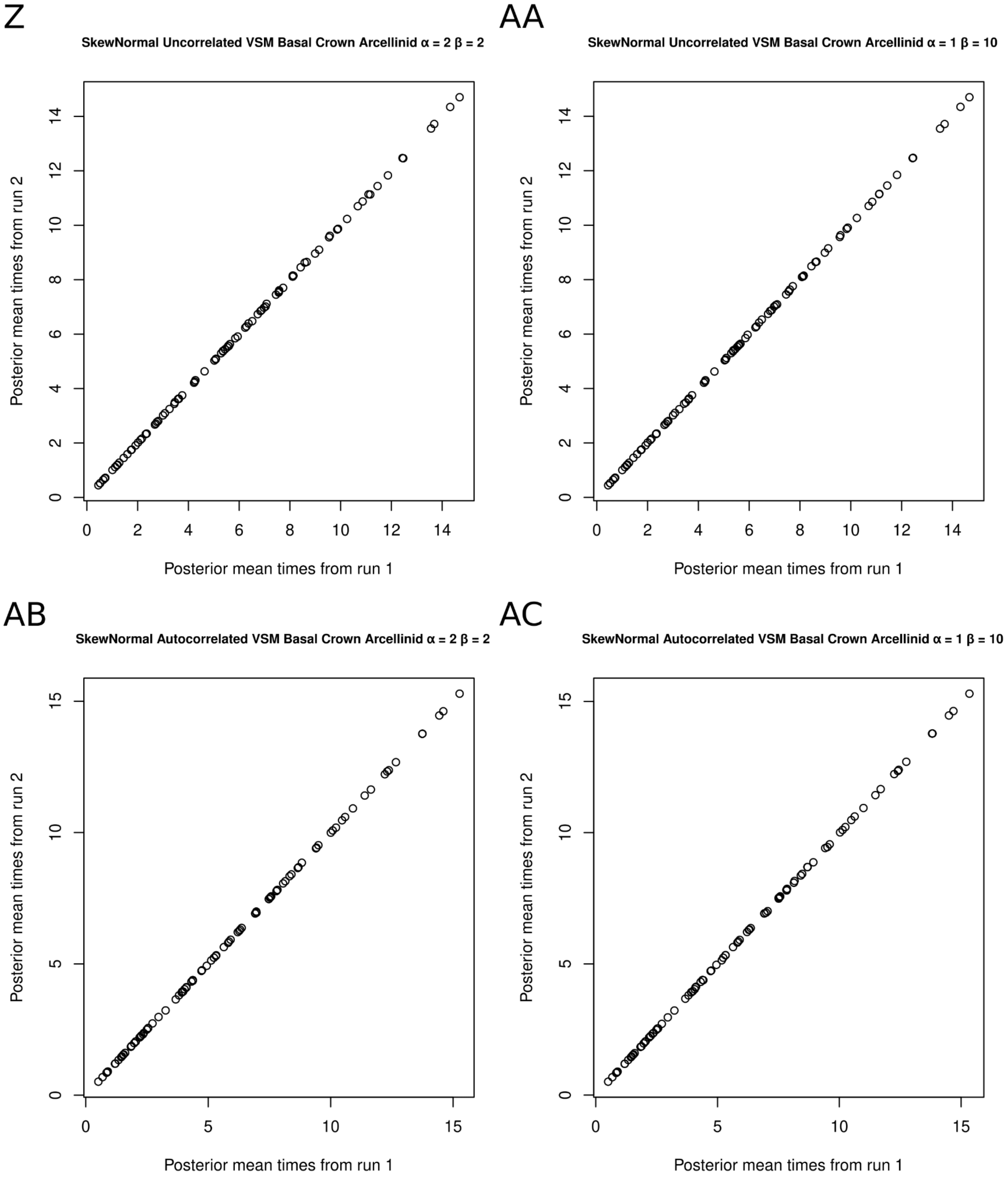


**Figure S5 (continued)**

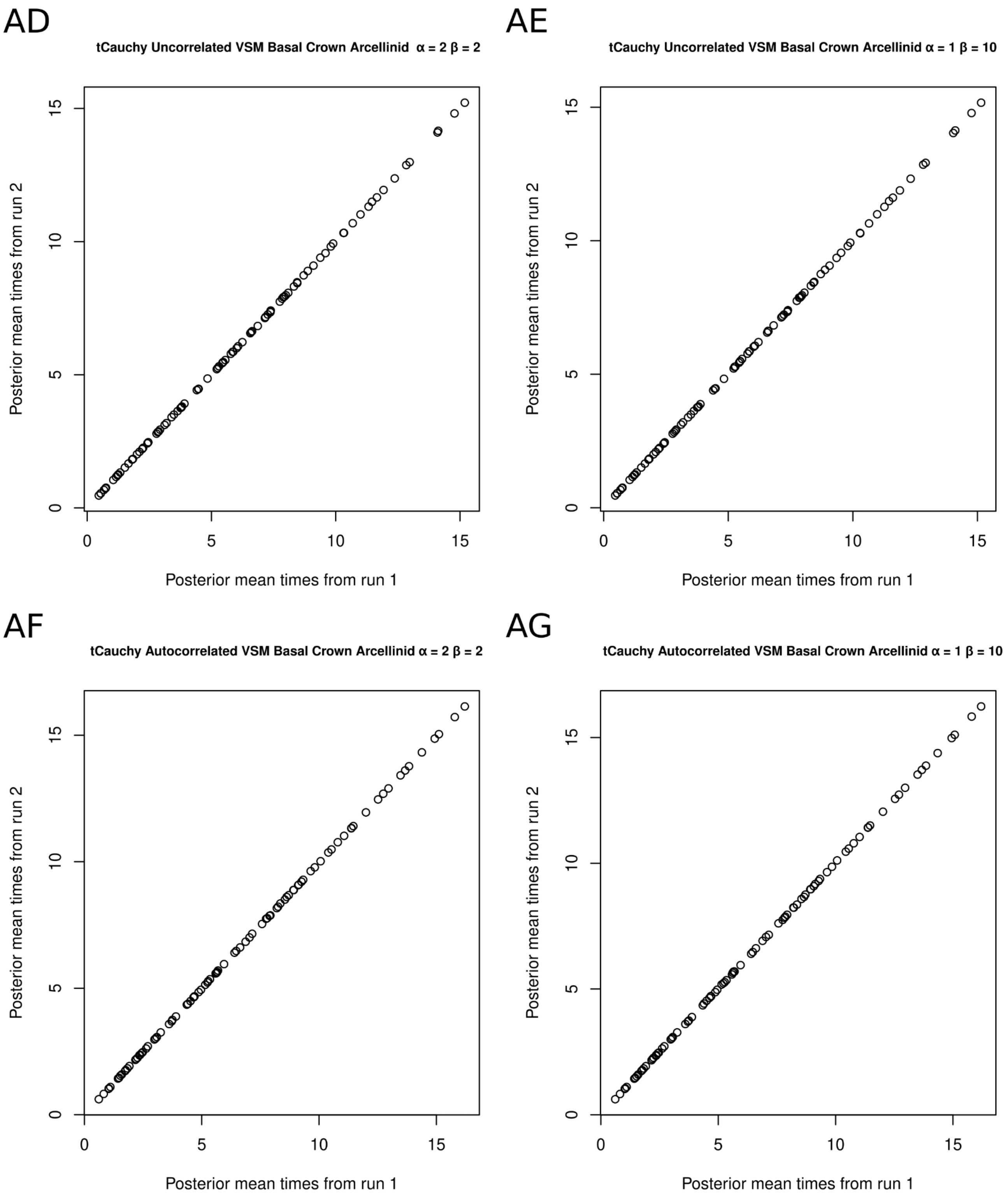


**Figure S5 (continued)**

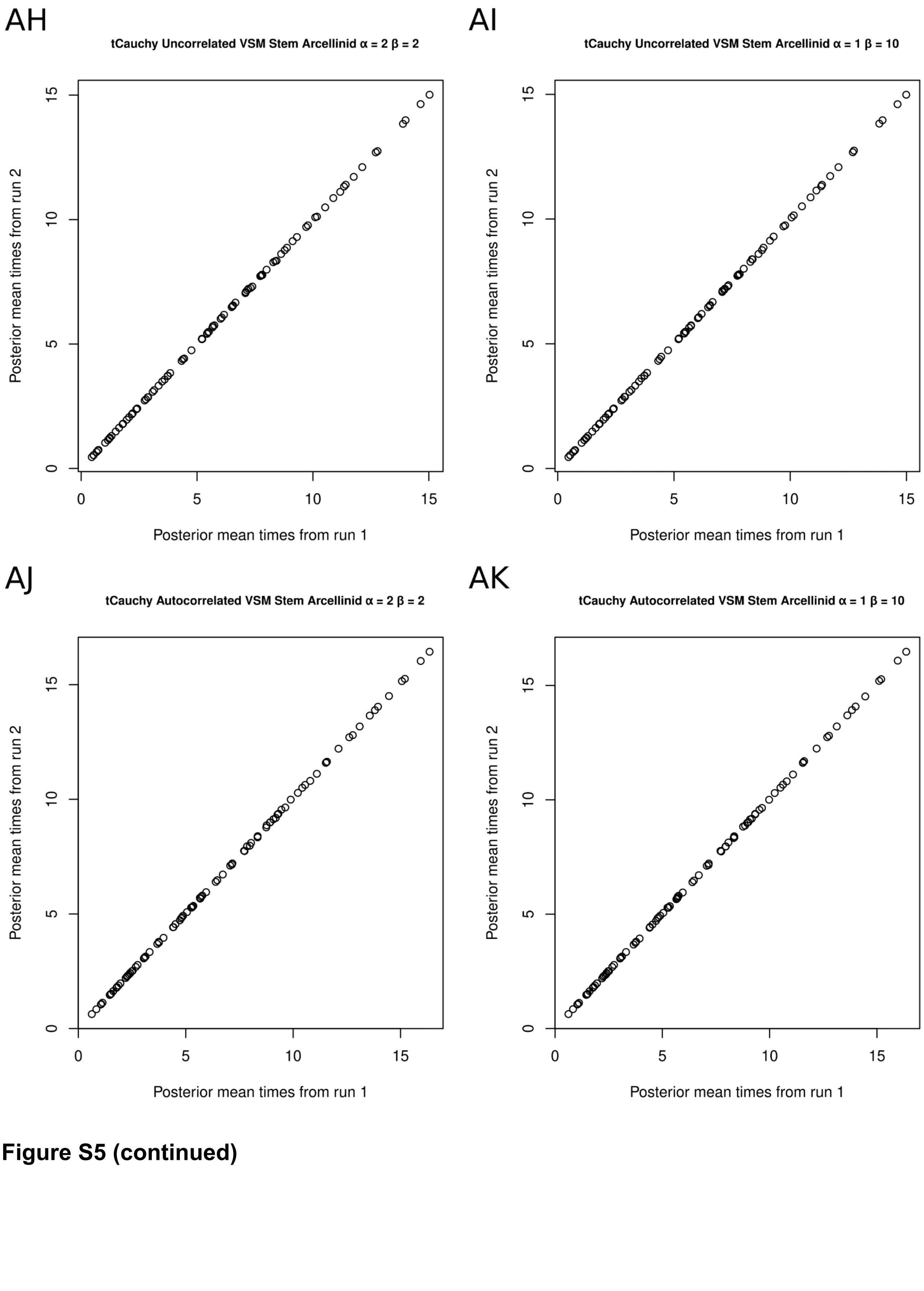


**Figure S5 (continued)**

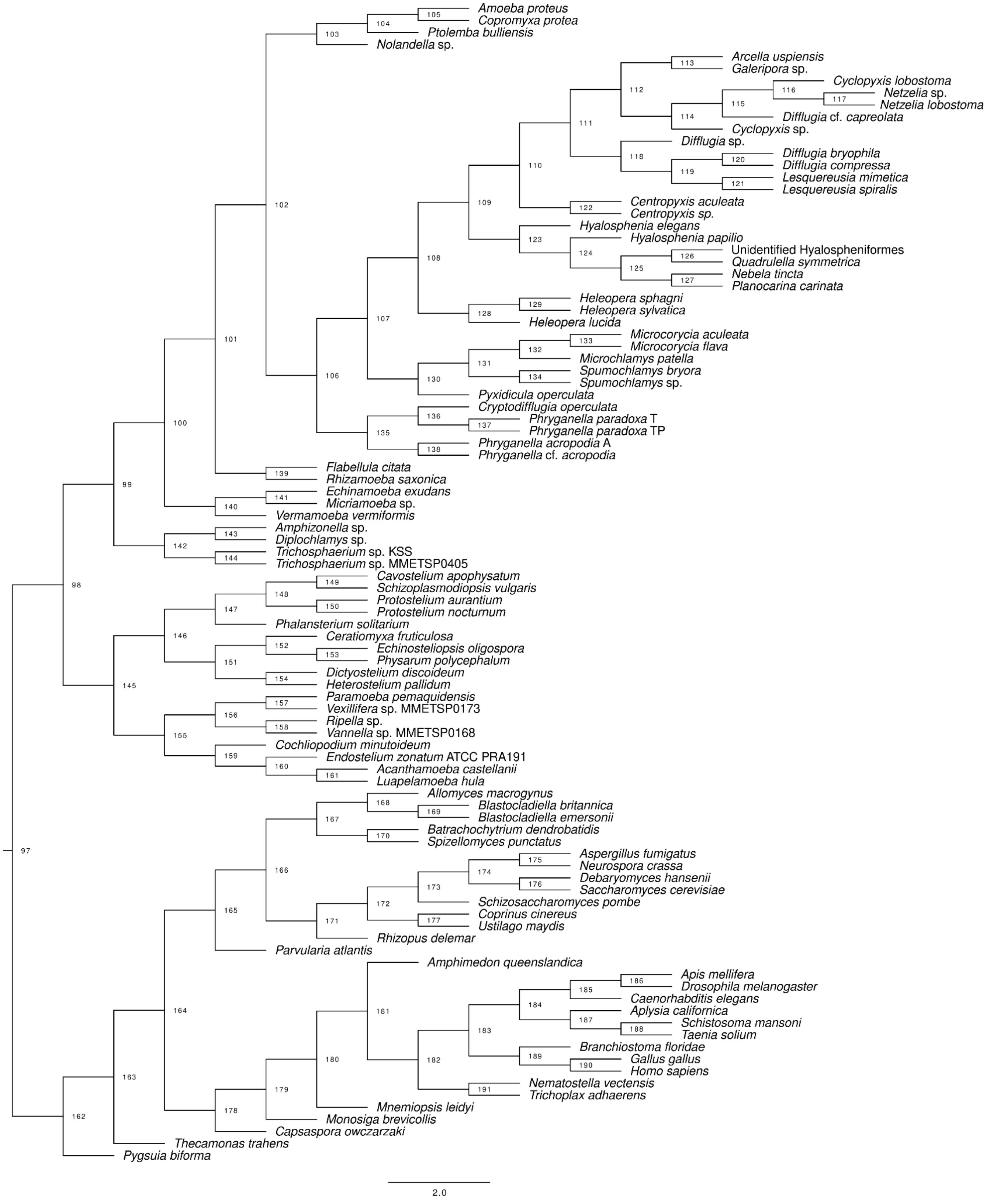


**Figure S6.** Tree with nodes annotated as appeared on **Dataset S01, Tables S10-S11**

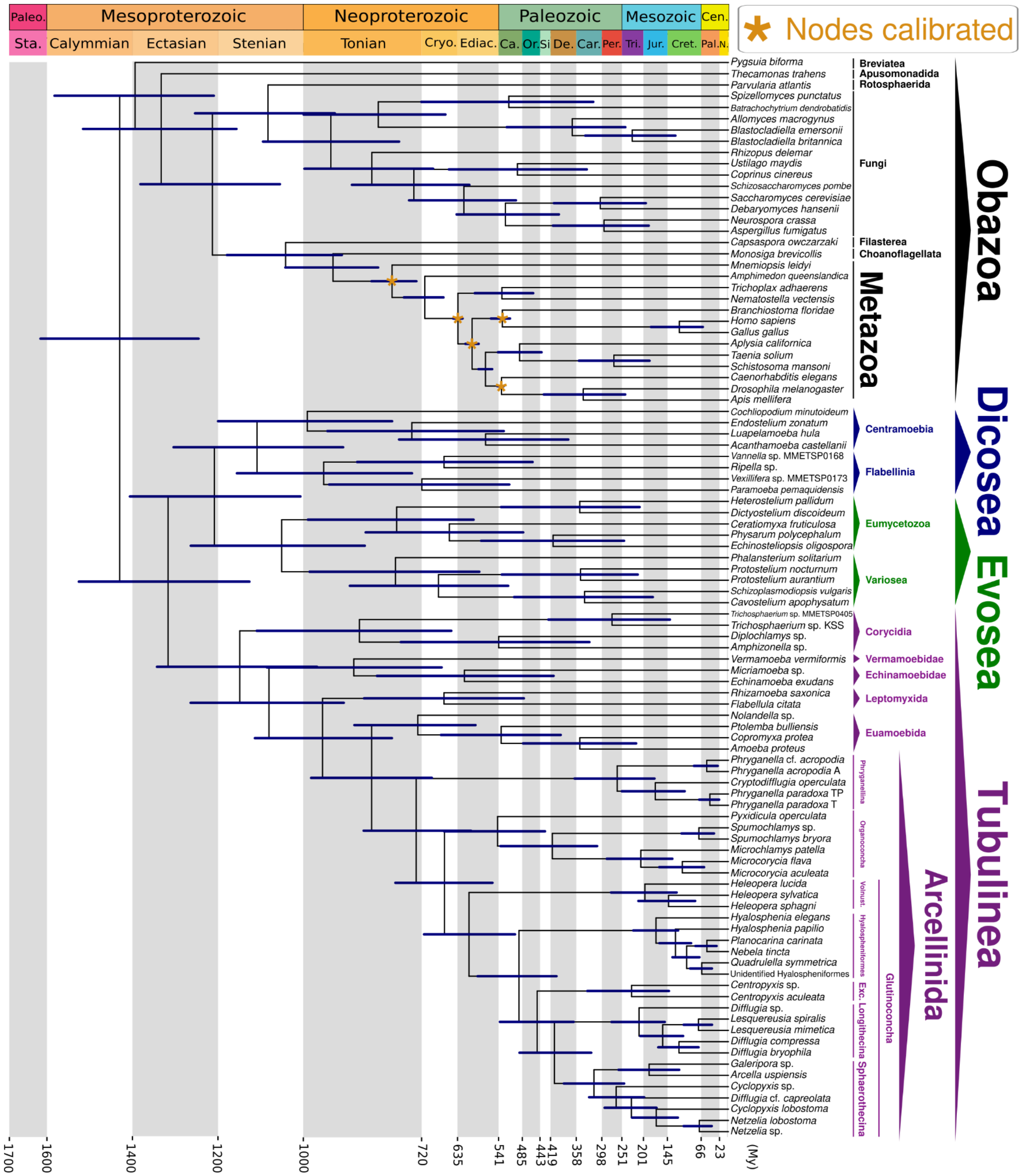


**Figure S7. Amorphea time-calibrated tree inferred under an uncorrelated relaxed clock model, applying a uniform distribution for node calibration and drift parameter of $\alpha$ = 2 and β = 2, calibrating nodes within the Metazoa group and excluding the VSM record to calibrate amoebozoan nodes (Dataset S01, Table S11).** Bars at nodes are 95% highest probability density confidence intervals. Abbreviations: Exc.- Excentrotoma; Vonust. - Volnustoma; Paleo. - Paleoproterozoic; Sta. - Statherian; Cryo. - Cryogenian; Ediac. - Ediacaran; Ca. - Cambrian; Or. Ordovician; Si - Silurian; De. - Devonian; Car. Carboniferous; Per. - Permian; Tri. - Triassic; Jur. - Jurassic; Cret. - Cretaceous; Cen. - Cenozoic; Pal. - Paleogene; N. - Neogene; My - Million Years.

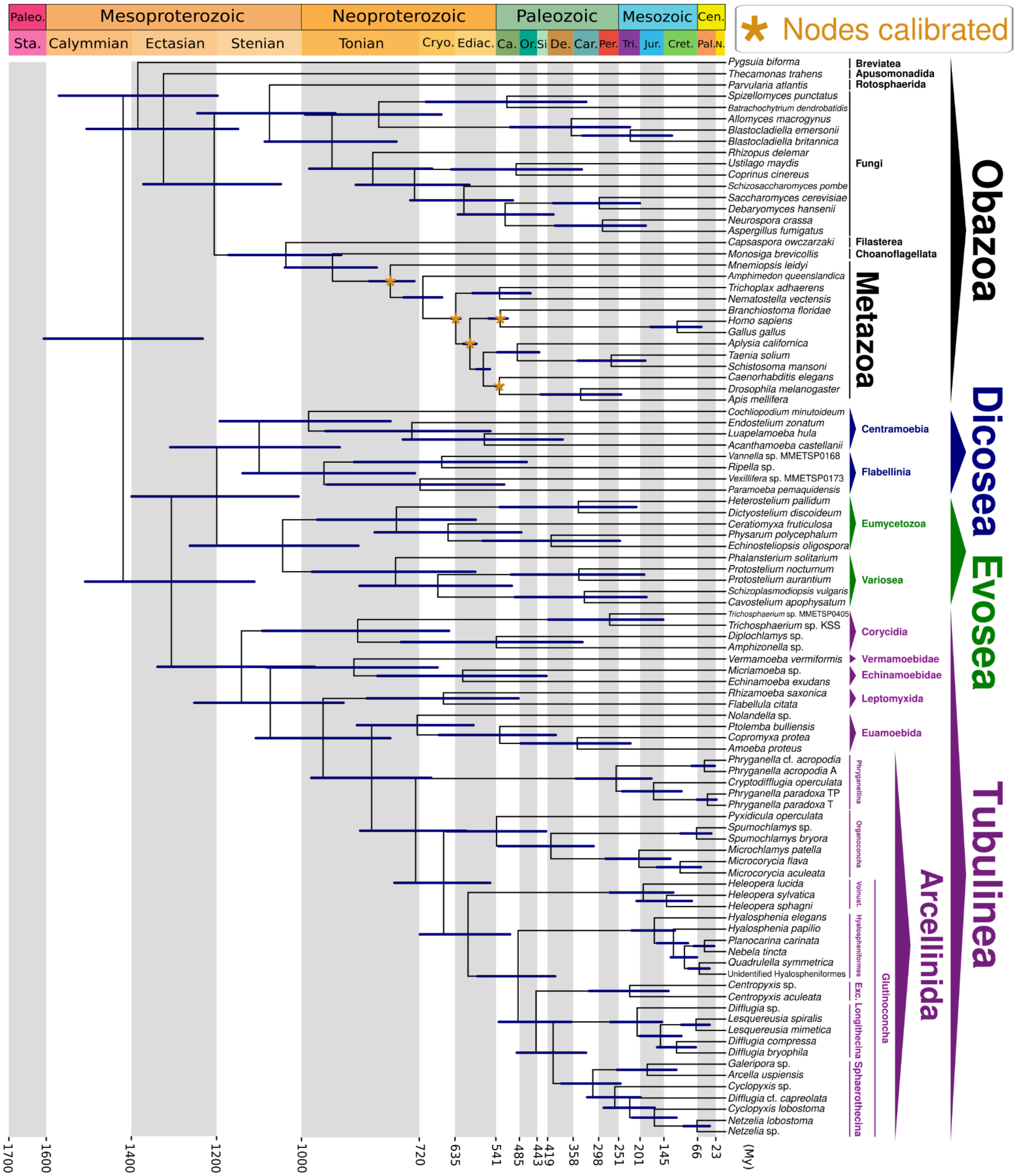


**Figure S8. Amorphea time-calibrated tree inferred under an uncorrelated relaxed clock model, applying a uniform distribution for node calibration and drift parameter of $\alpha$ = 1 and β = 10, calibrating nodes within the Metazoa group and excluding the VSM record to calibrate amoebozoan nodes (Dataset S01, Table S11).** Bars at nodes are 95% highest probability density confidence intervals. Abbreviations: Exc.- Excentrotoma; Vonust. - Volnustoma; Paleo. - Paleoproterozoic; Sta. - Statherian; Cryo. - Cryogenian; Ediac. - Ediacaran; Ca. - Cambrian; Or. Ordovician; Si - Silurian; De. - Devonian; Car. Carboniferous; Per. - Permian; Tri. - Triassic; Jur. - Jurassic; Cret. - Cretaceous; Cen. - Cenozoic; Pal. - Paleogene; N. - Neogene; My - Million Years.

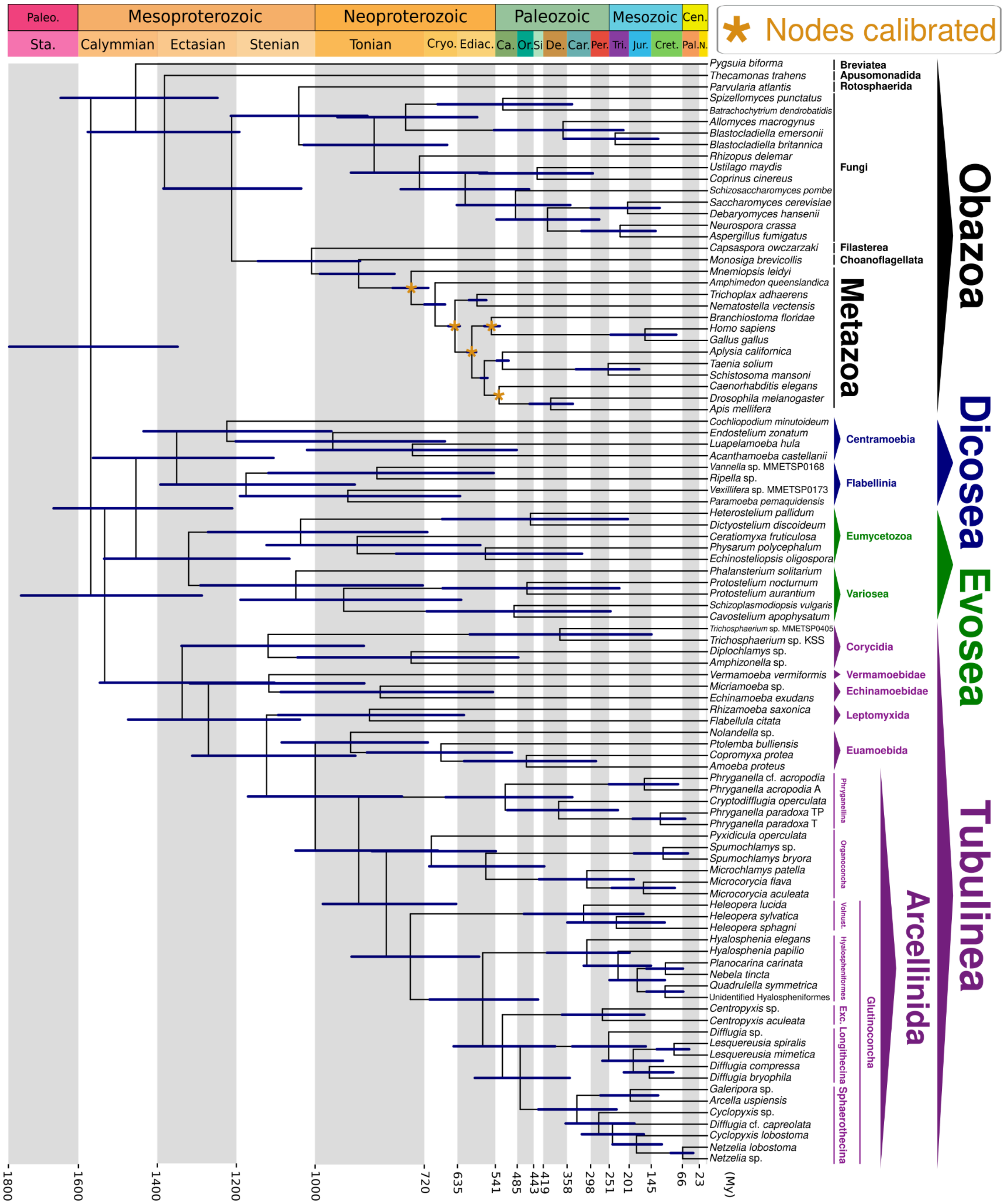

**Figure S9. Amorphea time-calibrated tree inferred under an autocorrelated relaxed clock model, applying a uniform distribution for node calibration and drift parameter of $\alpha$ = 2 and β = 2, calibrating nodes within the Metazoa group and excluding the VSM record to calibrate amoebozoan nodes (Dataset S01, Table S11).** Bars at nodes are 95% highest probability density confidence intervals. Abbreviations: Exc.- Excentrotoma; Vonust. - Volnustoma; Paleo. - Paleoproterozoic; Sta. - Statherian; Cryo. - Cryogenian; Ediac. - Ediacaran; Ca. - Cambrian; Or. Ordovician; Si - Silurian; De. - Devonian; Car. Carboniferous; Per. - Permian; Tri. - Triassic; Jur. - Jurassic; Cret. - Cretaceous; Cen. - Cenozoic; Pal. - Paleogene; N. - Neogene; My - Million Years.

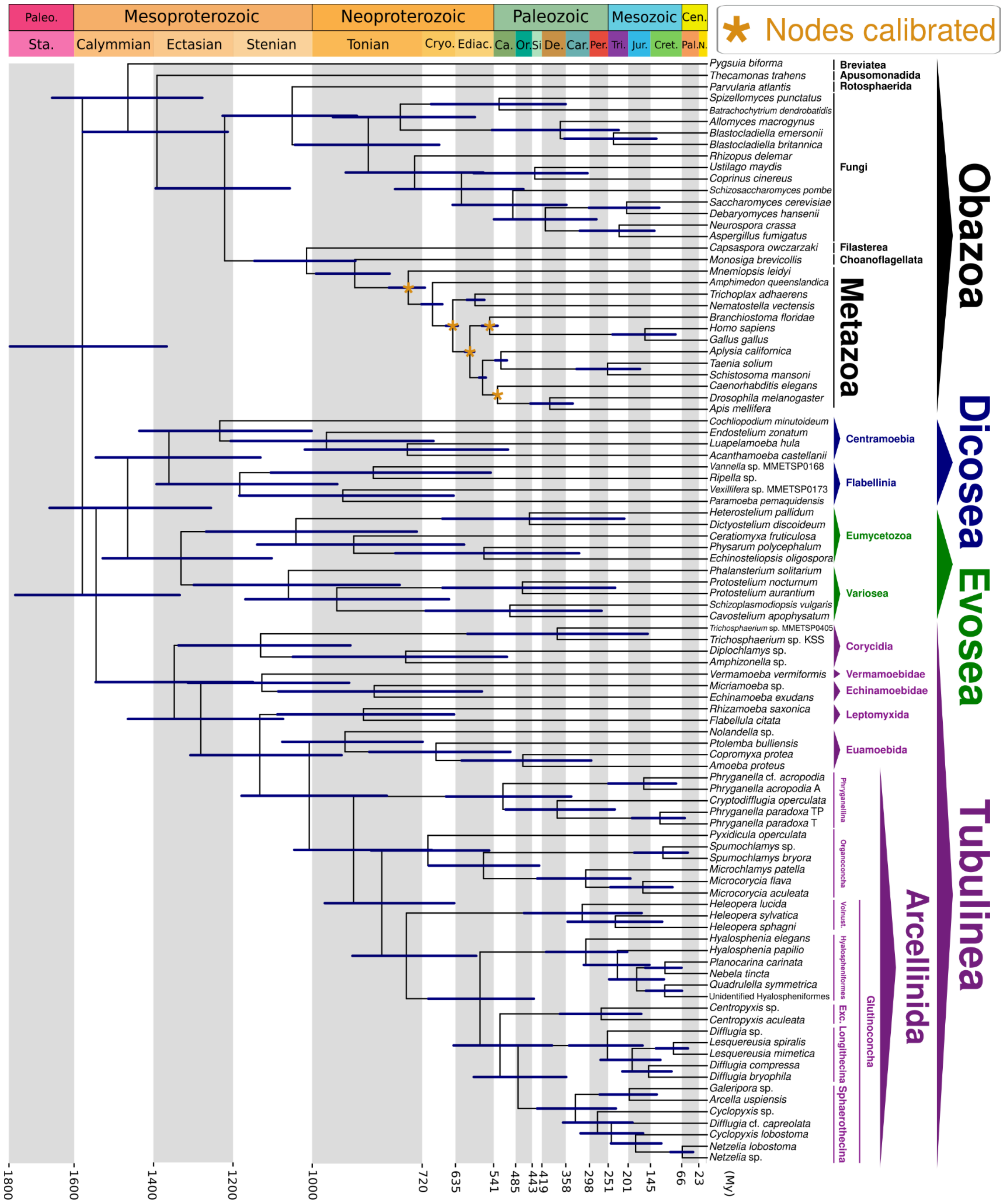


**Figure S10. Amorphea time-calibrated tree inferred under an autocorrelated relaxed clock model, applying a uniform distribution for node calibration and drift parameter of $\alpha$ = 1 and β = 10, calibrating nodes within the Metazoa group and excluding the VSM record to calibrate amoebozoan nodes (Dataset S01, Table S11).** Bars at nodes are 95% highest probability density confidence intervals. Abbreviations: Exc.- Excentrotoma; Vonust. - Volnustoma; Paleo. - Paleoproterozoic; Sta. - Statherian; Cryo. - Cryogenian; Ediac. - Ediacaran; Ca. - Cambrian; Or. Ordovician; Si - Silurian; De. - Devonian; Car. Carboniferous; Per. - Permian; Tri. - Triassic; Jur. - Jurassic; Cret. - Cretaceous; Cen. - Cenozoic; Pal. - Paleogene; N. - Neogene; My - Million Years.

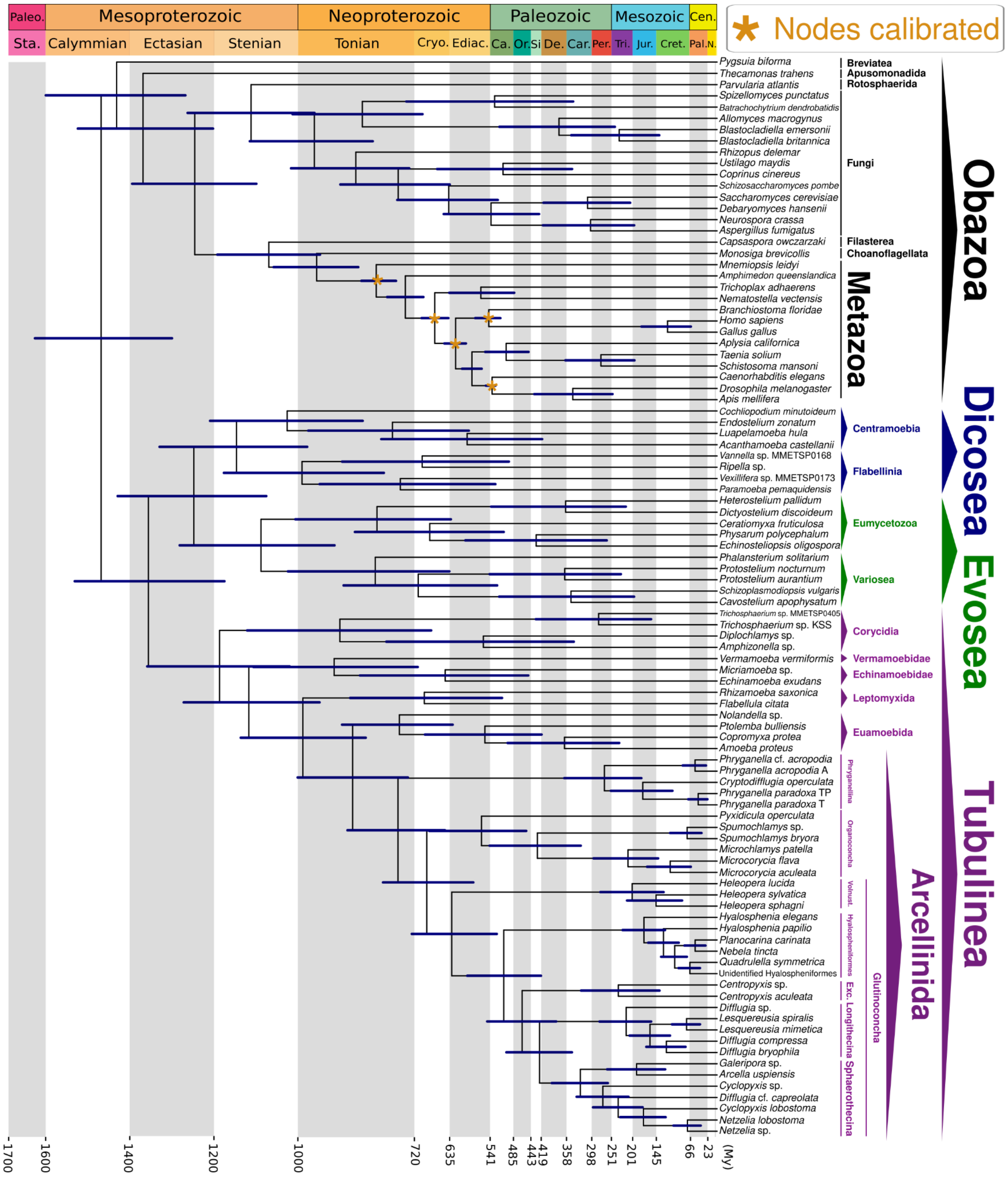


**Figure S11. Amorphea time-calibrated tree inferred under an uncorrelated relaxed clock model, applying a Skew-Normal distribution for node calibration and drift parameter of $\alpha$ = 2 and β = 2, calibrating nodes within the Metazoa group and excluding the VSM record to calibrate amoebozoan nodes (Dataset S01, Table S11).** Bars at nodes are 95% highest probability density confidence intervals. Abbreviations: Exc.- Excentrotoma; Vonust. - Volnustoma; Paleo. - Paleoproterozoic; Sta. - Statherian; Cryo. - Cryogenian; Ediac. - Ediacaran; Ca. - Cambrian; Or. Ordovician; Si - Silurian; De. - Devonian; Car. Carboniferous; Per. - Permian; Tri. - Triassic; Jur. - Jurassic; Cret. - Cretaceous; Cen. - Cenozoic; Pal. - Paleogene; N. - Neogene; My - Million Years.

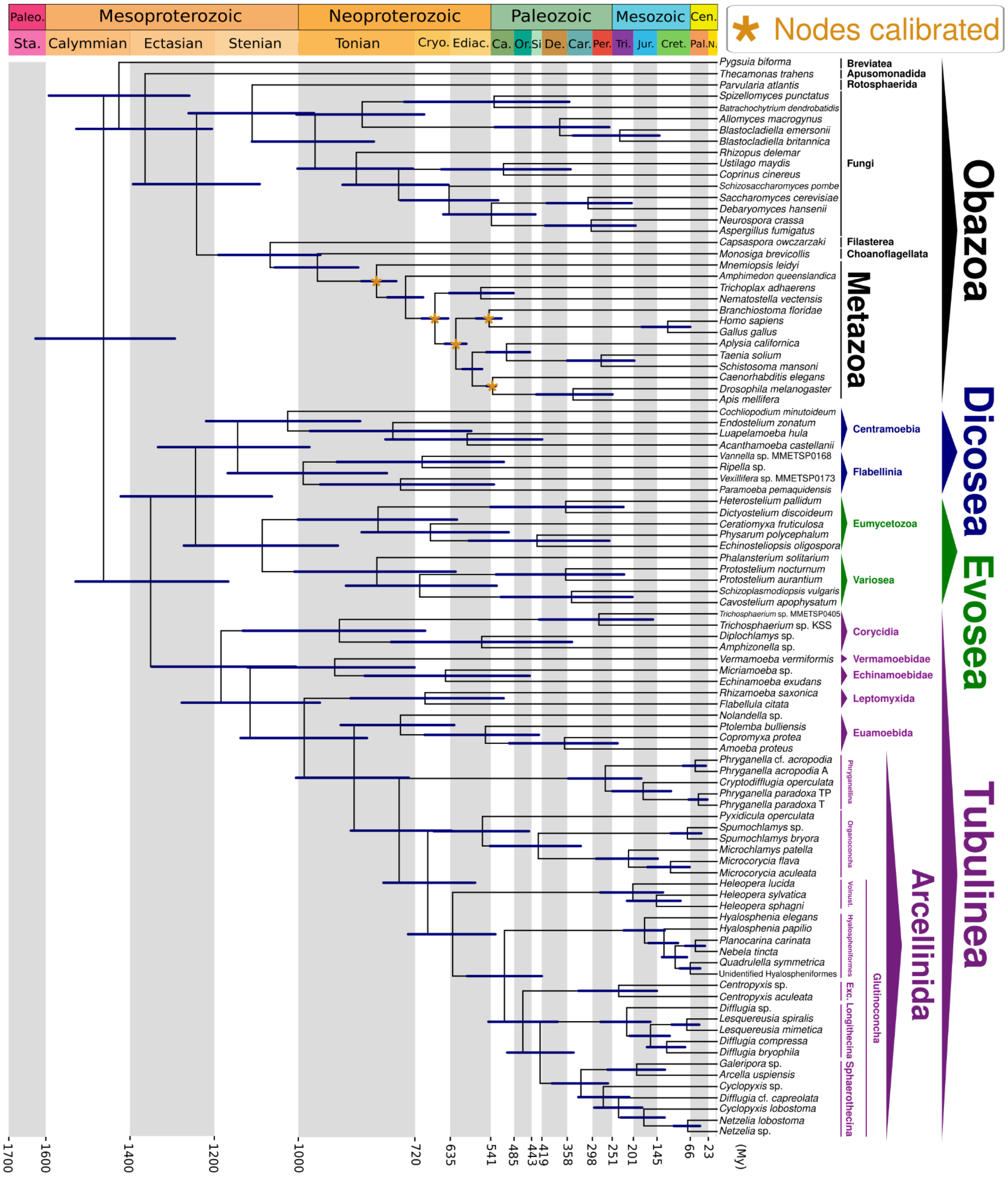


**Figure S12. Amorphea time-calibrated tree inferred under an uncorrelated relaxed clock model, applying a Skew-Normal distribution for node calibration and drift parameter of $\alpha$ = 1 and β = 10, calibrating nodes within the Metazoa group and excluding the VSM record to calibrate amoebozoan nodes (Dataset S01, Table S11).** Bars at nodes are 95% highest probability density confidence intervals. Abbreviations: Exc.- Excentrotoma; Vonust. - Volnustoma; Paleo. - Paleoproterozoic; Sta. - Statherian; Cryo. - Cryogenian; Ediac. - Ediacaran; Ca. - Cambrian; Or. Ordovician; Si - Silurian; De. - Devonian; Car. Carboniferous; Per. - Permian; Tri. - Triassic; Jur. - Jurassic; Cret. - Cretaceous; Cen. - Cenozoic; Pal. - Paleogene; N. - Neogene; My - Million Years.

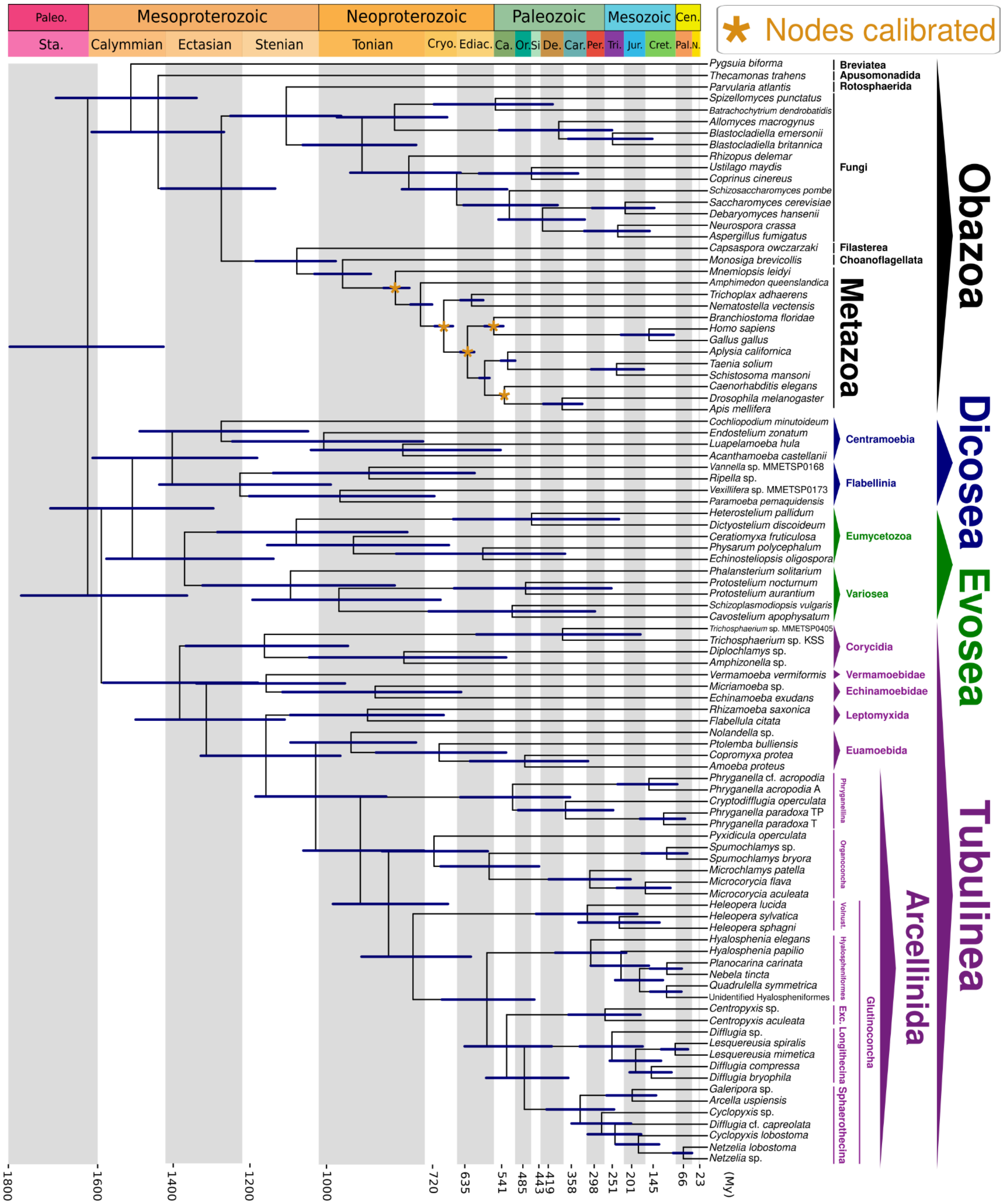


**Figure S13. Amorphea time-calibrated tree inferred under an autocorrelated relaxed clock model, applying a Skew-Normal distribution for node calibration and drift parameter of $\alpha$ = 2 and β = 2, calibrating nodes within the Metazoa group and excluding the VSM record to calibrate amoebozoan nodes (Dataset S01, Table S11).** Bars at nodes are 95% highest probability density confidence intervals. Abbreviations: Exc.- Excentrotoma; Vonust. - Volnustoma; Paleo. - Paleoproterozoic; Sta. - Statherian; Cryo. - Cryogenian; Ediac. - Ediacaran; Ca. - Cambrian; Or. Ordovician; Si - Silurian; De. - Devonian; Car. Carboniferous; Per. - Permian; Tri. - Triassic; Jur. - Jurassic; Cret. - Cretaceous; Cen. - Cenozoic; Pal. - Paleogene; N. - Neogene; My - Million Years.

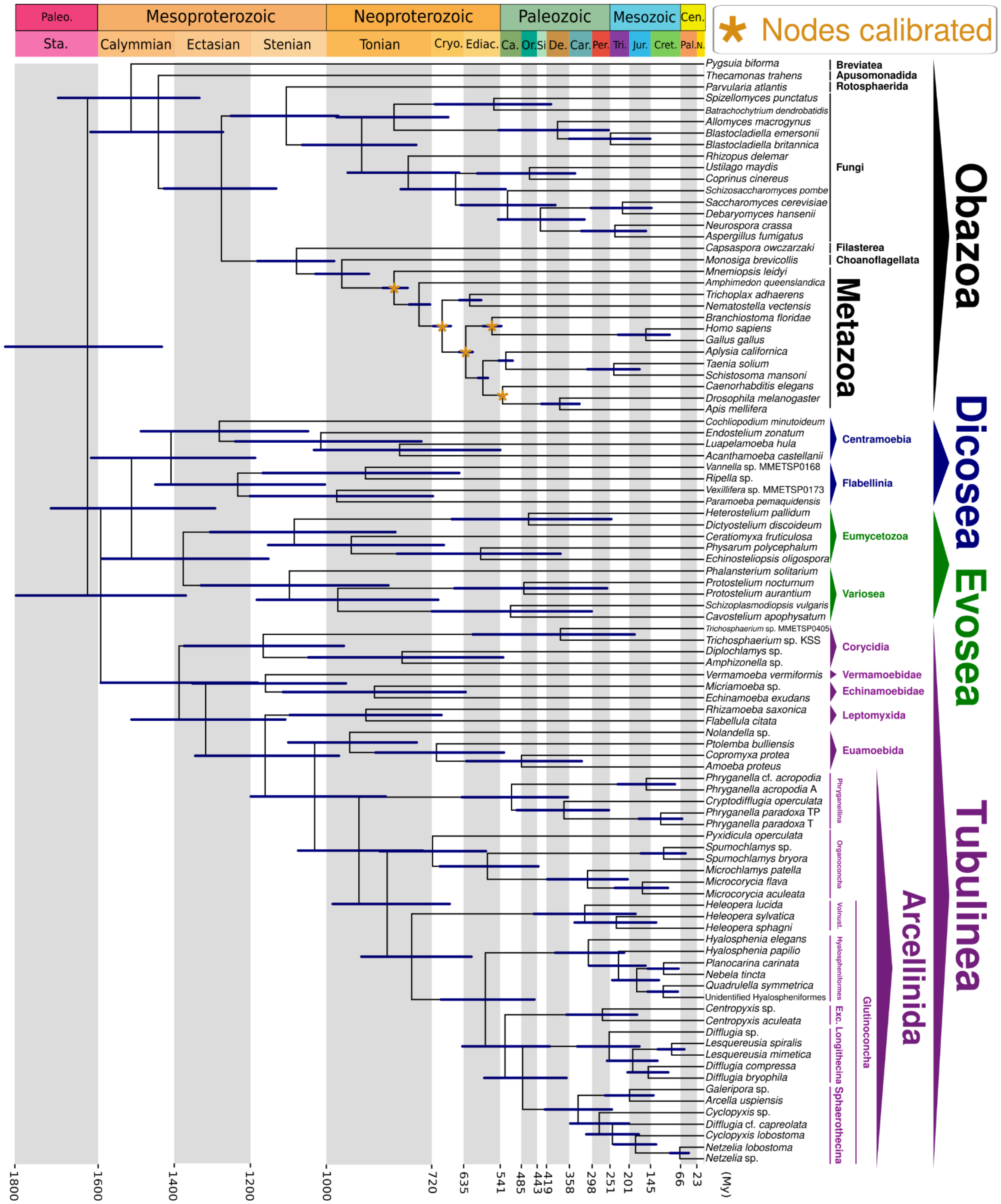

**Figure S14. Amorphea time-calibrated tree inferred under an autocorrelated relaxed clock model, applying a Skew-Normal distribution for node calibration and drift parameter of $\alpha$ = 1 and β = 10, calibrating nodes within the Metazoa group and excluding the VSM record to calibrate amoebozoan nodes (Dataset S01, Table S11).** Bars at nodes are 95% highest probability density confidence intervals. Abbreviations: Exc.- Excentrotoma; Vonust. - Volnustoma; Paleo. - Paleoproterozoic; Sta. - Statherian; Cryo. - Cryogenian; Ediac. - Ediacaran; Ca. - Cambrian; Or. Ordovician; Si - Silurian; De. - Devonian; Car. Carboniferous; Per. - Permian; Tri. - Triassic; Jur. - Jurassic; Cret. - Cretaceous; Cen. - Cenozoic; Pal. - Paleogene; N. - Neogene; My - Million Years.

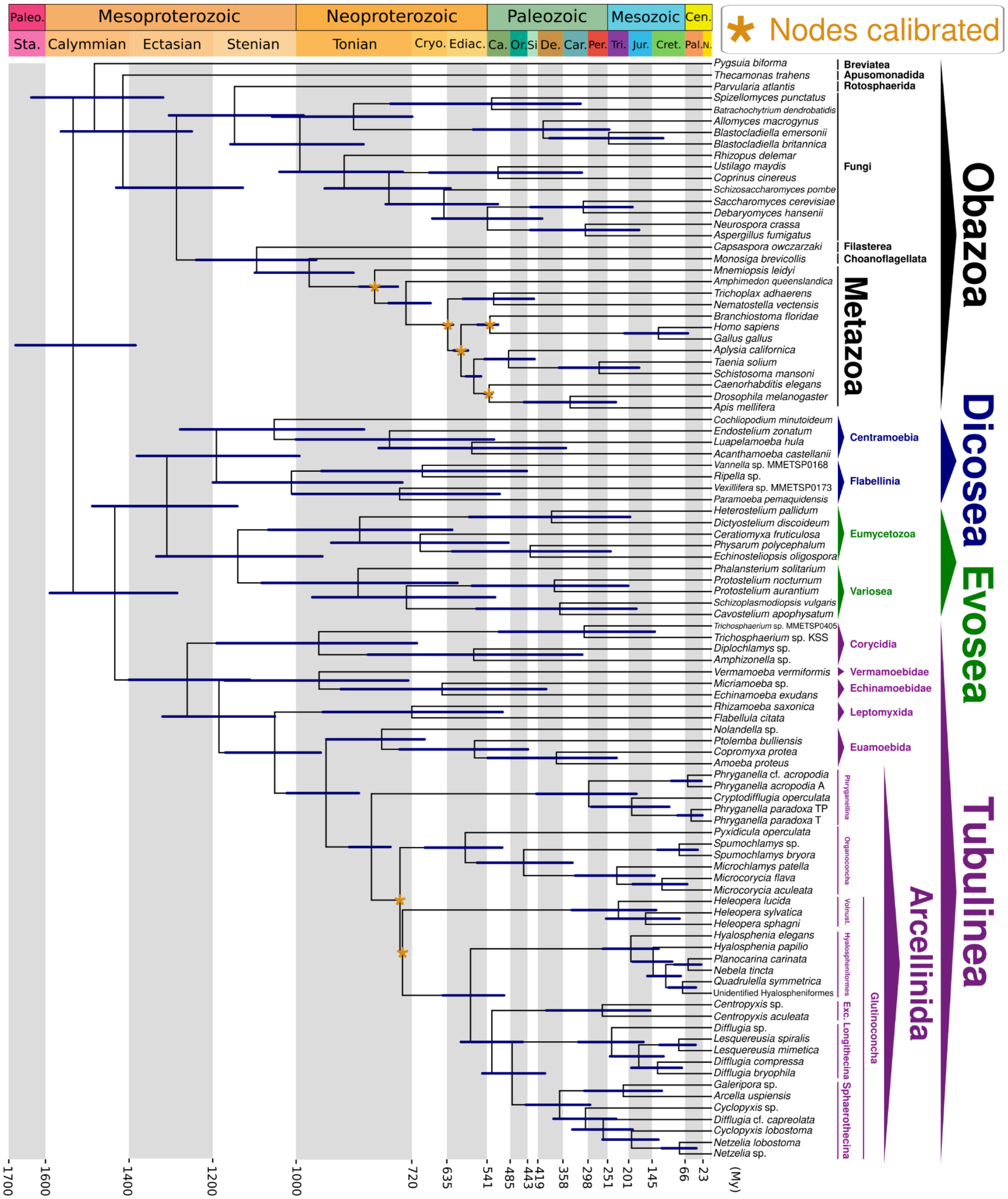


**Figure S15. Amorphea time-calibrated tree inferred under an uncorrelated relaxed clock model, applying a uniform distribution for node calibration and drift parameter of $\alpha$ = 2 and β = 2, calibrating nodes within the Metazoa group, as well as the Glutinoconcha+Organoconcha and Glutinoconcha nodes (Dataset S01, Table S11).** Bars at nodes are 95% highest probability density confidence intervals. Abbreviations: Exc.- Excentrotoma; Vonust. - Volnustoma; Paleo. - Paleoproterozoic; Sta. - Statherian; Cryo. - Cryogenian; Ediac. - Ediacaran; Ca. - Cambrian; Or. Ordovician; Si - Silurian; De. - Devonian; Car. Carboniferous; Per. - Permian; Tri. - Triassic; Jur. - Jurassic; Cret. - Cretaceous; Cen. - Cenozoic; Pal. - Paleogene; N. - Neogene; My - Million Years.

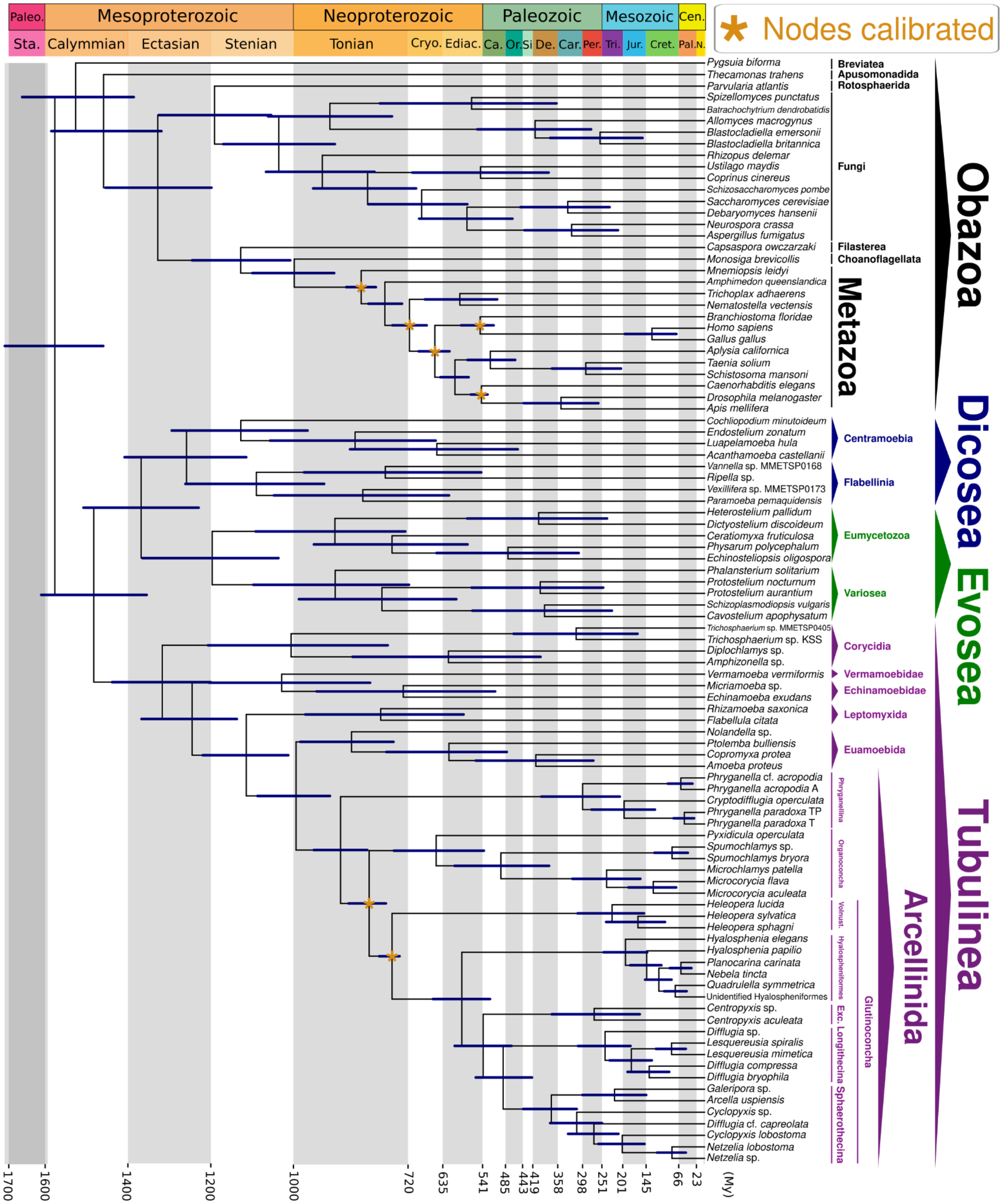

**Figure S16. Amorphea time-calibrated tree inferred under an uncorrelated relaxed clock model, applying a uniform distribution for node calibration and drift parameter of $\alpha$ = 1 and β = 10, calibrating nodes within the Metazoa group, as well as the Glutinoconcha+Organoconcha and Glutinoconcha nodes (Dataset S01, Table S11).** Bars at nodes are 95% highest probability density confidence intervals. Abbreviations: Exc.- Excentrotoma; Vonust. - Volnustoma; Paleo. - Paleoproterozoic; Sta. - Statherian; Cryo. - Cryogenian; Ediac. - Ediacaran; Ca. - Cambrian; Or. Ordovician; Si - Silurian; De. - Devonian; Car. Carboniferous; Per. - Permian; Tri. - Triassic; Jur. - Jurassic; Cret. - Cretaceous; Cen. - Cenozoic; Pal. - Paleogene; N. - Neogene; My - Million Years.

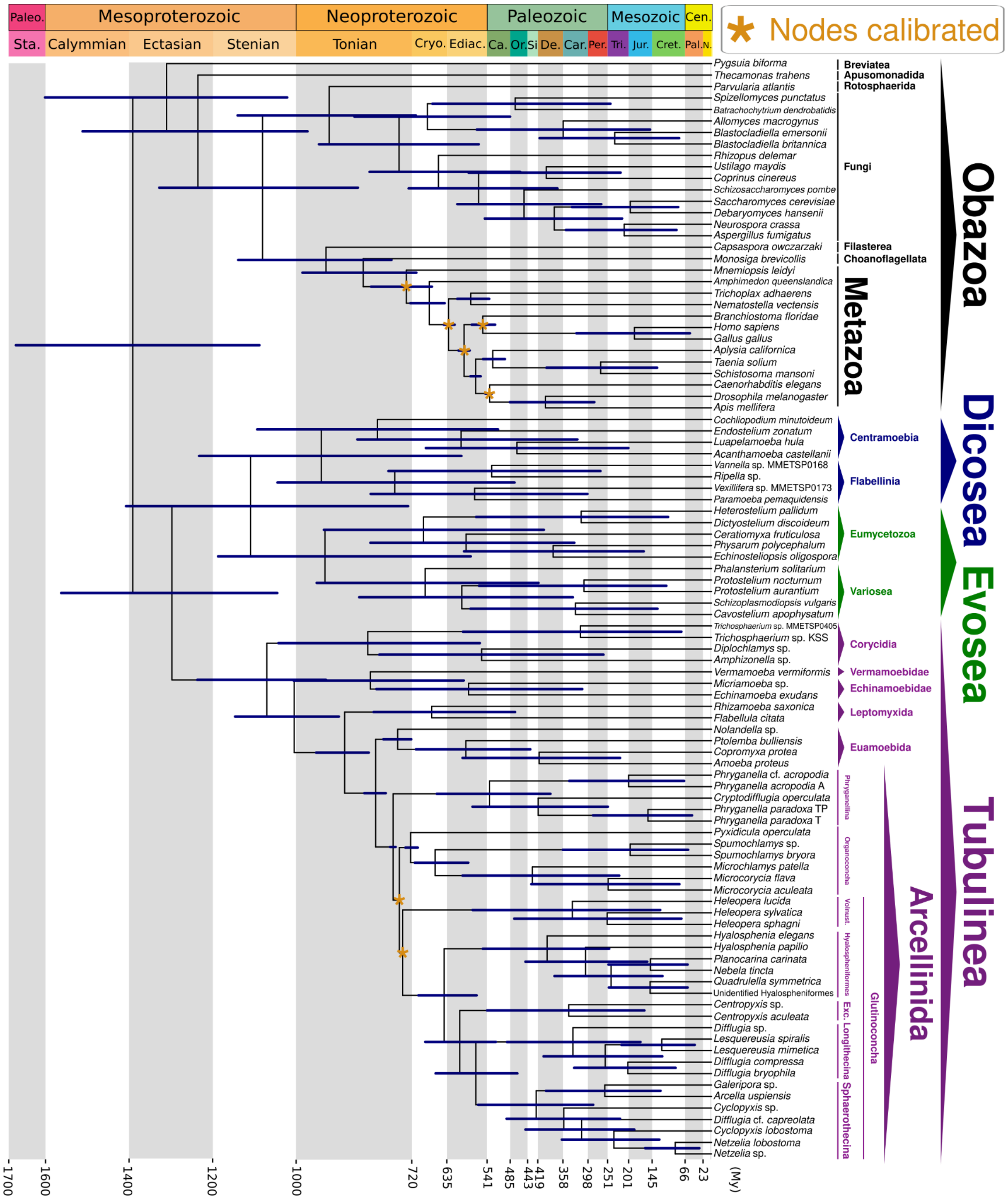


**Figure S17. Amorphea time-calibrated tree inferred under an autocorrelated relaxed clock model, applying a uniform distribution for node calibration and drift parameter of $\alpha$ = 2 and β = 2, calibrating nodes within the Metazoa group, as well as the Glutinoconcha+Organoconcha and Glutinoconcha nodes (Dataset S01, Table S11).** Bars at nodes are 95% highest probability density confidence intervals. Abbreviations: Exc.- Excentrotoma; Vonust. - Volnustoma; Paleo. - Paleoproterozoic; Sta. - Statherian; Cryo. - Cryogenian; Ediac. - Ediacaran; Ca. - Cambrian; Or. Ordovician; Si - Silurian; De. - Devonian; Car. Carboniferous; Per. - Permian; Tri. - Triassic; Jur. - Jurassic; Cret. - Cretaceous; Cen. - Cenozoic; Pal. - Paleogene; N. - Neogene; My - Million Years.

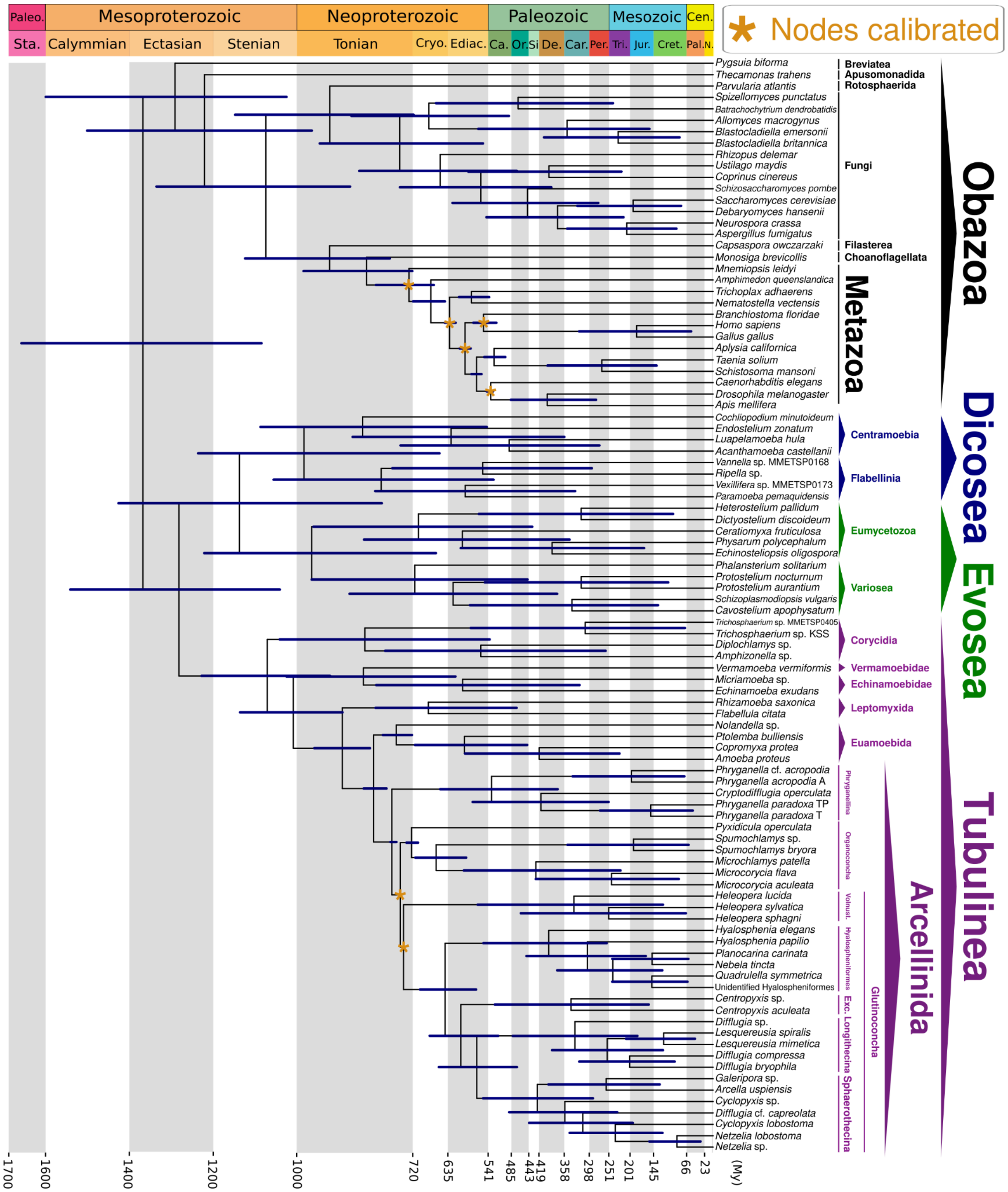


**Figure S18. Amorphea time-calibrated tree inferred under an autocorrelated relaxed clock model, applying a uniform distribution for node calibration and drift parameter of $\alpha$ = 1 and β = 10, calibrating nodes within the Metazoa group, as well as the Glutinoconcha+Organoconcha and Glutinoconcha nodes (Dataset S01, Table S11).** Bars at nodes are 95% highest probability density confidence intervals. Abbreviations: Exc.- Excentrotoma; Vonust. - Volnustoma; Paleo. - Paleoproterozoic; Sta. - Statherian; Cryo. - Cryogenian; Ediac. - Ediacaran; Ca. - Cambrian; Or. Ordovician; Si - Silurian; De. - Devonian; Car. Carboniferous; Per. - Permian; Tri. - Triassic; Jur. - Jurassic; Cret. - Cretaceous; Cen. - Cenozoic; Pal. - Paleogene; N. - Neogene; My - Million Years.

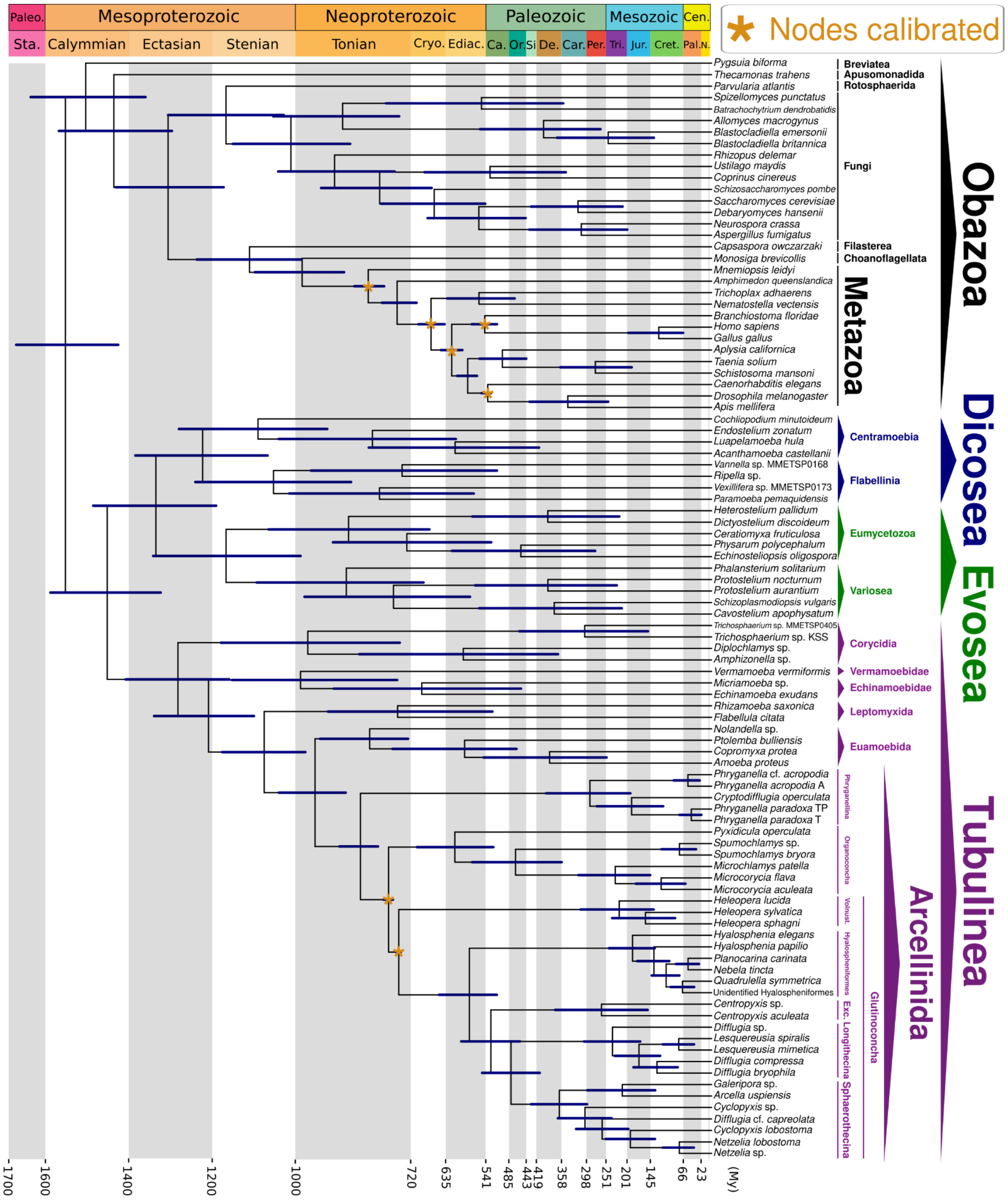


**Figure S19. Amorphea time-calibrated tree inferred under an uncorrelated relaxed clock model, applying a skew-normal distribution for node calibration and drift parameter of $\alpha$ = 2 and β = 2, calibrating nodes within the Metazoa group, as well as the Glutinoconcha+Organoconcha and Glutinoconcha nodes (Dataset S01, Table S11).** Bars at nodes are 95% highest probability density confidence intervals. Abbreviations: Exc.- Excentrotoma; Vonust. - Volnustoma; Paleo. - Paleoproterozoic; Sta. - Statherian; Cryo. - Cryogenian; Ediac. - Ediacaran; Ca. - Cambrian; Or. Ordovician; Si - Silurian; De. - Devonian; Car. Carboniferous; Per. - Permian; Tri. - Triassic; Jur. - Jurassic; Cret. - Cretaceous; Cen. - Cenozoic; Pal. - Paleogene; N. - Neogene; My - Million Years.

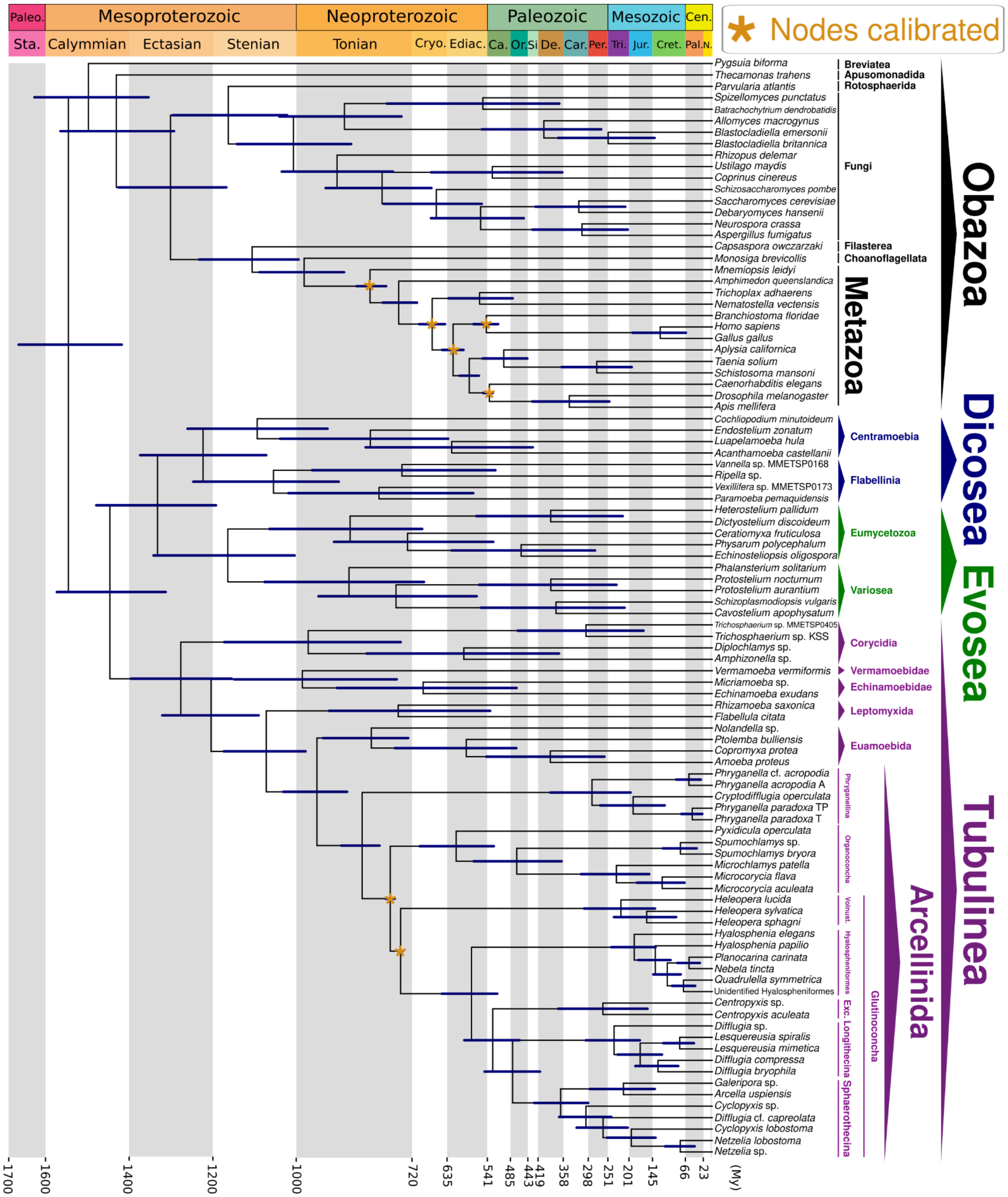

**Figure S20. Amorphea time-calibrated tree inferred under an uncorrelated relaxed clock model, applying a skew-normal distribution for node calibration and drift parameter of $\alpha$ = 1 and β = 10, calibrating nodes within the Metazoa group, as well as the Glutinoconcha+Organoconcha and Glutinoconcha nodes (Dataset S01, Table S11).** Bars at nodes are 95% highest probability density confidence intervals. Abbreviations: Exc.- Excentrotoma; Vonust. - Volnustoma; Paleo. - Paleoproterozoic; Sta. - Statherian; Cryo. - Cryogenian; Ediac. - Ediacaran; Ca. - Cambrian; Or. Ordovician; Si - Silurian; De. - Devonian; Car. Carboniferous; Per. - Permian; Tri. - Triassic; Jur. - Jurassic; Cret. - Cretaceous; Cen. - Cenozoic; Pal. - Paleogene; N. - Neogene; My - Million Years.

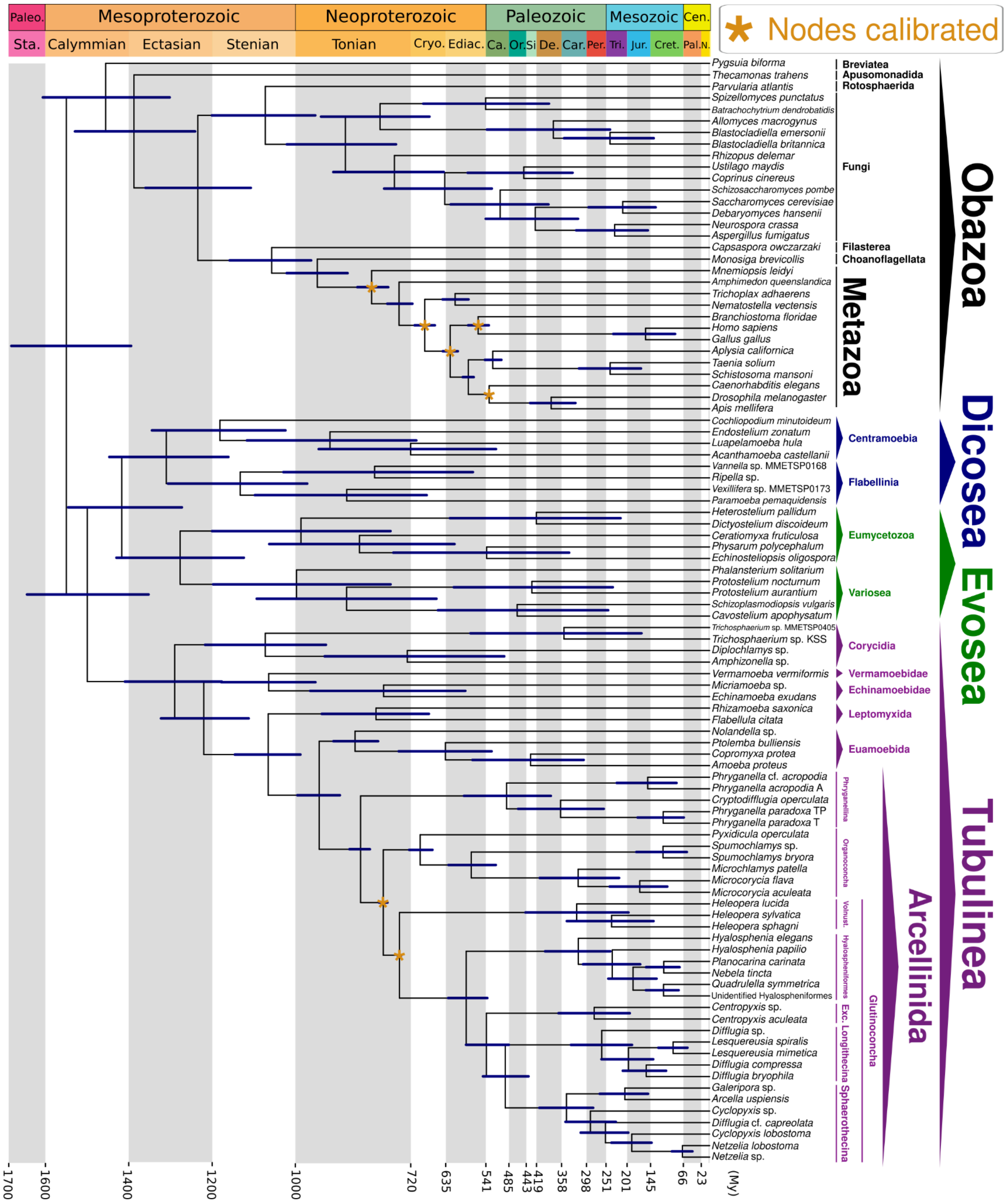


**Figure S21. Amorphea time-calibrated tree inferred under an autocorrelated relaxed clock model, applying a skew-normal distribution for node calibration and drift parameter of $\alpha$ = 2 and β = 2, calibrating nodes within the Metazoa group, as well as the Glutinoconcha+Organoconcha and Glutinoconcha nodes (Dataset S01, Table S11).** Bars at nodes are 95% highest probability density confidence intervals. Abbreviations: Exc.- Excentrotoma; Vonust. - Volnustoma; Paleo. - Paleoproterozoic; Sta. - Statherian; Cryo. - Cryogenian; Ediac. - Ediacaran; Ca. - Cambrian; Or. Ordovician; Si - Silurian; De. - Devonian; Car. Carboniferous; Per. - Permian; Tri. - Triassic; Jur. - Jurassic; Cret. - Cretaceous; Cen. - Cenozoic; Pal. - Paleogene; N. - Neogene; My - Million Years.

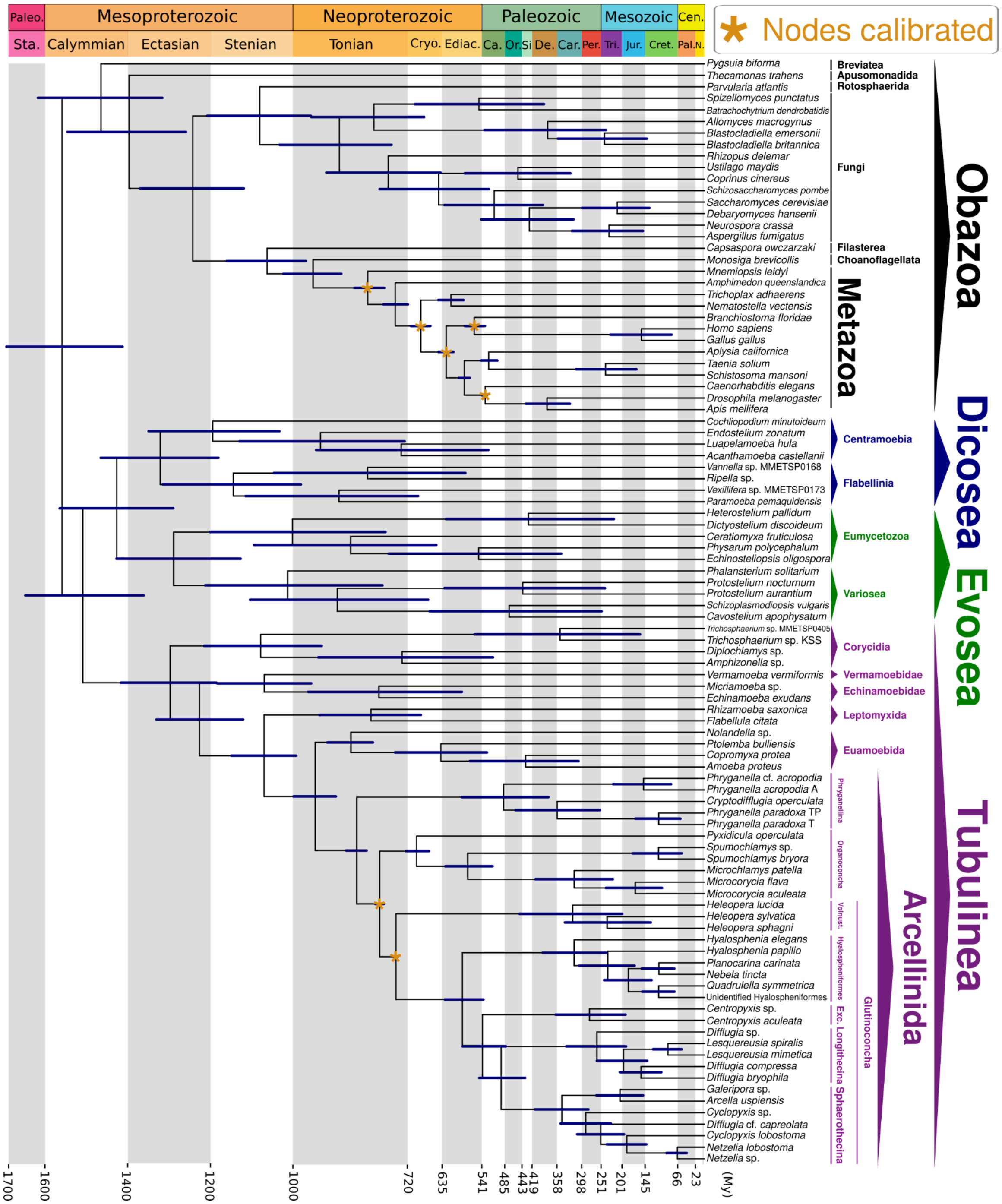


**Figure S22. Amorphea time-calibrated tree inferred under an autocorrelated relaxed clock model, applying a skew-normal distribution for node calibration and drift parameter of $\alpha$ = 1 and β = 10, calibrating nodes within the Metazoa group, as well as the Glutinoconcha+Organoconcha and Glutinoconcha nodes (Dataset S01, Table S11).** Bars at nodes are 95% highest probability density confidence intervals. Abbreviations: Exc.- Excentrotoma; Vonust. - Volnustoma; Paleo. - Paleoproterozoic; Sta. - Statherian; Cryo. - Cryogenian; Ediac. - Ediacaran; Ca. - Cambrian; Or. Ordovician; Si - Silurian; De. - Devonian; Car. Carboniferous; Per. - Permian; Tri. - Triassic; Jur. - Jurassic; Cret. - Cretaceous; Cen. - Cenozoic; Pal. - Paleogene; N. - Neogene; My - Million Years.

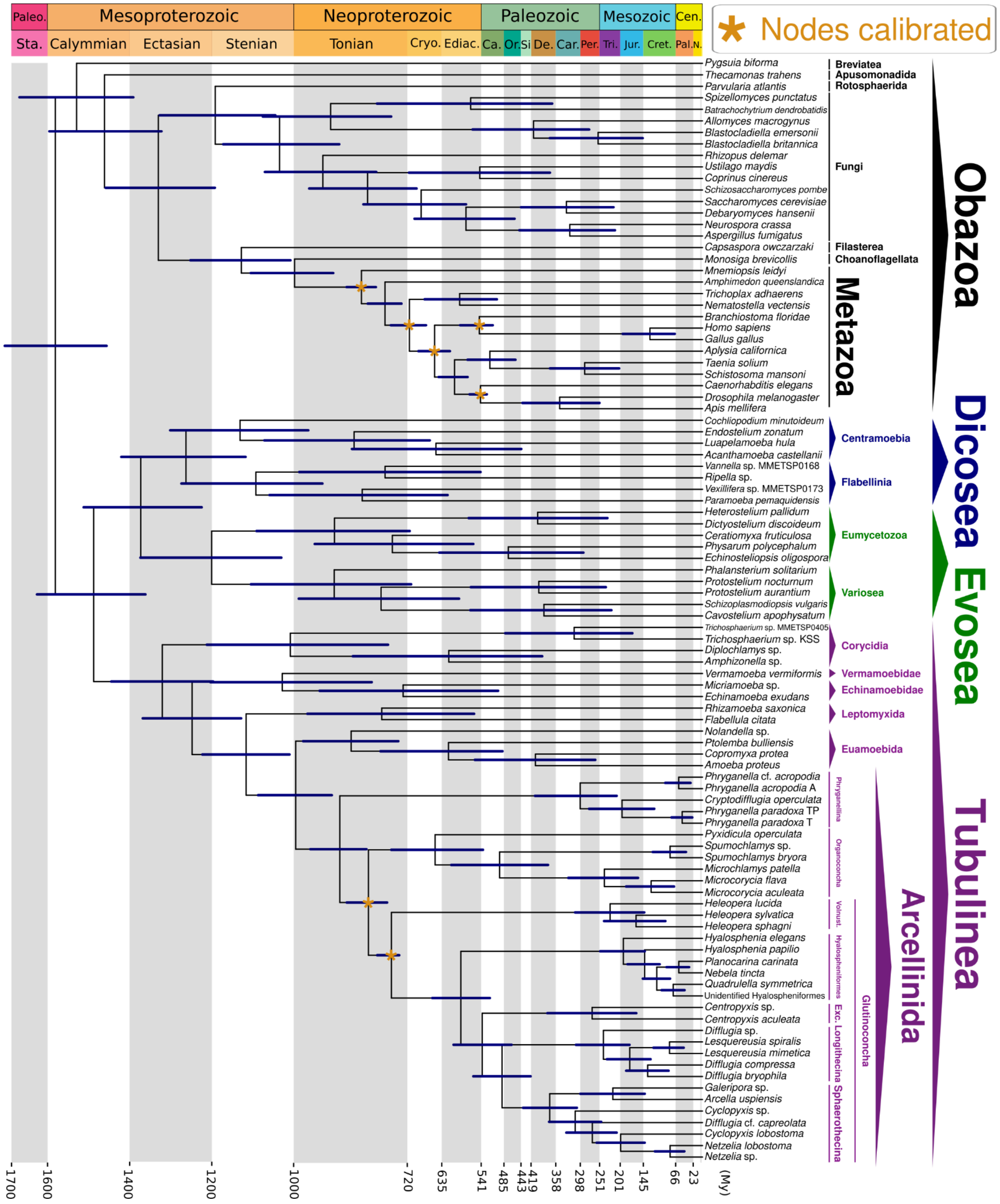


**Figure S23. Amorphea time-calibrated tree inferred under an uncorrelated relaxed clock model, applying a truncated-Cauchy short-tail distribution for node calibration and drift parameter of $\alpha$ = 2 and β = 2, calibrating nodes within the Metazoa group, as well as the Glutinoconcha+Organoconcha and Glutinoconcha nodes (Dataset S01, Table S11).** Bars at nodes are 95% highest probability density confidence intervals. Abbreviations: Exc.- Excentrotoma; Vonust. - Volnustoma; Paleo. - Paleoproterozoic; Sta. - Statherian; Cryo. - Cryogenian; Ediac. - Ediacaran; Ca. - Cambrian; Or. Ordovician; Si - Silurian; De. - Devonian; Car. Carboniferous; Per. - Permian; Tri. - Triassic; Jur. - Jurassic; Cret. - Cretaceous; Cen. - Cenozoic; Pal. - Paleogene; N. - Neogene; My - Million Years.

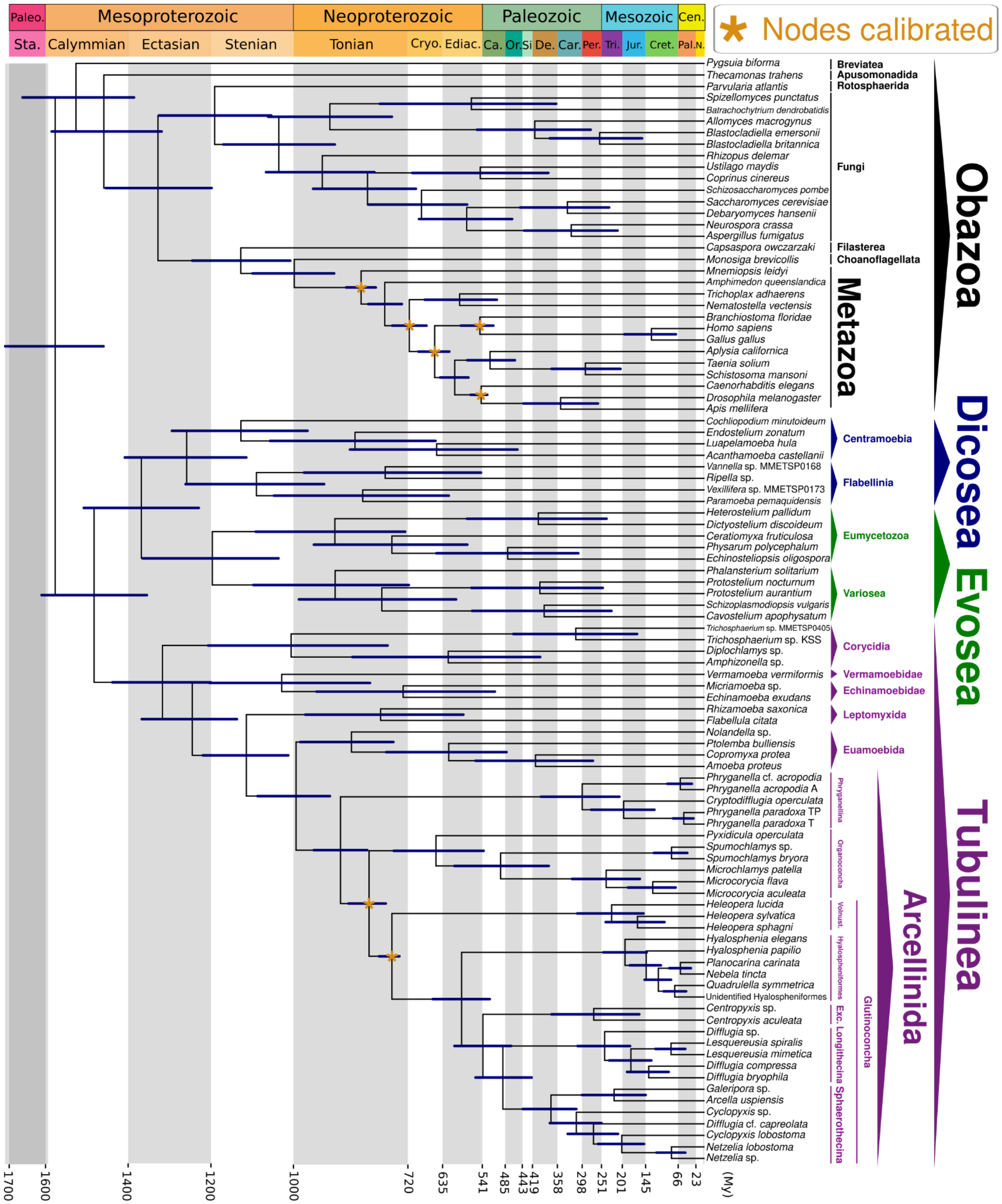


**Figure S24. Amorphea time-calibrated tree inferred under an uncorrelated relaxed clock model, applying a truncated-Cauchy short-tail distribution for node calibration and drift parameter of $\alpha$ = 1 and β = 10, calibrating nodes within the Metazoa group, as well as the Glutinoconcha+Organoconcha and Glutinoconcha nodes (Dataset S01, Table S11).** Bars at nodes are 95% highest probability density confidence intervals. Abbreviations: Exc.- Excentrotoma; Vonust. - Volnustoma; Paleo. - Paleoproterozoic; Sta. - Statherian; Cryo. - Cryogenian; Ediac. - Ediacaran; Ca. - Cambrian; Or. Ordovician; Si - Silurian; De. - Devonian; Car. Carboniferous; Per. - Permian; Tri. - Triassic; Jur. - Jurassic; Cret. - Cretaceous; Cen. - Cenozoic; Pal. - Paleogene; N. - Neogene; My - Million Years.

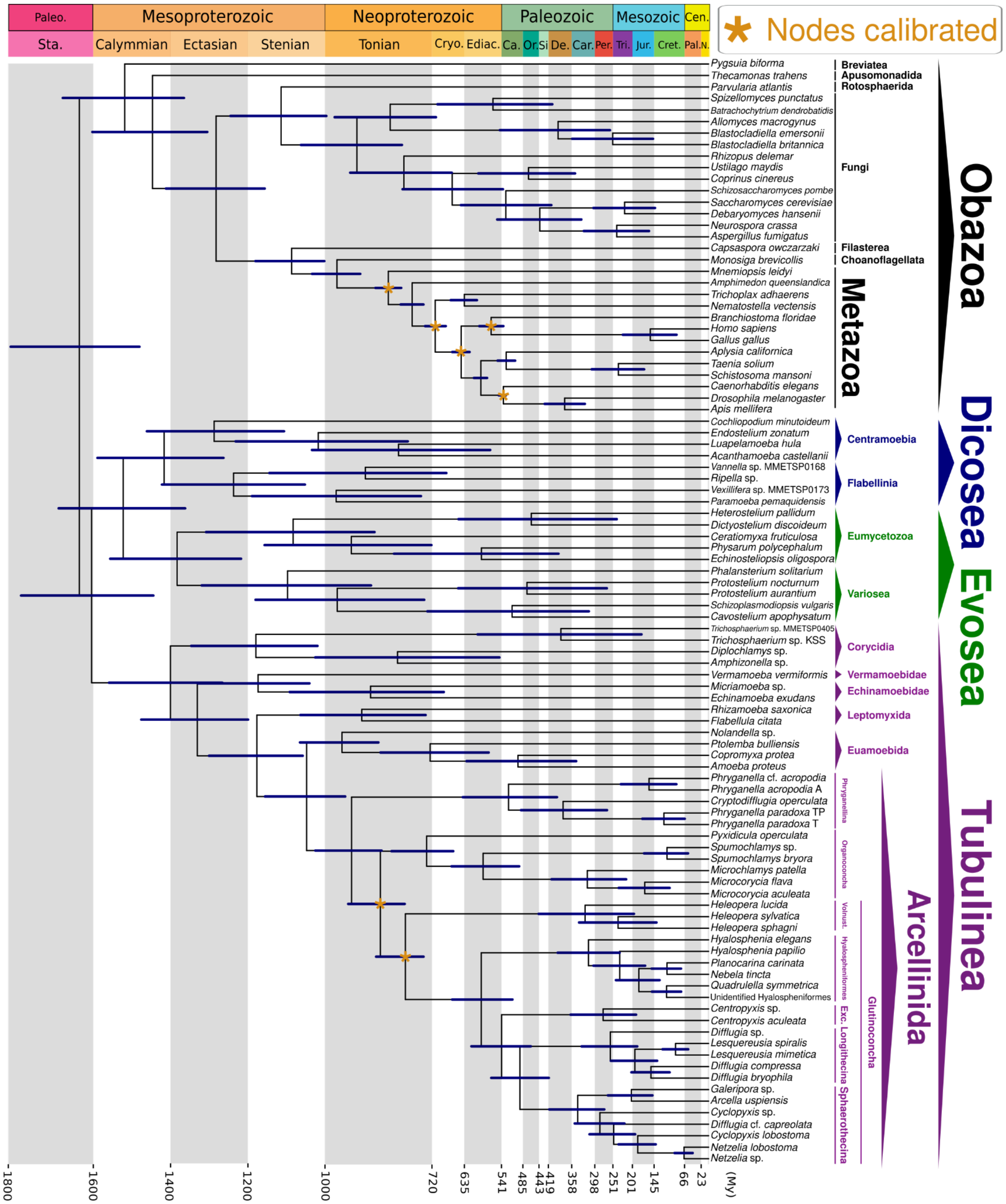


**Figure S25. Amorphea time-calibrated tree inferred under an autocorrelated relaxed clock model, applying a truncated-Cauchy short-tail distribution for node calibration and drift parameter of $\alpha$ = 2 and β = 2, calibrating nodes within the Metazoa group, as well as the Glutinoconcha+Organoconcha and Glutinoconcha nodes (Dataset S01, Table S11).** Bars at nodes are 95% highest probability density confidence intervals. Abbreviations: Exc.- Excentrotoma; Vonust. - Volnustoma; Paleo. - Paleoproterozoic; Sta. - Statherian; Cryo. - Cryogenian; Ediac. - Ediacaran; Ca. - Cambrian; Or. Ordovician; Si - Silurian; De. - Devonian; Car. Carboniferous; Per. - Permian; Tri. - Triassic; Jur. - Jurassic; Cret. - Cretaceous; Cen. - Cenozoic; Pal. - Paleogene; N. - Neogene; My - Million Years.

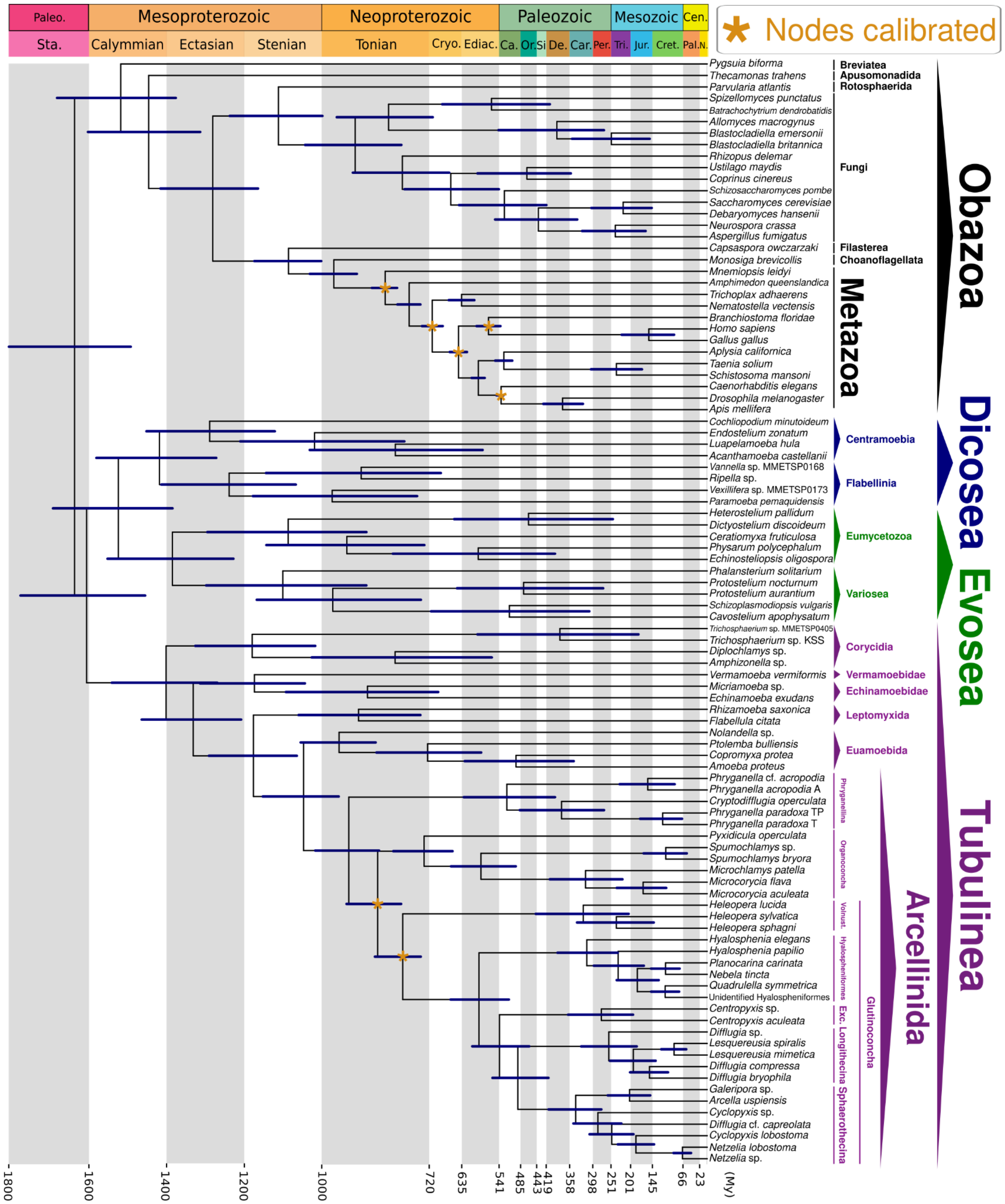


**Figure S26. Amorphea time-calibrated tree inferred under an autocorrelated relaxed clock model, applying a truncated-Cauchy short-tail distribution for node calibration and drift parameter of $\alpha$ = 1 and β = 10, calibrating nodes within the Metazoa group, as well as the Glutinoconcha+Organoconcha and Glutinoconcha nodes (Dataset S01, Table S11).** Bars at nodes are 95% highest probability density confidence intervals. Abbreviations: Exc.- Excentrotoma; Vonust. - Volnustoma; Paleo. - Paleoproterozoic; Sta. - Statherian; Cryo. - Cryogenian; Ediac. - Ediacaran; Ca. - Cambrian; Or. Ordovician; Si - Silurian; De. - Devonian; Car. Carboniferous; Per. - Permian; Tri. - Triassic; Jur. - Jurassic; Cret. - Cretaceous; Cen. - Cenozoic; Pal. - Paleogene; N. - Neogene; My - Million Years.

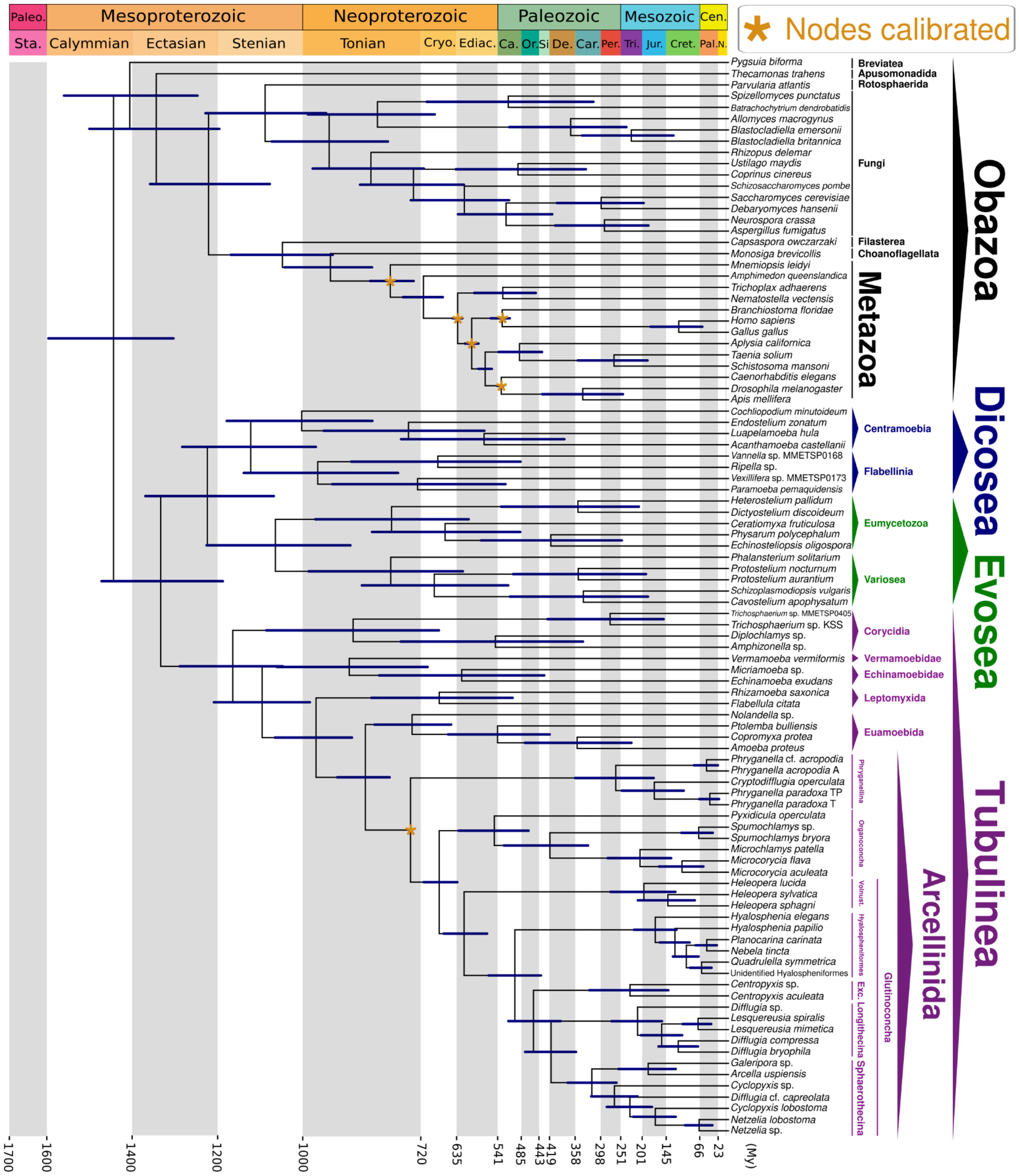


**Figure S27. Amorphea time-calibrated tree inferred under an uncorrelated relaxed clock model, applying a uniform distribution for node calibration and drift parameter of $\alpha$ = 2 and β = 2, calibrating nodes within the Metazoa group, as well as the Arcellinida node (Dataset S01, Table S11).** Bars at nodes are 95% highest probability density confidence intervals. Abbreviations: Exc.- Excentrotoma; Vonust. - Volnustoma; Paleo. - Paleoproterozoic; Sta. - Statherian; Cryo. - Cryogenian; Ediac. - Ediacaran; Ca. - Cambrian; Or. Ordovician; Si - Silurian; De. - Devonian; Car. Carboniferous; Per. - Permian; Tri. - Triassic; Jur. - Jurassic; Cret. - Cretaceous; Cen. - Cenozoic; Pal. - Paleogene; N. - Neogene; My - Million Years.

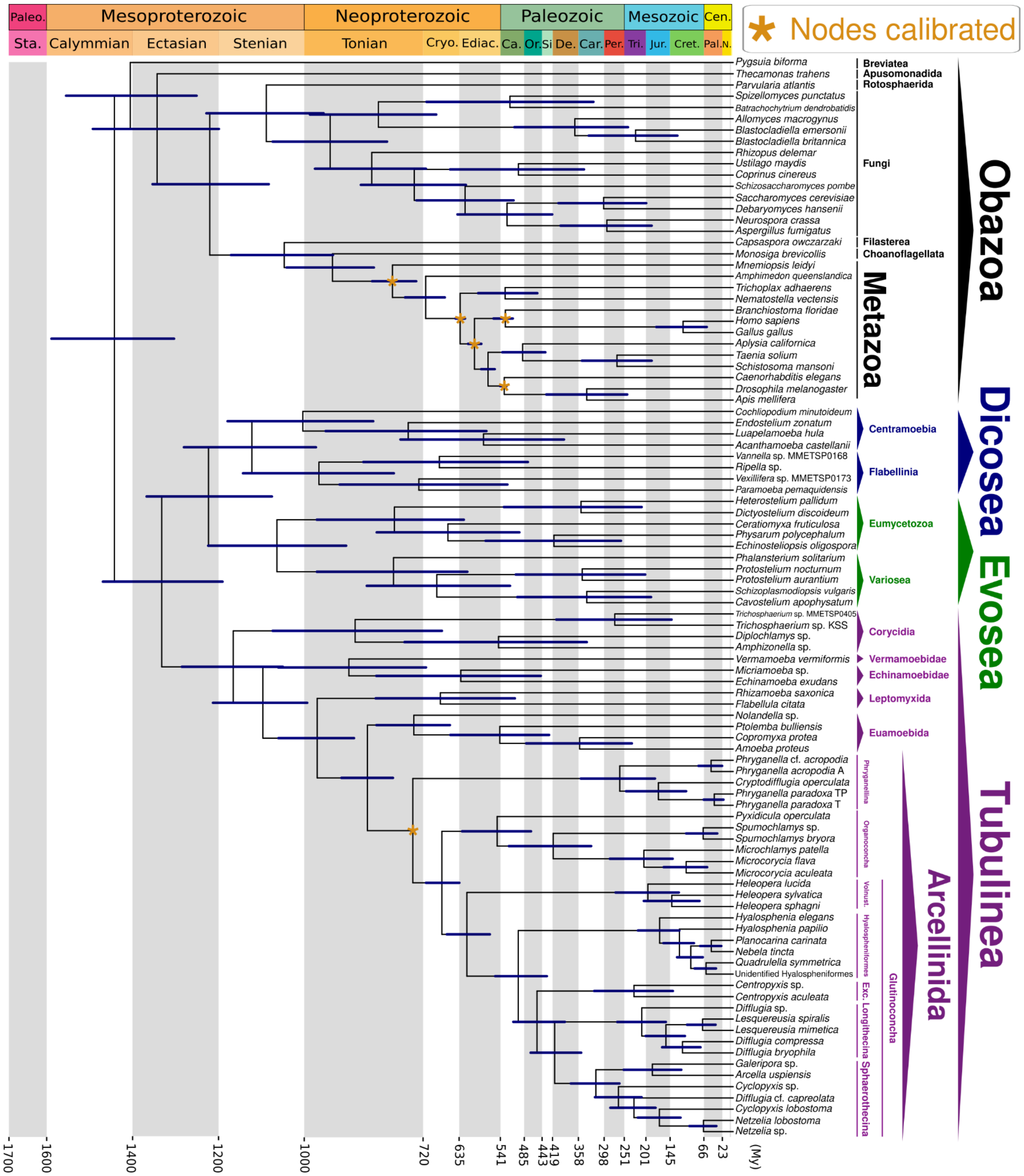


**Figure S28. Amorphea time-calibrated tree inferred under an uncorrelated relaxed clock model, applying a uniform distribution for node calibration and drift parameter of $\alpha$ = 1 and β = 10, calibrating nodes within the Metazoa group, as well as the Arcellinida node (Dataset S01, Table S11).** Bars at nodes are 95% highest probability density confidence intervals. Abbreviations: Exc.- Excentrotoma; Vonust. - Volnustoma; Paleo. - Paleoproterozoic; Sta. - Statherian; Cryo. - Cryogenian; Ediac. - Ediacaran; Ca. - Cambrian; Or. Ordovician; Si - Silurian; De. - Devonian; Car. Carboniferous; Per. - Permian; Tri. - Triassic; Jur. - Jurassic; Cret. - Cretaceous; Cen. - Cenozoic; Pal. - Paleogene; N. - Neogene; My - Million Years.

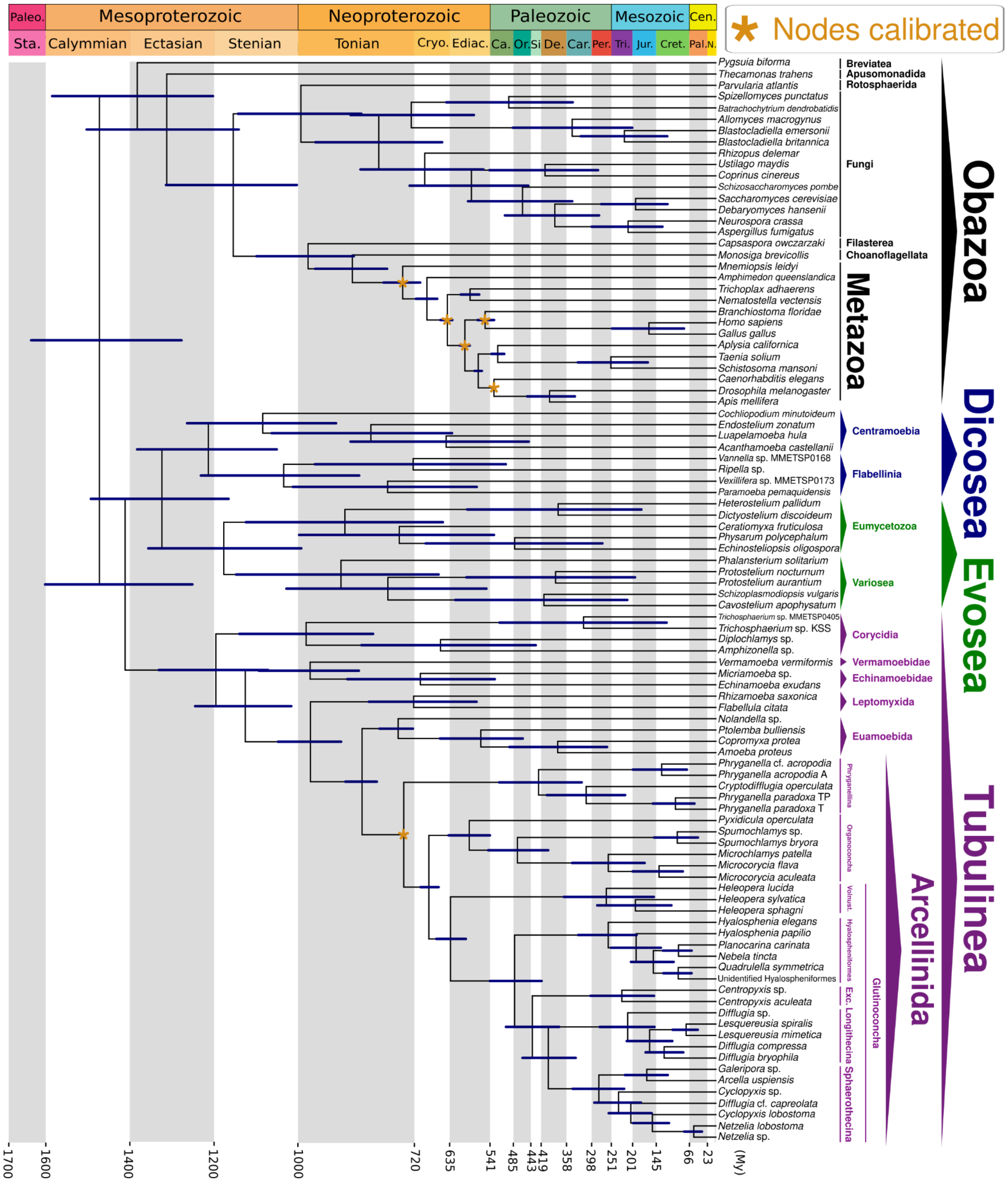


**Figure S29. Amorphea time-calibrated tree inferred under an autocorrelated relaxed clock model, applying a uniform distribution for node calibration and drift parameter of $\alpha$ = 2 and β = 2, calibrating nodes within the Metazoa group, as well as the Arcellinida node (Dataset S01, Table S11).** Bars at nodes are 95% highest probability density confidence intervals. Abbreviations: Exc.- Excentrotoma; Vonust. - Volnustoma; Paleo. - Paleoproterozoic; Sta. - Statherian; Cryo. - Cryogenian; Ediac. - Ediacaran; Ca. - Cambrian; Or. Ordovician; Si - Silurian; De. - Devonian; Car. Carboniferous; Per. - Permian; Tri. - Triassic; Jur. - Jurassic; Cret. - Cretaceous; Cen. - Cenozoic; Pal. - Paleogene; N. - Neogene; My - Million Years.

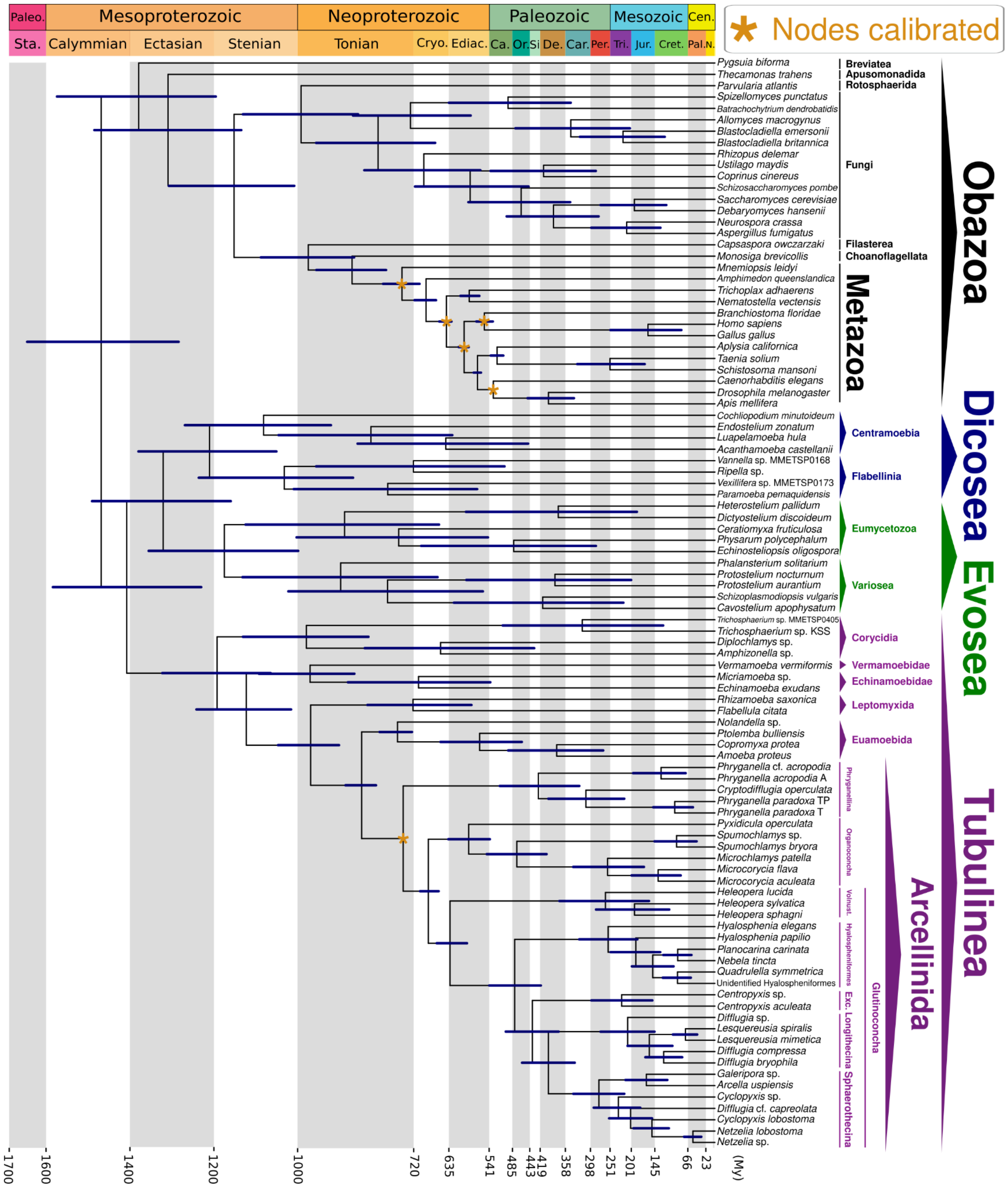


**Figure S30. Amorphea time-calibrated tree inferred under an autocorrelated relaxed clock model, applying a uniform distribution for node calibration and drift parameter of $\alpha$ = 1 and β = 10, calibrating nodes within the Metazoa group, as well as the Arcellinida node (Dataset S01, Table S11).** Bars at nodes are 95% highest probability density confidence intervals. Abbreviations: Exc.- Excentrotoma; Vonust. - Volnustoma; Paleo. - Paleoproterozoic; Sta. - Statherian; Cryo. - Cryogenian; Ediac. - Ediacaran; Ca. - Cambrian; Or. Ordovician; Si - Silurian; De. - Devonian; Car. Carboniferous; Per. - Permian; Tri. - Triassic; Jur. - Jurassic; Cret. - Cretaceous; Cen. - Cenozoic; Pal. - Paleogene; N. - Neogene; My - Million Years.

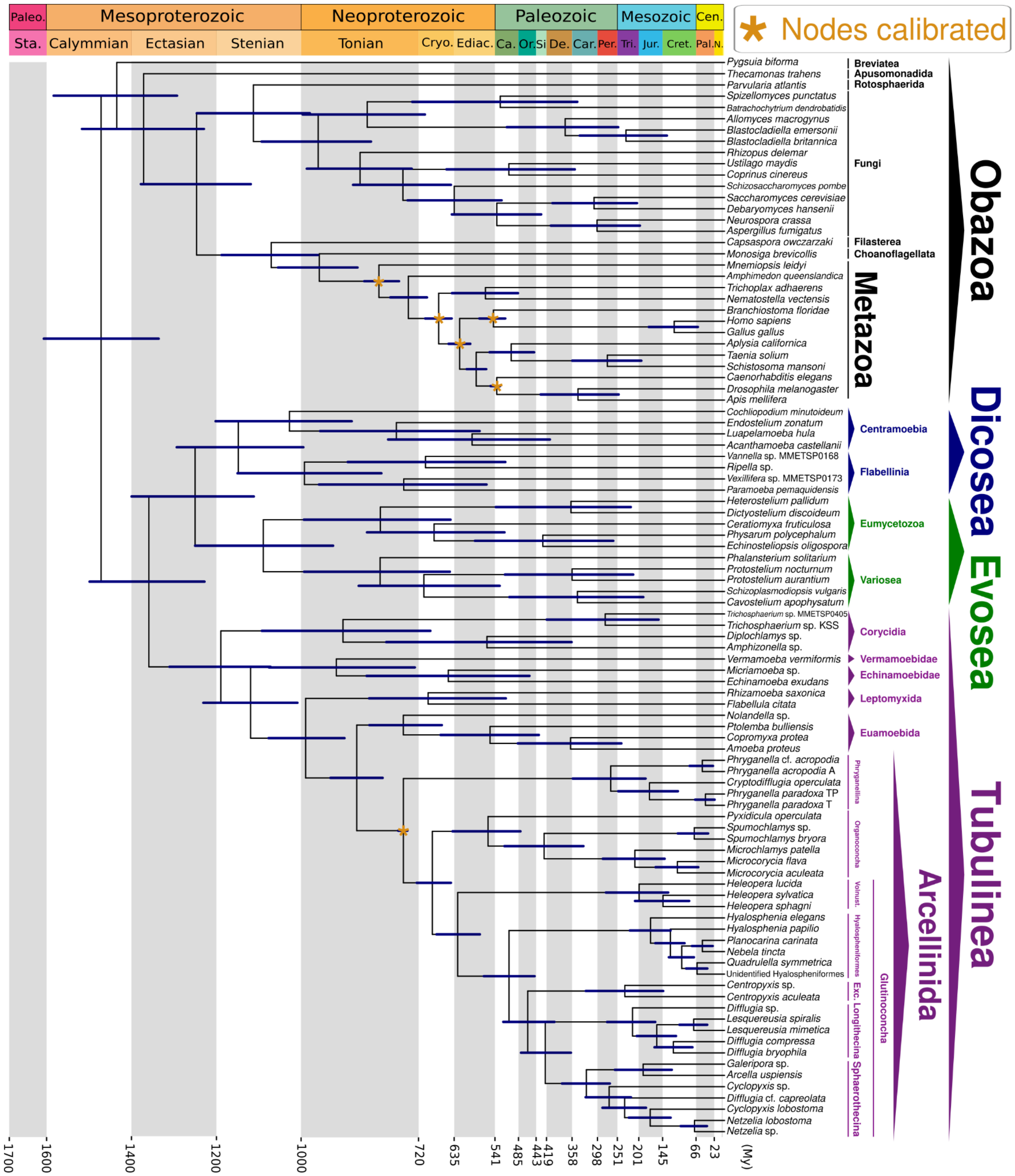

**Figure S31. Amorphea time-calibrated tree inferred under an uncorrelated relaxed clock model, applying a skew-normal distribution for node calibration and drift parameter of $\alpha$ = 2 and β = 2, calibrating nodes within the Metazoa group, as well as the Arcellinida node (Dataset S01, Table S11).** Bars at nodes are 95% highest probability density confidence intervals. Abbreviations: Exc.- Excentrotoma; Vonust. - Volnustoma; Paleo. - Paleoproterozoic; Sta. - Statherian; Cryo. - Cryogenian; Ediac. - Ediacaran; Ca. - Cambrian; Or. Ordovician; Si - Silurian; De. - Devonian; Car. Carboniferous; Per. - Permian; Tri. - Triassic; Jur. - Jurassic; Cret. - Cretaceous; Cen. - Cenozoic; Pal. - Paleogene; N. - Neogene; My - Million Years.

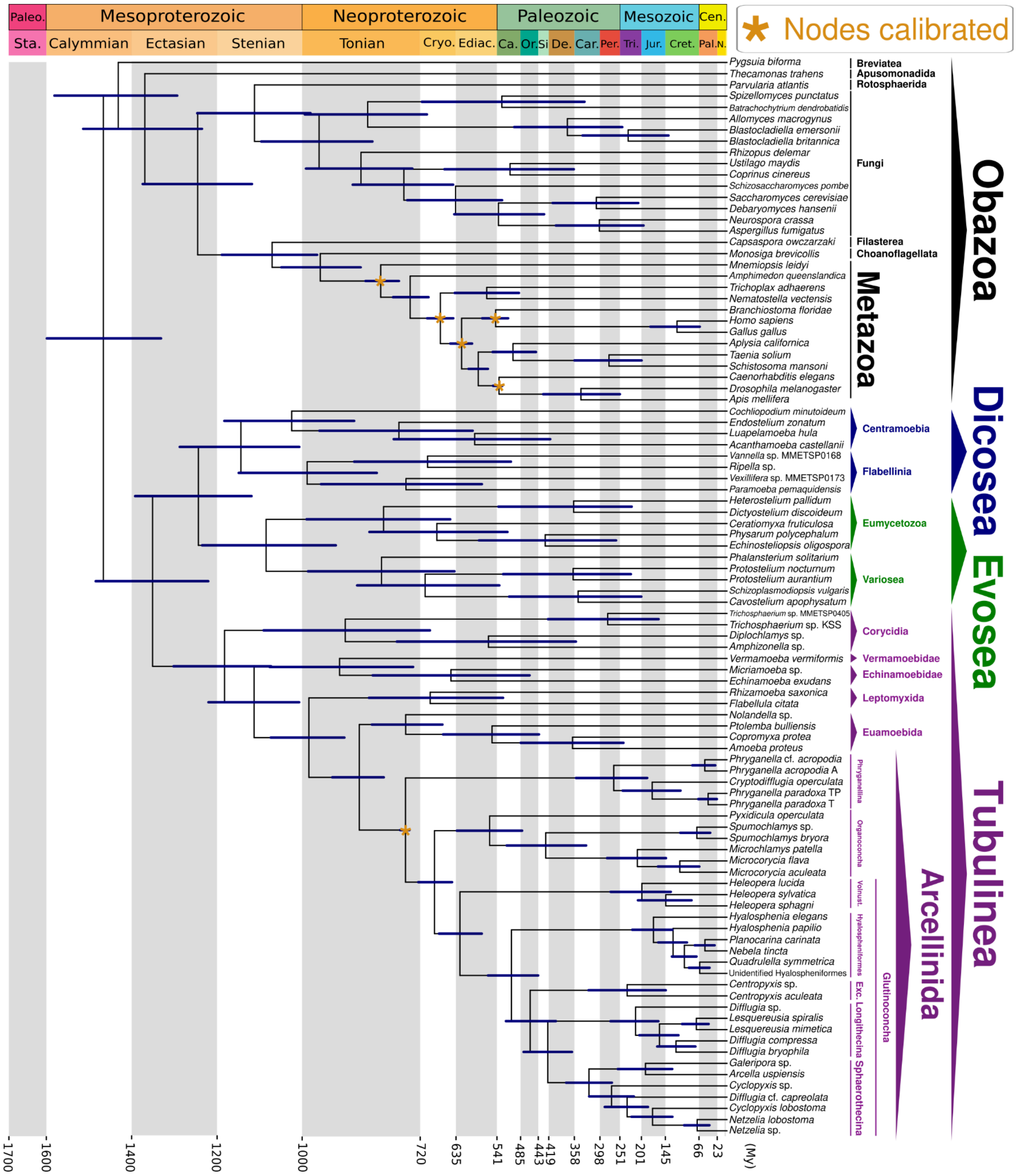



**Figure S33. Amorphea time-calibrated tree inferred under an uncorrelated relaxed clock model, applying a skew-normal distribution for node calibration and drift parameter of $\alpha$ = 1 and β = 10, calibrating nodes within the Metazoa group, as well as the Arcellinida node (Dataset S01, Table S11).** Bars at nodes are 95% highest probability density confidence intervals. Abbreviations: Exc.- Excentrotoma; Vonust. - Volnustoma; Paleo. - Paleoproterozoic; Sta. - Statherian; Cryo. - Cryogenian; Ediac. - Ediacaran; Ca. - Cambrian; Or. Ordovician; Si - Silurian; De. - Devonian; Car. Carboniferous; Per. - Permian; Tri. - Triassic; Jur. - Jurassic; Cret. - Cretaceous; Cen. - Cenozoic; Pal. - Paleogene; N. - Neogene; My - Million Years.

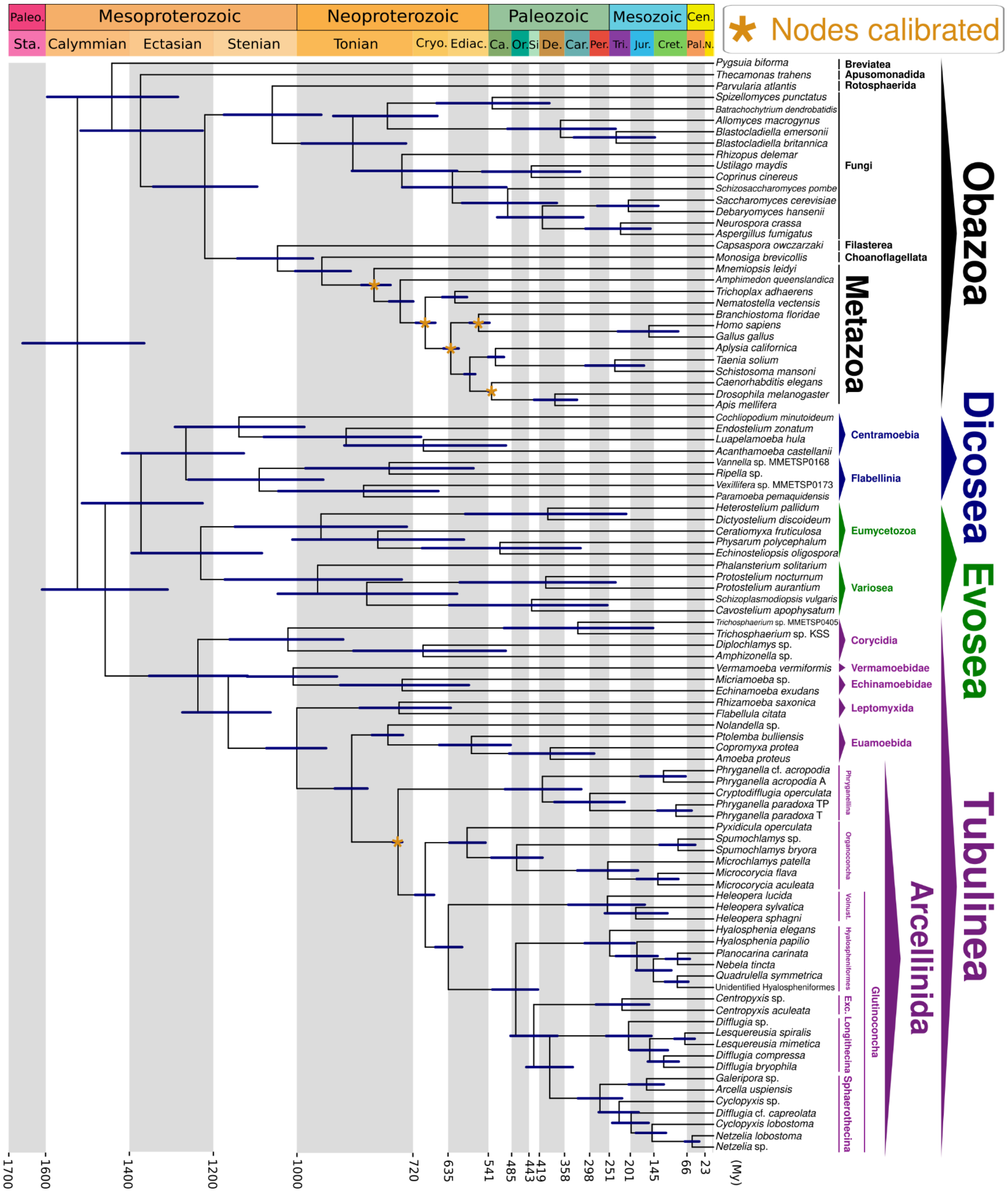


**Figure S34. Amorphea time-calibrated tree inferred under an autocorrelated relaxed clock model, applying a skew-normal distribution for node calibration and drift parameter of $\alpha$ = 2 and β = 2, calibrating nodes within the Metazoa group, as well as the Arcellinida node (Dataset S01, Table S11).** Bars at nodes are 95% highest probability density confidence intervals. Abbreviations: Exc.- Excentrotoma; Vonust. - Volnustoma; Paleo. - Paleoproterozoic; Sta. - Statherian; Cryo. - Cryogenian; Ediac. - Ediacaran; Ca. - Cambrian; Or. Ordovician; Si - Silurian; De. - Devonian; Car. Carboniferous; Per. - Permian; Tri. - Triassic; Jur. - Jurassic; Cret. - Cretaceous; Cen. - Cenozoic; Pal. - Paleogene; N. - Neogene; My - Million Years.

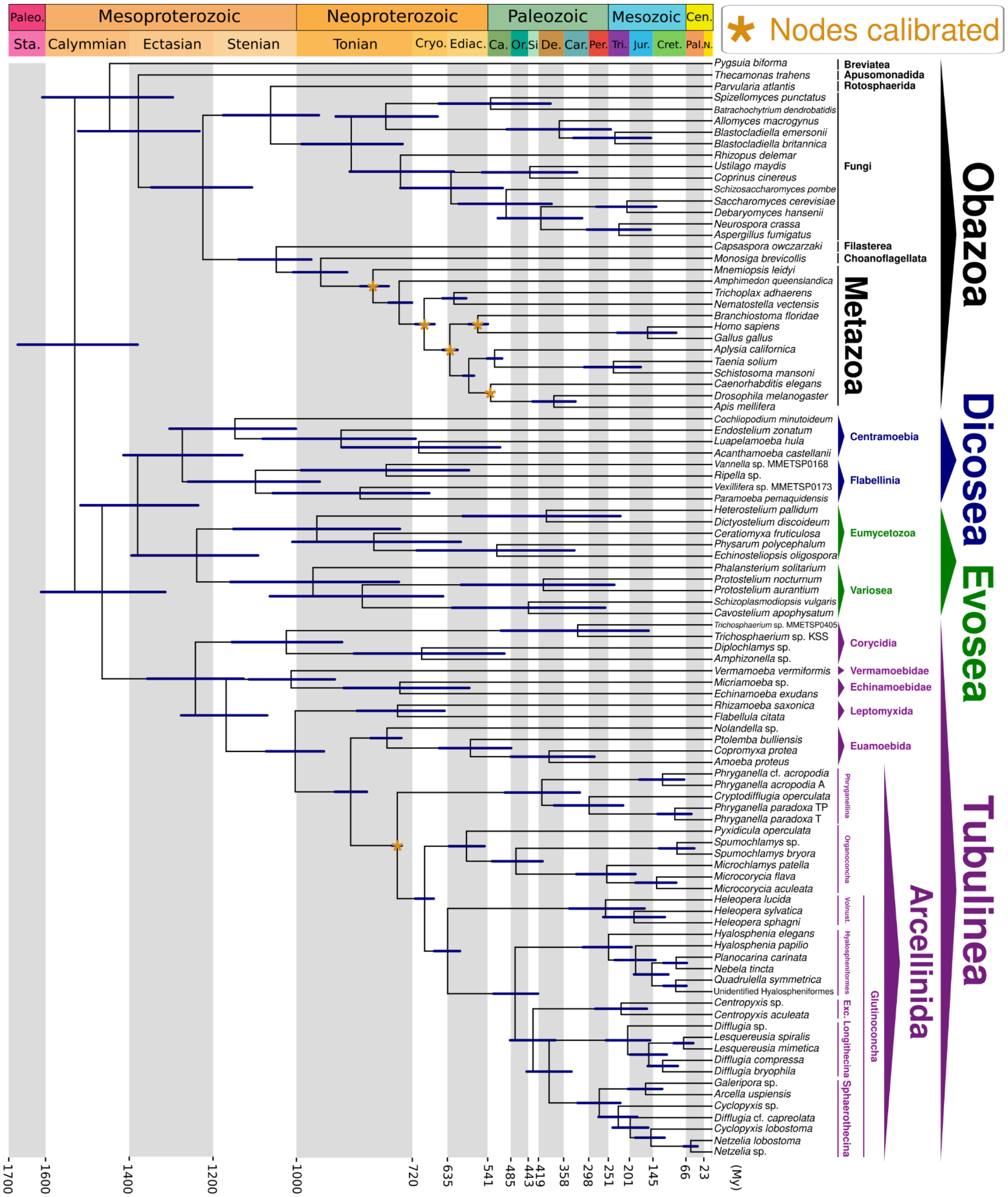


**Figure S35. Amorphea time-calibrated tree inferred under an autocorrelated relaxed clock model, applying a skew-normal distribution for node calibration and drift parameter of $\alpha$ = 1 and β = 10, calibrating nodes within the Metazoa group, as well as the Arcellinida node (Dataset S01, Table S11).** Bars at nodes are 95% highest probability density confidence intervals. Abbreviations: Exc.- Excentrotoma; Vonust. - Volnustoma; Paleo. - Paleoproterozoic; Sta. - Statherian; Cryo. - Cryogenian; Ediac. - Ediacaran; Ca. - Cambrian; Or. Ordovician; Si - Silurian; De. - Devonian; Car. Carboniferous; Per. - Permian; Tri. - Triassic; Jur. - Jurassic; Cret. - Cretaceous; Cen. - Cenozoic; Pal. - Paleogene; N. - Neogene; My - Million Years.

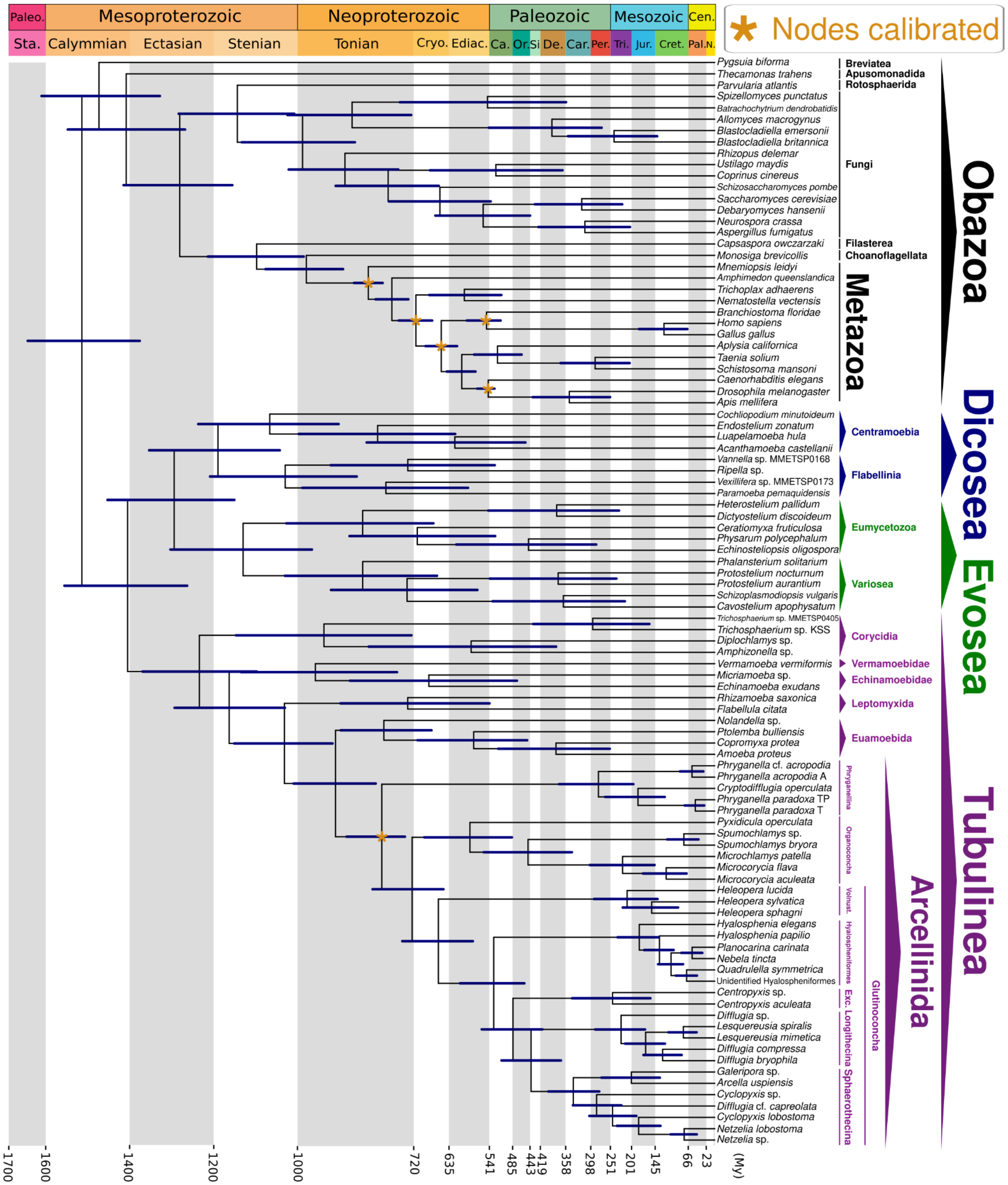


**Figure S36. Amorphea time-calibrated tree inferred under an uncorrelated relaxed clock model, applying a truncated-cauchy short-tail distribution for node calibration and drift parameter of $\alpha$ = 2 and β = 2, calibrating nodes within the Metazoa group, as well as the Arcellinida node (Dataset S01, Table S11).** Bars at nodes are 95% highest probability density confidence intervals. Abbreviations: Exc.- Excentrotoma; Vonust. - Volnustoma; Paleo. - Paleoproterozoic; Sta. - Statherian; Cryo. - Cryogenian; Ediac. - Ediacaran; Ca. - Cambrian; Or. Ordovician; Si - Silurian; De. - Devonian; Car. Carboniferous; Per. - Permian; Tri. - Triassic; Jur. - Jurassic; Cret. - Cretaceous; Cen. - Cenozoic; Pal. - Paleogene; N. - Neogene; My - Million Years.

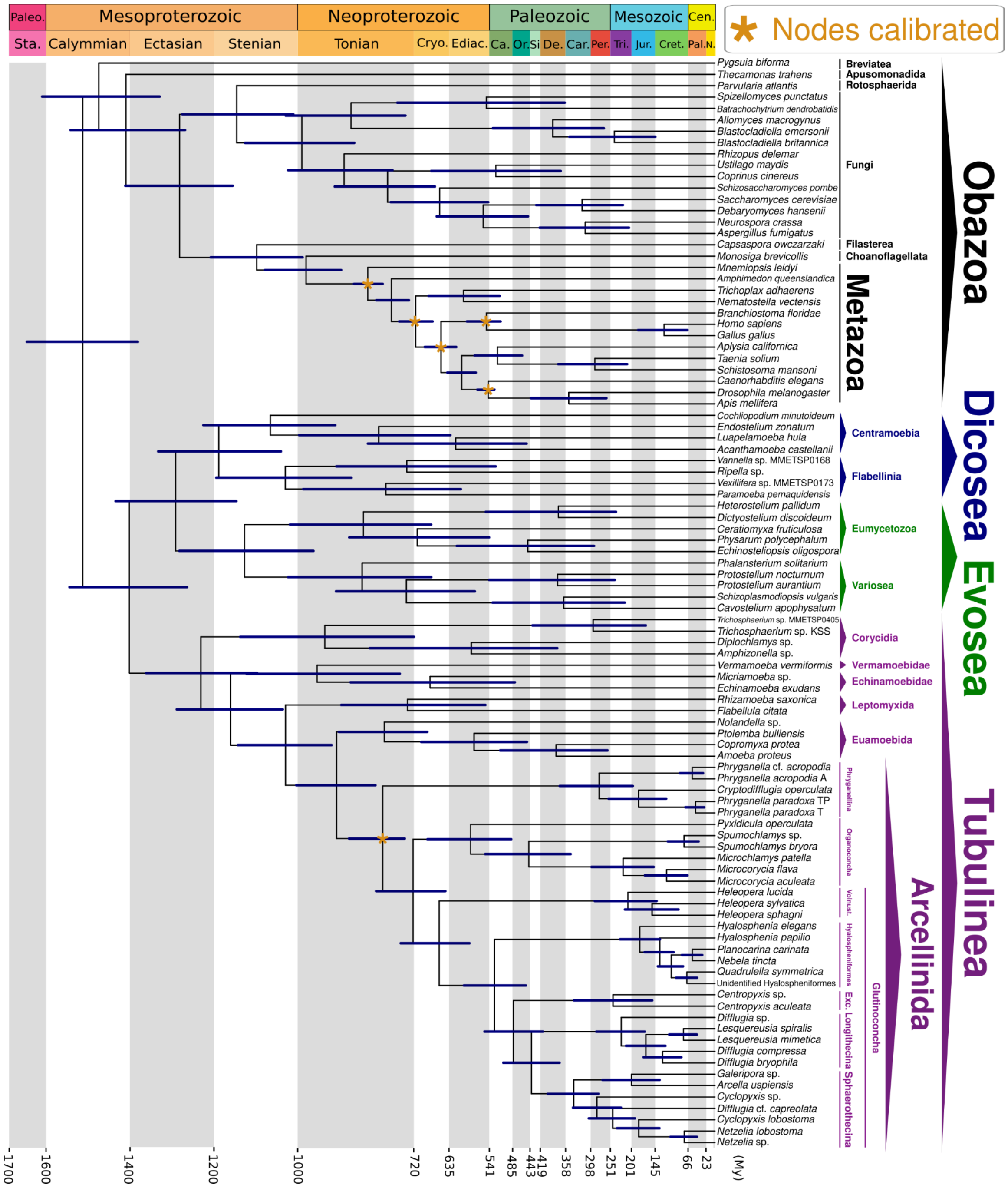


**Figure S37. Amorphea time-calibrated tree inferred under an uncorrelated relaxed clock model, applying a truncated-cauchy short-tail distribution for node calibration and drift parameter of $\alpha$ = 1 and β = 10, calibrating nodes within the Metazoa group, as well as the Arcellinida node (Dataset S01, Table S11).** Bars at nodes are 95% highest probability density confidence intervals. Abbreviations: Exc.- Excentrotoma; Vonust. - Volnustoma; Paleo. - Paleoproterozoic; Sta. - Statherian; Cryo. - Cryogenian; Ediac. - Ediacaran; Ca. - Cambrian; Or. Ordovician; Si - Silurian; De. - Devonian; Car. Carboniferous; Per. - Permian; Tri. - Triassic; Jur. - Jurassic; Cret. - Cretaceous; Cen. - Cenozoic; Pal. - Paleogene; N. - Neogene; My - Million Years.

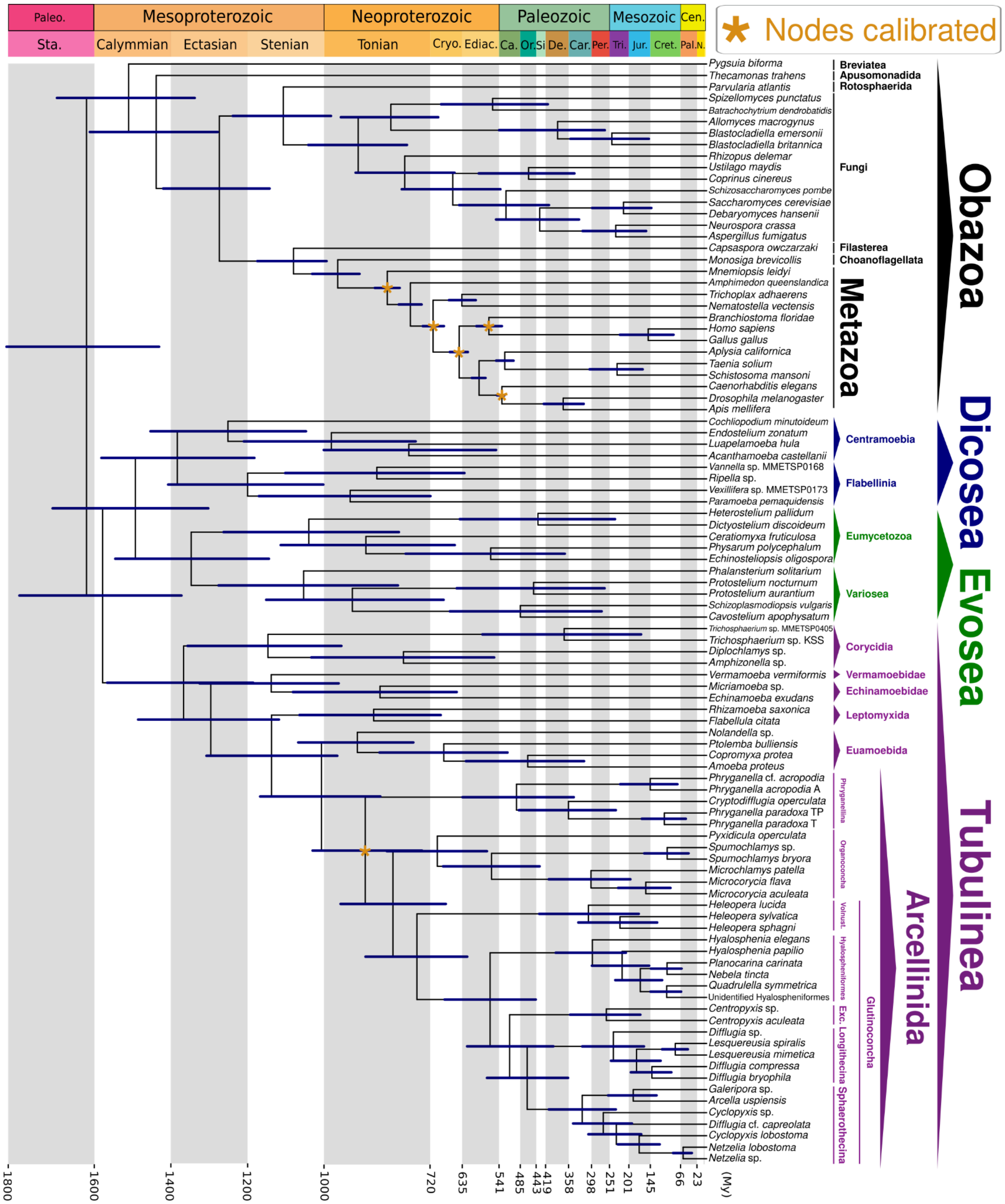


**Figure S38. Amorphea time-calibrated tree inferred under an autocorrelated relaxed clock model, applying a truncated-cauchy short-tail distribution for node calibration and drift parameter of $\alpha$ = 2 and β = 2, calibrating nodes within the Metazoa group, as well as the Arcellinida node (Dataset S01, Table S11).** Bars at nodes are 95% highest probability density confidence intervals. Abbreviations: Exc.- Excentrotoma; Vonust. - Volnustoma; Paleo. - Paleoproterozoic; Sta. - Statherian; Cryo. - Cryogenian; Ediac. - Ediacaran; Ca. - Cambrian; Or. Ordovician; Si - Silurian; De. - Devonian; Car. Carboniferous; Per. - Permian; Tri. - Triassic; Jur. - Jurassic; Cret. - Cretaceous; Cen. - Cenozoic; Pal. - Paleogene; N. - Neogene; My - Million Years.

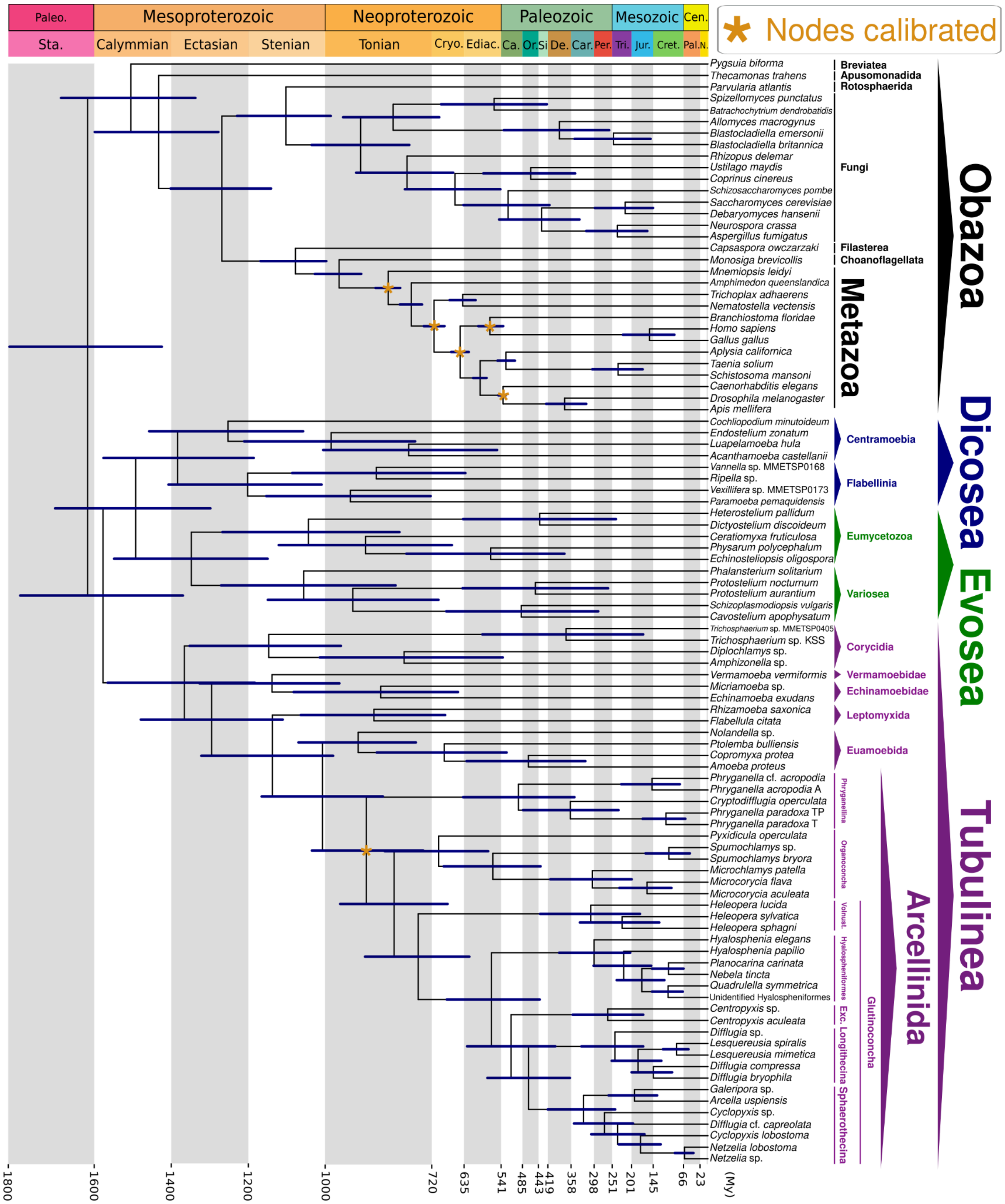

**Figure S39. Amorphea time-calibrated tree inferred under an autocorrelated relaxed clock model, applying a truncated-cauchy short-tail distribution for node calibration and drift parameter of $\alpha$ = 1 and β = 10, calibrating nodes within the Metazoa group, as well as the Arcellinida node (Dataset S01, Table S11).** Bars at nodes are 95% highest probability density confidence intervals. Abbreviations: Exc.- Excentrotoma; Vonust. - Volnustoma; Paleo. - Paleoproterozoic; Sta. - Statherian; Cryo. - Cryogenian; Ediac. - Ediacaran; Ca. - Cambrian; Or. Ordovician; Si - Silurian; De. - Devonian; Car. Carboniferous; Per. - Permian; Tri. - Triassic; Jur. - Jurassic; Cret. - Cretaceous; Cen. - Cenozoic; Pal. - Paleogene; N. - Neogene; My - Million Years.

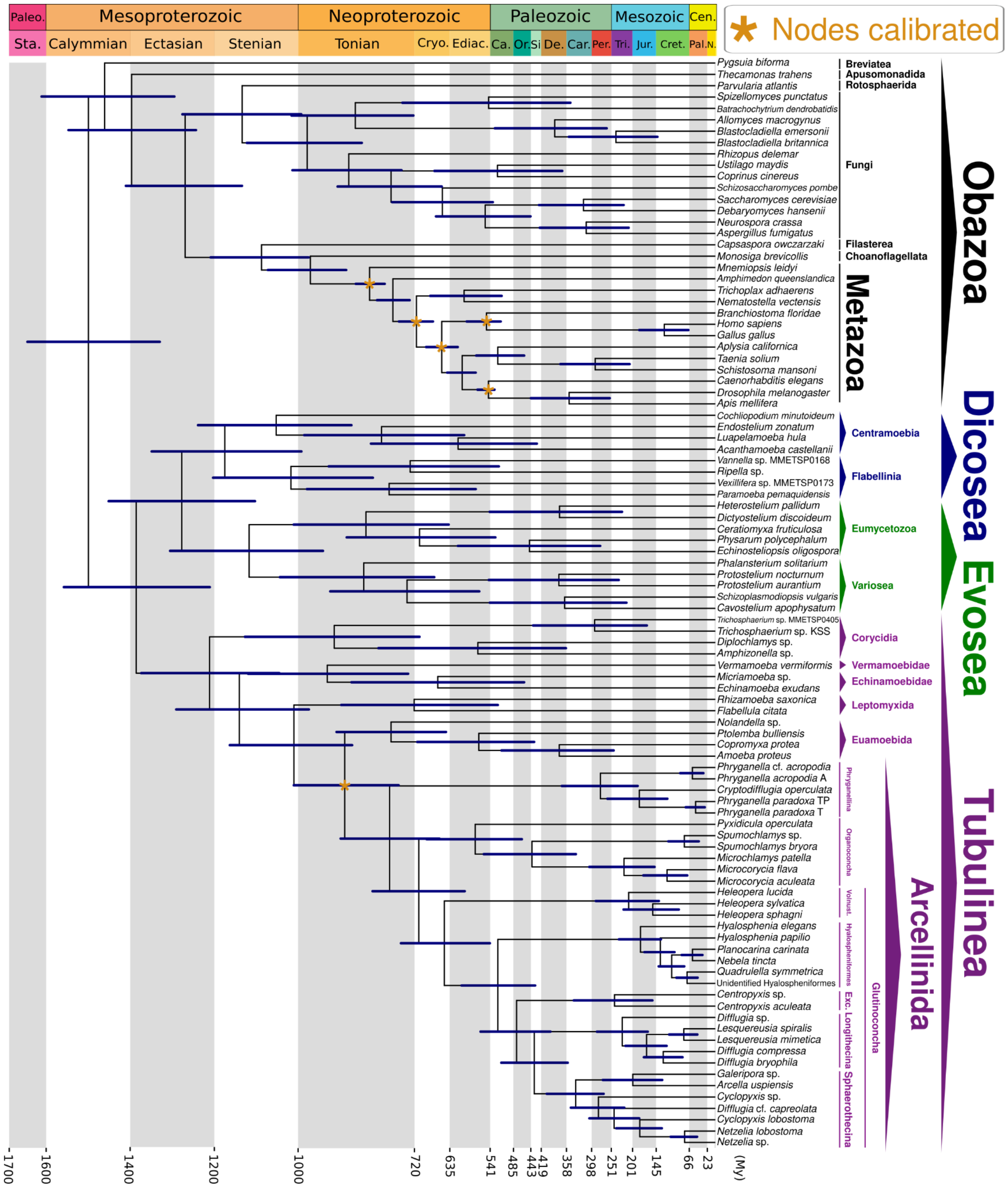


**Figure S40. Amorphea time-calibrated tree inferred under an uncorrelated relaxed clock model, applying a truncated-cauchy short-tail distribution for node calibration and drift parameter of $\alpha$ = 2 and β = 2, calibrating nodes within the Metazoa group, as well as the Euamoebida+Arcellinida node (Dataset S01, Table S11).** Bars at nodes are 95% highest probability density confidence intervals. Abbreviations: Exc.- Excentrotoma; Vonust. - Volnustoma; Paleo. - Paleoproterozoic; Sta. - Statherian; Cryo. - Cryogenian; Ediac. - Ediacaran; Ca. - Cambrian; Or. Ordovician; Si - Silurian; De. - Devonian; Car. Carboniferous; Per. - Permian; Tri. - Triassic; Jur. - Jurassic; Cret. - Cretaceous; Cen. - Cenozoic; Pal. - Paleogene; N. - Neogene; My - Million Years.

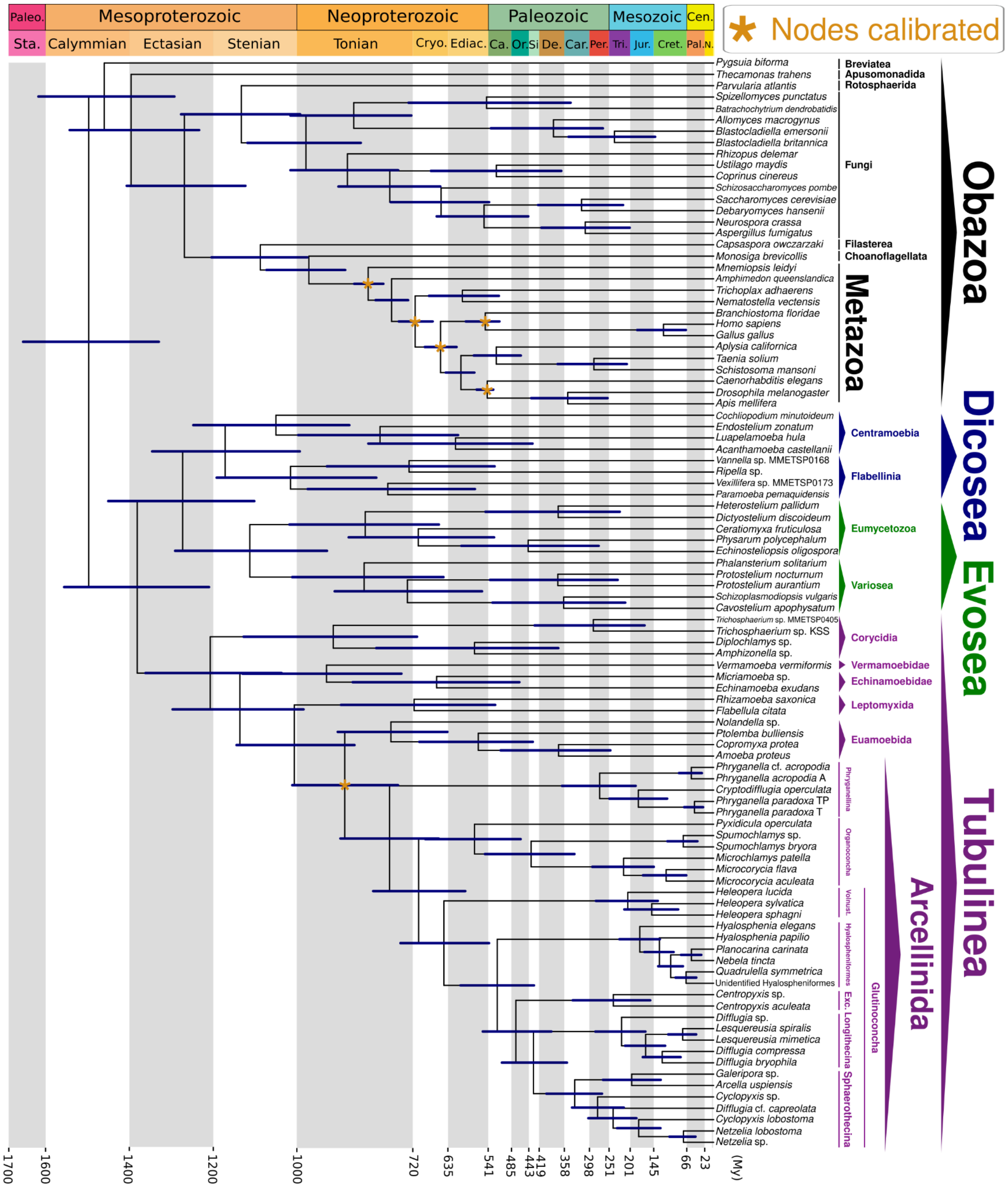


**Figure S41. Amorphea time-calibrated tree inferred under an uncorrelated relaxed clock model, applying a truncated-cauchy short-tail distribution for node calibration and drift parameter of $\alpha$ = 1 and β = 10, calibrating nodes within the Metazoa group, as well as the Euamoebida+Arcellinida node (Dataset S01, Table S11).** Bars at nodes are 95% highest probability density confidence intervals. Abbreviations: Exc.- Excentrotoma; Vonust. - Volnustoma; Paleo. - Paleoproterozoic; Sta. - Statherian; Cryo. - Cryogenian; Ediac. - Ediacaran; Ca. - Cambrian; Or. Ordovician; Si - Silurian; De. - Devonian; Car. Carboniferous; Per. - Permian; Tri. - Triassic; Jur. - Jurassic; Cret. - Cretaceous; Cen. - Cenozoic; Pal. - Paleogene; N. - Neogene; My - Million Years.

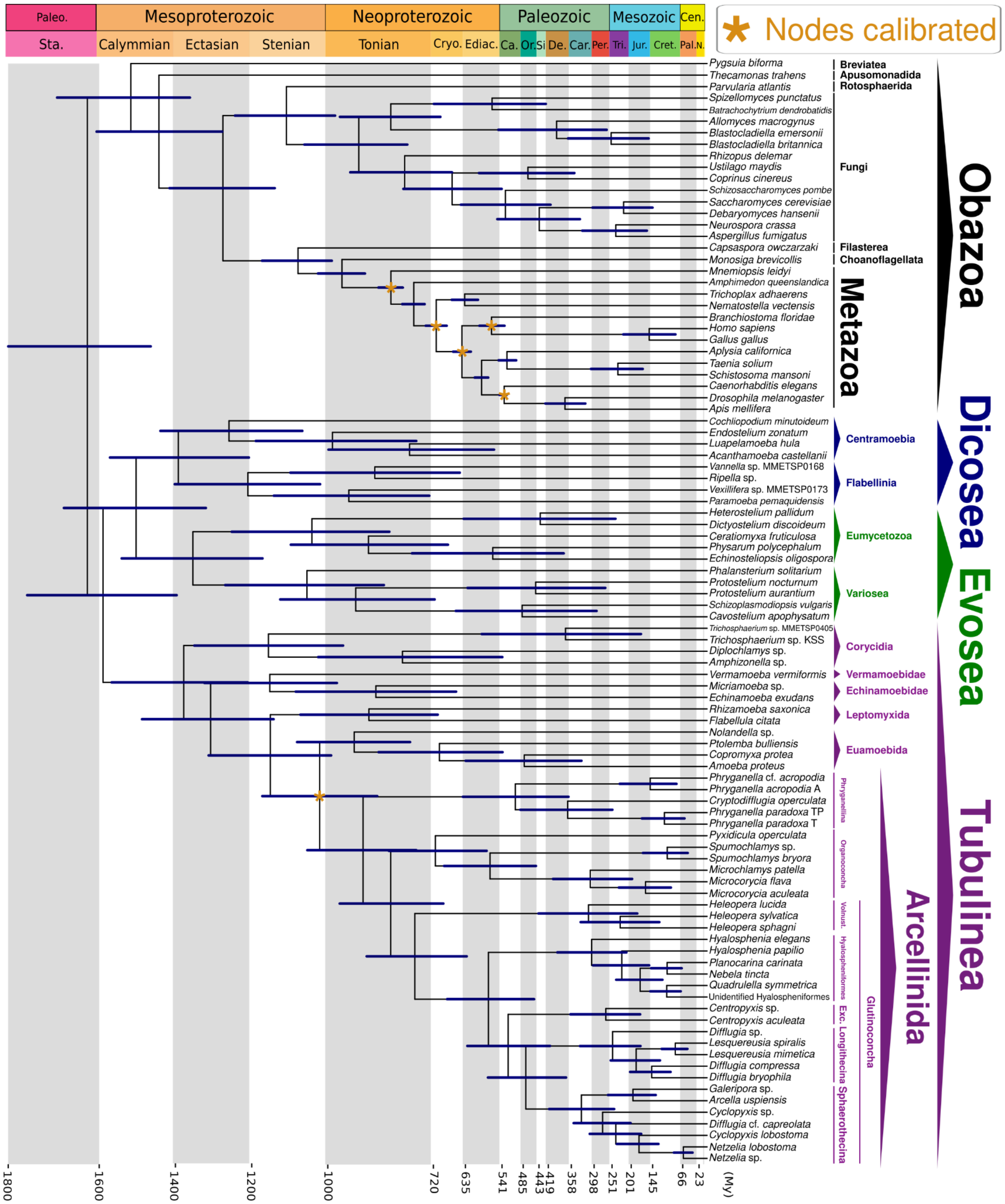

**Figure S42. Amorphea time-calibrated tree inferred under an autocorrelated relaxed clock model, applying a truncated-cauchy short-tail distribution for node calibration and drift parameter of $\alpha$ = 2 and β = 2, calibrating nodes within the Metazoa group, as well as the Euamoebida+Arcellinida node (Dataset S01, Table S11).** Bars at nodes are 95% highest probability density confidence intervals. Abbreviations: Exc.- Excentrotoma; Vonust. - Volnustoma; Paleo. - Paleoproterozoic; Sta. - Statherian; Cryo. - Cryogenian; Ediac. - Ediacaran; Ca. - Cambrian; Or. Ordovician; Si - Silurian; De. - Devonian; Car. Carboniferous; Per. - Permian; Tri. - Triassic; Jur. - Jurassic; Cret. - Cretaceous; Cen. - Cenozoic; Pal. - Paleogene; N. - Neogene; My - Million Years.

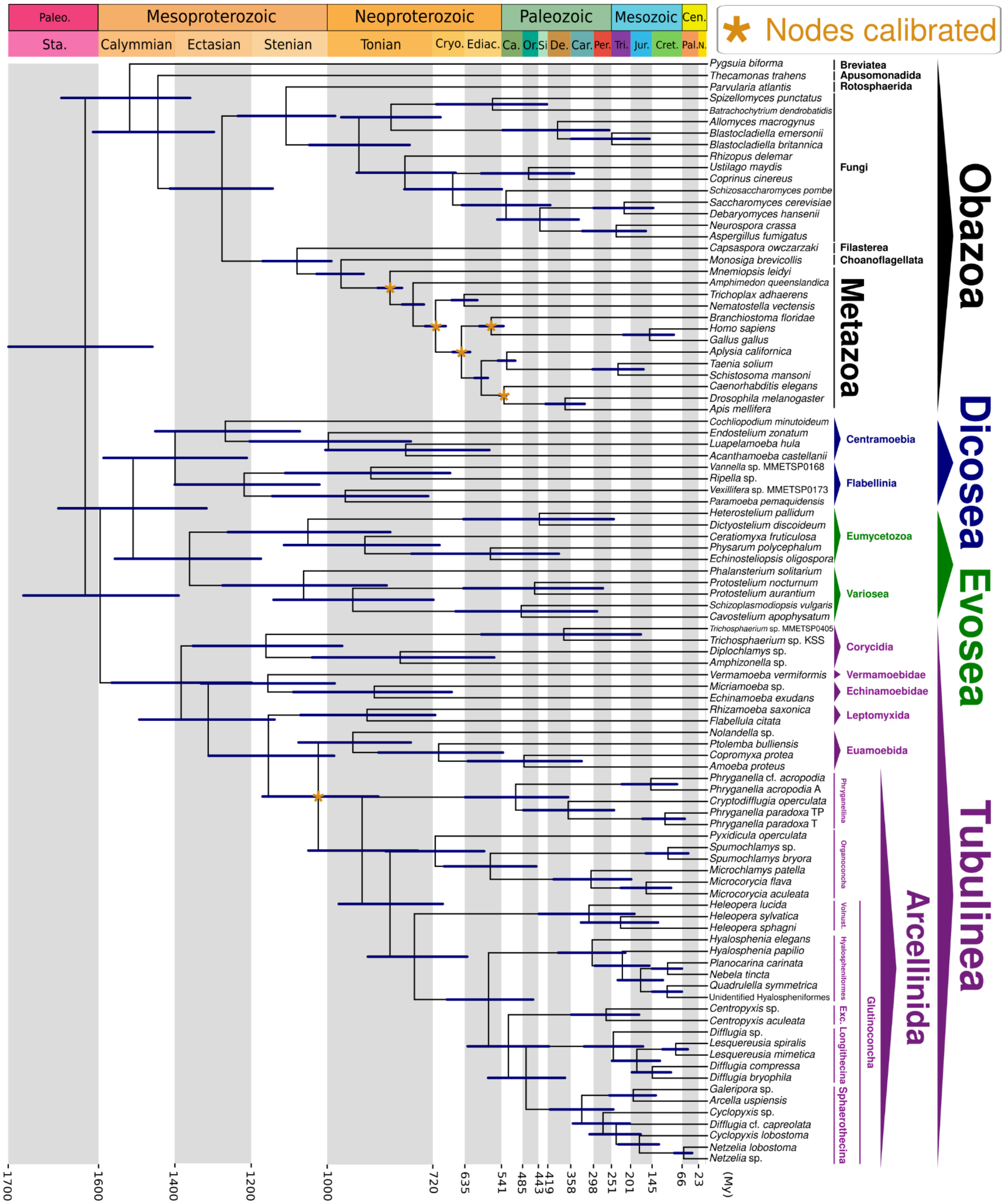


**Figure S43. Amorphea time-calibrated tree inferred under an autocorrelated relaxed clock model, applying a truncated-cauchy short-tail distribution for node calibration and drift parameter of $\alpha$ = 1 and β = 10, calibrating nodes within the Metazoa group, as well as the Euamoebida+Arcellinida node (Dataset S01, Table S11).** Bars at nodes are 95% highest probability density confidence intervals. Abbreviations: Exc.- Excentrotoma; Vonust. - Volnustoma; Paleo. - Paleoproterozoic; Sta. - Statherian; Cryo. - Cryogenian; Ediac. - Ediacaran; Ca. - Cambrian; Or. Ordovician; Si - Silurian; De. - Devonian; Car. Carboniferous; Per. - Permian; Tri. - Triassic; Jur. - Jurassic; Cret. - Cretaceous; Cen. - Cenozoic; Pal. - Paleogene; N. - Neogene; My - Million Years.

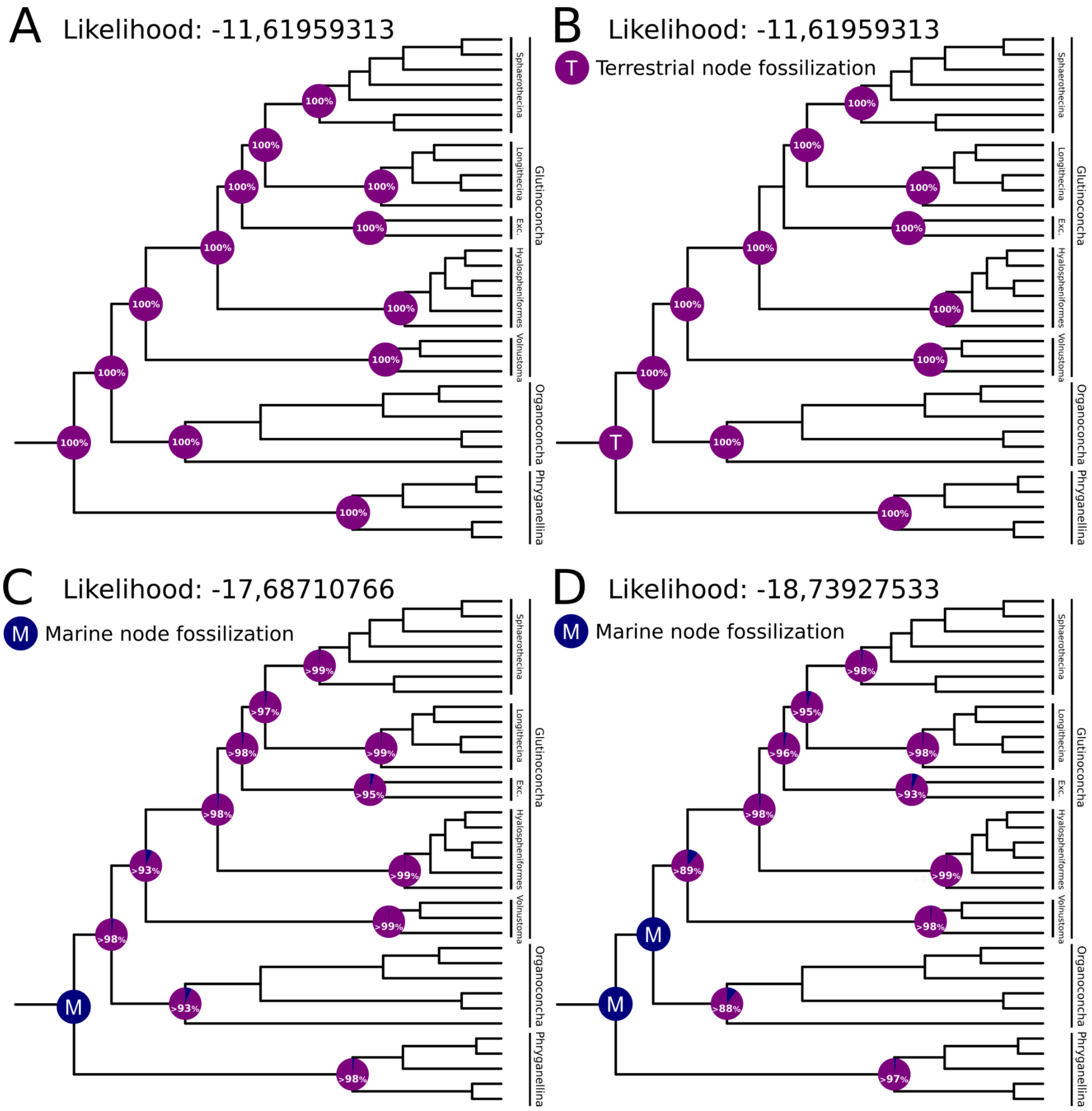


**Figure S44. Ancestral habitat state reconstruction of Arcellinida (DatasetS01, Table S10**). **A.** Ancestral habitat state reconstruction of arcellinids without assigning ancestral state (fossilizing) nodes. **B.** Ancestral habitat state reconstruction of arcellinids fossilizing Arcellinida node as terrestrial, which interprets the organisms represented by VSMs as terrestrial. **C.** Ancestral habitat state reconstruction of arcellinids fossilizing Arcellinida node as marine, which interprets the organisms represented by VSMs as marine. **D.** Ancestral habitat state reconstruction of arcellinids fossilizing Arcellinida and Organoconcha+Glutinoconcha nodes as marine, which interprets the organisms represented by VSMs as marine. Based on the likelihood ratio test (LRT), which we considered as significant a difference of LRT ≥ 2 (41), the scenarios presented in **A** and **B** are statistically superior to the other scenarios (**Dataset S01, Table S12**). Purple circles represent nodes fossilized as terrestrial (T) ancestral habitat state and blue circles represent nodes fossilized as marine (M) ancestral habitat state. Likelihood values represent the mean calculated likelihood for each scenario (**Dataset S01, Table S12**).

**Legends of supplemental Tables (Dataset S01)**

**Table S1.** Taxa information and source of the data used for the phylogenomic and molecular clock analyses. **a.** Novel 'Omic Data. **b.** Previously Publicly Available 'Omic Data.

**TableS2.** Measured morphometric characteristics of the newly sampled testate amoebae taxa.

**Table S3.** Phylogenomic and Molecular Clock matrices data information. **a.** Phylogenomic Matrix (Fig. 2) Taxon Composition and Completeness Matrice. **b.** Phylogenomic Matrix (Fig. 2) Gene Composition and Indices for Concatenated Matrix. **c.** Phylogenomic Matrix (Fig. 2) Gene Occupancy Data for Each Taxon.

**Table S4.** Amoebozoan phylogenomic supermatrix.

**Table S5.** Small Subunit ribosomal RNA (SSU) dataset, sequences ID and NCBI accession number.

**Table S6.** Cytochrome C oxidase subunit I (COI), sequences ID and NCBI accession number.

**Table S7.** Amorphea phylogenomic supermatrix – Input for MCMCTree.

**TableS8.** Calibrations used for dating the Amorphea tree. Arcellinida ages based on the Arcellinida vase-shaped microfossil record (Table S9 and 20).

**Table S9.** Arcellinida vase-shaped microfossil record showing formation ages in million years (MA), dating methods, and references.

**Table S10.** Calibration strategies and MCMCTree input files. The nodes numbering is shown on Appendix S01, Fig. S6. **a.** Node calibration ages. **b.** Topology with node calibration mapped. **c.** Control file for clock model 2 or 3 and a drift parameter of $\alpha$ = 2 and $\beta$ = 2 or of $\alpha$ = 1 and $\beta$ = 10.

**Table S11.** MCMCTree results. The nodes numbering is shown on Appendix S01, Fig. S6. **a.** Complete results considering three calibration strategies: i. calibration of nodes within the animals' group, not considering VSM record to calibrate amoebozoan nodes (No VSM); ii. calibration of nodes within the animals' group and calibration of Glutinoconcha+Organoconcha and Glutinoconcha nodes, considering VSMs as derived crown arcellinids (VSM derived Crown arcellinid); iii. calibration of nodes within the animals' group and calibration of the Arcellinida node, considering VSMs as basal crown arcellinids (VSM basal Crown arcellinid); iv. calibration of Arcellinida+Euamoebida node, considering VSMs as stem arcellinids (VSM stem arcellinid). **b.** Summary of estimated means, medians, and likelihoods of each experiment considering chain 1 based on the complete results shown on Table S11a. **c.** Summary of the complete results (a.) focusing on the key nodes discussed in the main text of the manuscript.

**Table S12.** BayesTraits input files and ancestral habitat (marine (M) or terrestrial (T)) reconstruction. The results presented and discussed on Figure 5 and in the main text represent the calculated mean considering the likelihood and probability values calculated for each of the 100 input trees. P(M) indicates the probability of a marine ancestor and P(T) indicates the probability of a terrestrial ancestor. **a.** 100 trees with branch lengths used as input to BayesTraits. **b.** Data file input to BaysTraits. **c.** BayesTrait command file used to reconstruct the ancestral habitat states of arcellinids without assigning ancestral state (fossilizing) nodes. **d.** BaysTraits results following the command file shown in TableS12c to reconstruct ancestral habitat states without assigning (fossilizing) ancestral state to any node. **e.** BayesTrait command file used to reconstruct the ancestral habitat states of arcellinids fossilizing Arcellinida node as terrestrial, which interprets the organisms represented by VSMs as terrestrial. **f.** BaysTraits results following the command file shown in TableS12e to reconstruct ancestral habitat states fossilizing Arcellinida node as terrestrial. **g.** BayesTrait command file used to reconstruct the ancestral habitat states of arcellinids fossilizing Arcellinida node as marine, which interprets the organisms represented by VSMs as marine. **h.** BaysTraits results following the command file shown in TableS12g to reconstruct ancestral habitat states fossilizing Arcellinida node as marine. **i.** BayesTrait command file used to reconstruct the ancestral habitat states of arcellinids fossilizing Arcellinida and Organoconcha+Glutinoconcha nodes as marine, which interprets the organisms represented by VSMs as derived crown Arcellinida that lived in marine habitat. **j.** BaysTraits results following the command file shown in TableS12i to reconstruct ancestral habitat states fossilizing Arcellinida and Organoconcha+Glutinoconcha nodes as marine. **k.** We performed likelihood ratio tests (LRT) to compare the scenario inferred by the ancestral state reconstructions (Fig. S16). LRT= 2[log-likelihood(scenario1) – log-likelihood(alternative scenario)]; we considered as significant a difference of LRT ≥ 2 following Pagel, (41).
